# Characterisation of the SARS-CoV-2 ExoN (nsp14^ExoN^-nsp10) complex: implications for its role in viral genome stability and inhibitor identification

**DOI:** 10.1101/2020.08.13.248211

**Authors:** Hannah T. Baddock, Sanja Brolih, Yuliana Yosaatmadja, Malitha Ratnaweera, Marcin Bielinski, Lonnie P. Swift, Abimael Cruz-Migoni, Garrett M. Morris, Christopher J. Schofield, Opher Gileadi, Peter J. McHugh

**Affiliations:** Department of Oncology, MRC Weatherall Institute of Molecular Medicine, University of Oxford, John Radcliffe Hospital, Oxford OX3 9DS, UK; Structural Genomics Consortium, University of Oxford, Old Road Campus Research Building, Roosevelt Drive, Oxford OX3 7DQ, UK; Chemistry Research Laboratory, University of Oxford, Mansfield Road, Oxford OX1 3TA, UK; Department of Statistics, University of Oxford, 24-29 St Giles′, Oxford, OX1 3LB, UK.

**Author notes:** These authors contributed equally: Hannah T. Baddock, Sanja Brolih and Yuliana Yosaatmadja.

## Abstract

The SARS-CoV-2 coronavirus (CoV) causes COVID-19, a current global pandemic. SARS-CoV-2 belongs to an order of Nidovirales with very large RNA genomes. It is proposed that the fidelity of CoV genome replication is aided by an RNA nuclease complex, formed of non-structural proteins 14 and 10 (nsp14-nsp10), an attractive target for antiviral inhibition. Here, we confirm that the SARS-CoV-2 nsp14-nsp10 complex is an RNase. Detailed functional characterisation reveals nsp14-nsp10 is a highly versatile nuclease capable of digesting a wide variety of RNA structures, including those with a blocked 3’-terminus. We propose that the role of nsp14-nsp10 in maintaining replication fidelity goes beyond classical proofreading and purges the nascent replicating RNA strand of a range of potentially replication terminating aberrations. Using the developed assays, we identify a series of drug and drug-like molecules that potently inhibit nsp14-nsp10, including the known Sars-Cov-2 major protease (M^pro^) inhibitor ebselen and the HIV integrase inhibitor raltegravir, revealing the potential for bifunctional inhibitors in the treatment of COVID-19.

## INTRODUCTION

From late 2019 and throughout 2020 the SARS-CoV-2 virus, which causes the disease COVID-19, has spread across the globe, to date infecting upwards of ten million people and killing over half a million of these (coronavirus.jhu.edu). A detailed understanding of the mechanistic aspects of this virus’ life and infectivity cycle are urgently required as are drugs that are able to curb its replication and virulence.

SARS-CoV-2 is a coronavirus (CoV), of the coronaviridae family in the Nidovirales order. One characteristic of these CoVs is their (relatively) large single-stranded RNA genomes (∼30 kb in the case of SARS-CoV-2)^1^. Perhaps necessarily, CoVs typically have a replication fidelity rate of an order of 10^-6^ to 10^-7^, which is several orders of magnitude more accurate than that of most RNA viruses (typically ∼10^-3^ to 10^-5^)^2^. In order to maintain the fidelity of these genomes during replication, CoVs rely upon a complex of two non-structural proteins, nsp14 (also known as ExoN) and nsp10^2–4^. The importance of this enhanced level of replication fidelity has been demonstrated in studies that disrupt or inactivate the activity of nsp14-nsp10, where reduced virulence and pathogenesis is seen in mouse and cellular models^4–6^. Therefore, targeting nsp14-nsp10 is an attractive therapeutic strategy, either as a standalone option or as an adjuvant to other agents that target other features of the replication cycle of the virus^7^.

The nsp14-nsp10 proteins form a complex where a ribonuclease activity is conferred by the DEDD catalytic motif of nsp14, but where nsp10 plays a key role in conferring full activity^3, 8, 9^. A study of the SARS-CoV-1 complex implied that its ribonuclease activity is dsRNA selective^3^. This complex has been reported to possess 3′-5′ exonuclease activity and, based on its ability to excise terminally mismatched ribonucleotides, one function ascribed to nsp14-nsp10 (by analogy with replicative DNA polymerases) is ′proofreading′ activity^3, 10^, although other key roles in viral replication have been postulated^11^. Consistent with a role in proofreading of maintaining genome stability, nuclease-inactivating mutations in CoV DEDD motifs cause an elevated level of replication errors, impaired replication and in some cases lethal mutagenesis^4, 5, 12^. Moreover, nsp14 interacts with the CoV RNA-dependent RNA polymerase (RdRp), (i.e. the nsp12 subunit of the nsp12-nsp7-nsp8 complex), integrating the nuclease activities of nsp14 with the replicative process, although the molecular details of this interaction remain only partly characterised^10, 11^. The nsp14 protein also contains a functionally distinct (from its nuclease) SAM-dependent (*S*-adenosyl methionine) guanine-N7 methyl transferase (MTase) activity^13–15^. This activity is required in the third step in production of the mature 5′-RNA CoV cap structure (cap-1), methylating a 5′-5′ triphophosphate GpppN generating the cap-0 intermediate.

Here, we describe purification of the SARS-CoV-2 nsp14-nsp10 complex and detailed characterisation of its nuclease activity and substrate profile. We find that the complex is a highly versatile nuclease able to not only potentially able act in a proofreading capacity, but which is also capable of processing a structurally diverse range of RNA molecules, in both ssRNA and dsRNA. Notably, some of the activities observed do not require a free 3′-OH group, implying that nsp14-nsp10 exhibits both exo- and endonuclease like activities. We propose that nsp14-nsp10 could act broadly to remove structures accumulating within the nascent replicating RNA strand that would potentially affect high fidelity extension by the SARS-CoV-2 RNA-dependent RNA polymerase (RdRp)^16^. We also used the assay systems we developed to screen for inhibitors, identifying several drugs and drug-like molecules that might be repurposed to inhibit the nuclease activity of this complex.

## RESULTS

### SARS-CoV-2 nsp14-nsp10 is a dimeric nuclease

Employing codon optimised constructs expressed in *E. coli*, we purified the nsp14-nsp10 complex to near homogeneity (Fig. 1a). The presence of faint protein bands between the nsp14 and nsp10 proteins which were visible by SDS-PAGE were confirmed to be nsp-14 degradation products by tryptic digest MS/MS analysis. The complex eluted from the size-exclusion column with a volume consistent with a heterodimer (Supplementary Fig. 1a), and the identity of the protein was confirmed by intact mass spectrometry (Supplementary Fig. 1b). In addition to the wild-type protein, we purified a control complex bearing substitutions at residues D113 and E115 (nsp14^D113A,E115A^-nsp10) which by comparison with previously evaluated SARS-CoV-1 complex is expected to be catalytically inactive (Fig. 1a)^8^. Moreover, we purified the nsp14 (ExoN) subunit alone to determine its individual activity (bacterial expressed nsp10 rapidly precipitated in solution, presumably requiring association with nsp14 to maintain solubility, as previously proposed^8^) (Fig. 1a).

**Figure 1.**
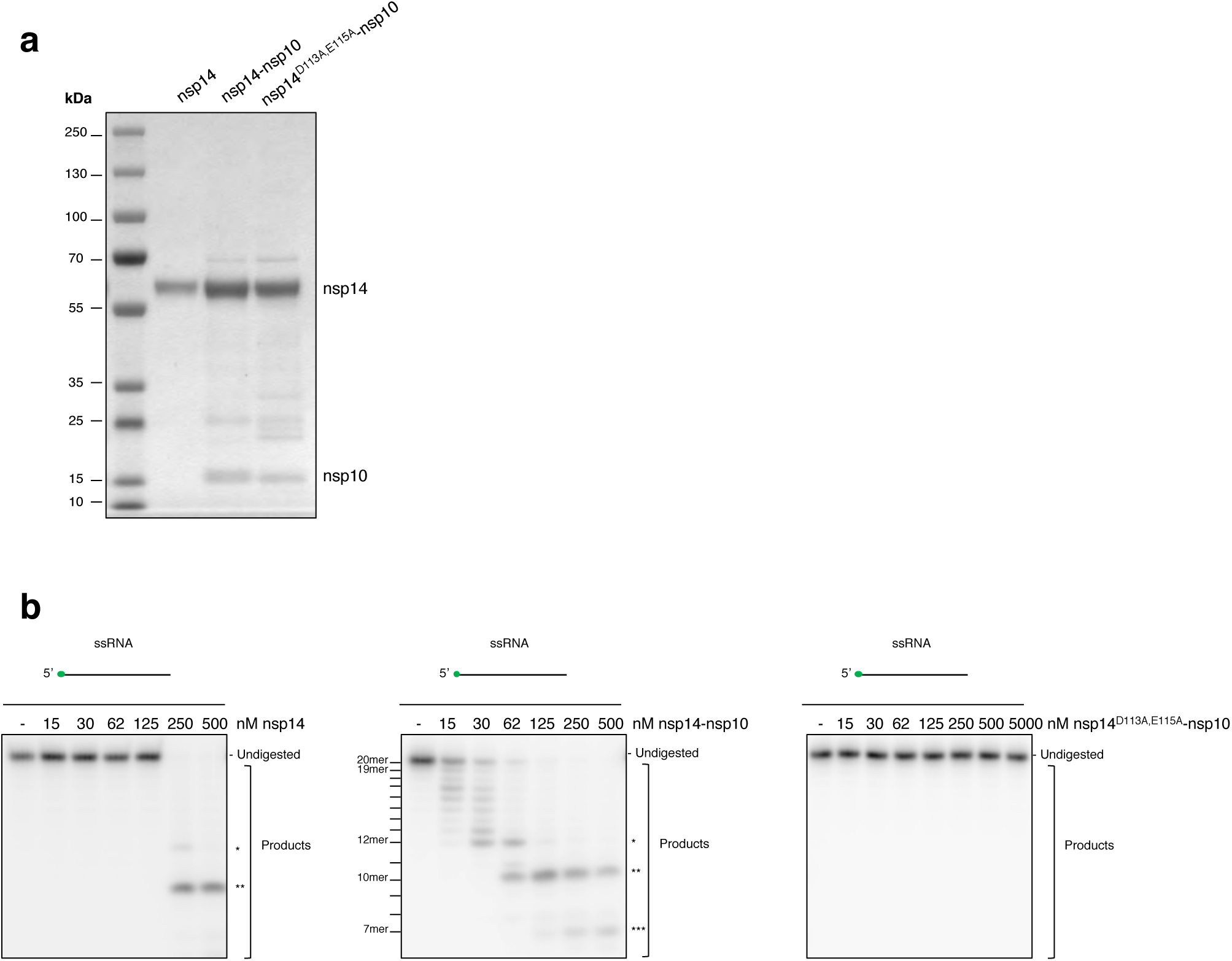
The nuclease activity of nsp14 (ExoN) is substantially increased by nsp10. (a) SDS-PAGE gel of purified nsp14 alone and WT nsp14-nsp10 (nsp14-nsp10) and a control ‘nuclease-dead’ complex bearing Ala-substitutions at D113 and E115 (nsp14^D113A,E115A^-nsp10) showing the purity of the purified proteins. Predicted molecular weights are 59 463 Da for nsp14 and 15 281 Da for nsp10. (b) Nsp14-nsp10 is an RNase that digests a 20-mer ssRNA oligo (oligo 2 as indicated in Suppl. Table 1A) in a single-nucleotide fashion from the 3’-end terminating at the 8^th^ ribonucleotide from the 3’-end (labelled *); it further incises closer to the 5’-end to generate 10-mer and 7-mer products (labelled ** and *** respectively). Nsp14 alone can generate the 12-mer and 10-mer products, but when using significantly higher protein concentrations. The predicted nuclease-dead nsp14^D113A,E115A^-nsp10 mutant complex exhibits no discernible nuclease activity, even at ten-fold higher concentrations compared with the wildtype complex. Increasing concentrations of protein (as indicated) were incubated with substrate at 37°C for 45 min and reactions were subsequently analysed by 20% denaturing PAGE to visualize product formation. Size of products was determined as shown in Suppl. Fig 1D. Main products are labelled *, ** and *** corresponding to 12-mer, 10-mer and 7-mer respectively.

Previous studies of the CoV nsp14-nsp10 complexes have been limited to the SARS-Cov-1 strain proteins^3, 9^. In the latter study, a preferential activity on double-stranded DNA (dsRNA) over single-stranded DNA (ssRNA) combined with an enhanced capacity to process substrates with mismatches within 4-ribonucleotides of a 3′-OH was reported. For the Sars-CoV-2 complex, we performed an extensive evaluation of RNA structures, sequences and modifications to comprehensively define the major activities of the complex. We initially utilised a 20-mer ssRNA substrate radiolabelled at the 5′-end. It has previously been reported that while nsp14 alone is active as a nuclease, its activity is substantially increased by association with nsp10^3, 8^. To test this, we incubated the ssRNA substrate with nsp14 alone, and the nsp14-nsp10 complex. Substrate digestion was observed for nsp14 at concentrations above 250 nM (on 10 nM of substrate). We confirmed that nsp10 dramatically enhances the activities of nsp14, as efficient digestion by nsp14-nsp10 was observed at the lowest concentration employed (15 nM) (Fig. 1b). We next examined the activity of nsp14-nsp10 on sequence-related ssRNA substrates of varying length. Whilst no activity could be observed with a 10-mer, possibly because this substrate is too short to permit stable binding of the nsp14-nsp10 complex, robust activity at concentrations above 250 nM was observed with a 30-mer (Supplementary Fig. 1c). Moreover, for both the 20-mer and 30-mer substrates a complex digestion pattern was observed (Supplementary Fig. 1c). We saw RNA laddering close to the 3′-terminus, consistent with the previously reported 3′-exonuclease activity of this complex, but also several additional prominent additional bands representing cleavage further from the 3′-end. We next precisely identified the size of the major products released by the nsp14-nsp10 on the 20-mer substrate. In order to do this, we performed limited hydrolysis on the 20-mer substrate to provide a single-nucleotide molecular weight marker (Supplementary Fig. 1d). This enabled us to determine that the laddering products observed at the top of the gel correspond to fragments digested in a single-nucleotide fashion from the 3′-end, and that this processing terminates at the 8^th^ ribonucleotide from the 3′-end (labelled with *). Two additional prominent bands were identified, corresponding to cleaved 10^th^ and 13^th^ ribonucleotides from the 3′-end, releasing a 10-mer and 7-mer product (labelled on Fig. 1b and Supplementary Fig. 1d as ** and ***, respectively).

We then confirmed that the activities we observed are intrinsic to the nsp14-nsp10 complex by two complementary methods. First, we purified the predicted nuclease inactive complex nsp14^D113A,E115A^-nsp10 and assayed its activity on the 20-mer substrate. No activity was observed, even at concentrations 10-fold higher that the highest concentration employed with the wild-type complex (5000 nM for nsp14^D113A,E115A^-nsp10 versus 500 nM for nsp14-nsp10; Fig. 1b). Second, we ran the wild-type nsp14-nsp10 complex on a gel filtration (size exclusion) column and then assayed the eluted fractions for RNase activity. The elution profile of the heterodimeric protein from this column precisely coincided with the peak in the characteristic RNase activity we observe (Supplementary Fig. 1e), providing further reassurance that the activities we are observe are intrinsic to the complex we have purified and not due to a contaminant.

Finally, as previous biochemical and structural studies imply that the nuclease activity of the ExoN family of nucleases is dependent upon divalent metal cations, we sought evidence that this is the case for SARS-CoV-2 nsp14-nsp10^3, 8, 9^. First, we determined which divalent cations support maximal activity of the complex. Both magnesium and manganese promoted the RNase activity of nsp14-nsp10, whereas zinc was inhibitory (Supplementary Fig. 2a. Consistent with a requirement for metal ions for activity, three metal-chelating agents, EDTA, EGTA and ο-phenanthroline were strongly inhibitory, with a particular sensitivity to EDTA (Supplementary Fig. 2b). For further studies, we chose to perform reactions in the presence of magnesium as this efficiently supports nsp14-nsp10 activity while being highly abundant in the mammalian cytoplasm.

### Effects of RNA sequence and 3′-terminus structure on nsp14-nsp10 activity

The RNA sequence dependency of CoV nsp14-nsp10 complexes has yet to be examined systematically. Initially, we compared the activity of nsp14-nsp10 on the 20-mer substrate employed throughout Fig. 1 with its activity on a sequence-unrelated 20-mer ssRNA. Both substrates are of mixed sequence and contain all four bases. It is striking that the pattern and efficiency of digestion on these two substrates was indistinguishable suggesting that for RNA substrates of mixed sequence, the precise sequence is not a major determinant of digestion pattern (Fig. 2a). Interestingly, when nsp14-nsp10 was presented with 20-mer poly(A) or 20-mer poly(U) substrates, the activity of the complex was dramatically reduced and qualitatively altered (Fig. 2b). For poly(U), step-wise digestion to the 11^th^ ribonucleotide from the 3′-end was observed, whereas for the poly(A) substrate the pattern of digestion was similar but was curtailed at the 8^th^ or 9^th^ nucleotide from the 3′-terminus. However, it is important to note that the concentration of enzyme required to observe any digestion on the poly(U) and poly(A) substrates (250 nM) is substantially higher than those required to efficiently digest RNA substrates of mixed sequence (compare Figs. 2a and 2b). We tentatively propose that nsp14-nsp10 might possess reduced activity on poly(U) and poly(A) sequences to prevent the degradation of the 3′-tails of both viral and host transcripts, which could be inhibitory to efficient viral replication. Notably, unlike poly(A) tails in positive sense viral transcripts, poly(U) sequences in negative sense transcripts can act as pathogen-associated molecular patterns (PAMPs) and alert the innate immune system. Poly(U) transcripts can be degraded by CoV nsp15 which is perhaps dedicated to performing this reaction, but in a regulated fashion^17^.

**Figure 2:**
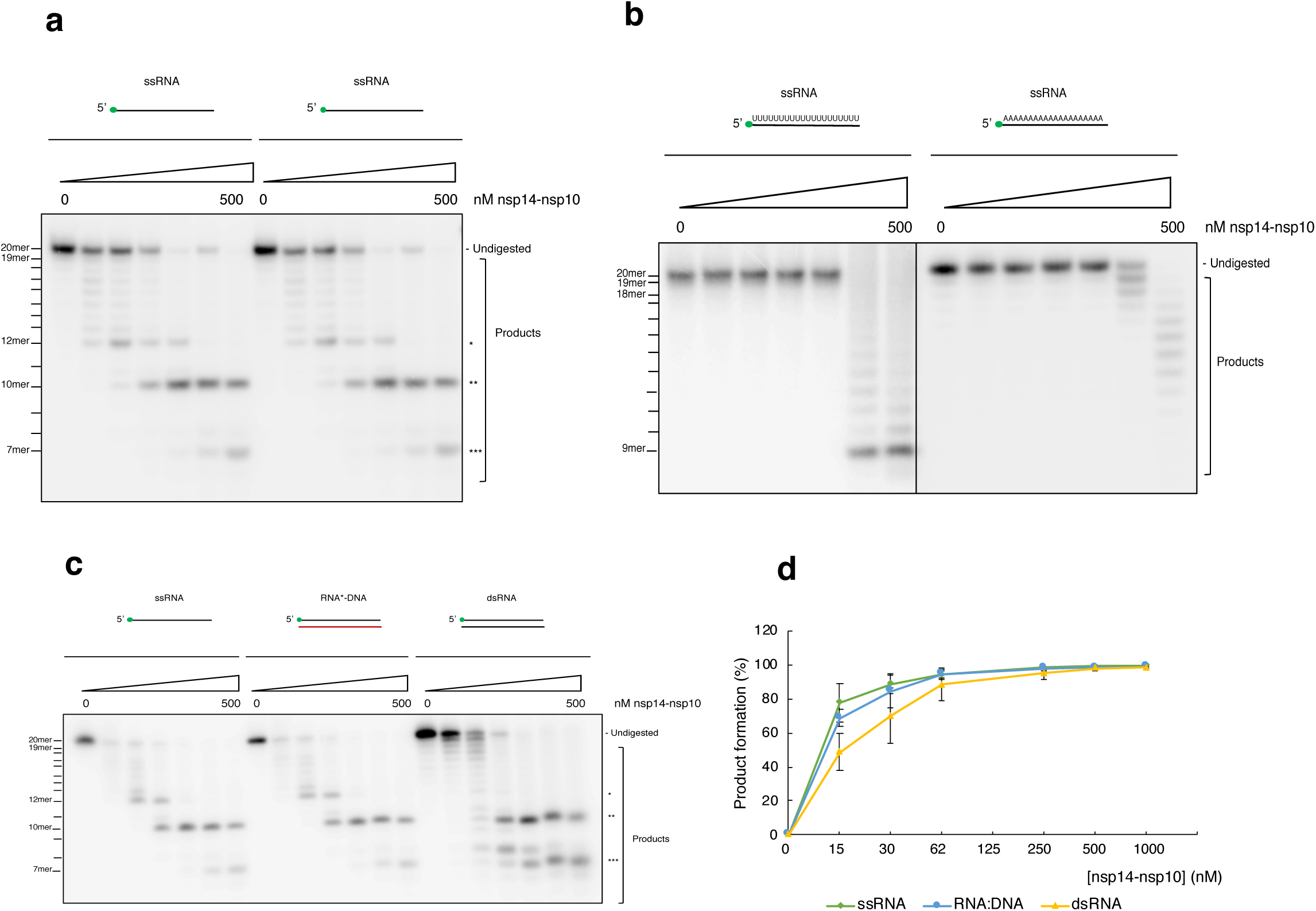
Nsp14-nsp10 is a versatile RNA nuclease. (a) Nsp14-nsp10 nuclease activity is not sequence specific. The complex shows indistinguishable digestion patterns on two 20-mer ssRNA substrates of mixed sequence and containing all four bases. Oligos 2 and 3 were used, respectively (see Suppl. Table 1A). (b) When presented with 20-mer Poly(U) and Poly(A) ssRNA (oligos 13 and 14 respectively; Suppl. Table 1A), nsp14-nsp10 shows reduced and qualitatively altered activity, with a single nucleotide step-wise digestion from the 3’-end curtailing at the 9^th^-11^th^ nucleotide from the 3’-end. (c) Nsp14-nsp10 processes ssRNA, the RNA strand of an RNA:DNA hybrid, and dsRNA with no preference for double-stranded substrates. For all structures, the labelled strand is oligo 2, also used in Fig. 1b (see Suppl. Table 1A and B). (d) Product formation (%) was quantified for Fig. 2 C comparing nsp14-nsp10 nuclease activity on ssRNA, the RNA strand of an RNA:DNA hybrid and dsRNA as outlined in Methods and Materials. All data are shown as mean ± s.e.m, and at least three biological replicates were used for each substrate. Nsp14-nsp10 shows equivalent activity on ssRNA and the RNA strand of an RNA:DNA hybrid with a slight decrease in activity for dsRNA. Increasing concentrations of protein (as indicated) were incubated with substrate at 37°C for 45 min; reactions were subsequently analysed by 20% denaturing PAGE to visualize product formation. The s of products was determined as shown in Suppl. Fig 1D. Main products are labelled *, ** and *** corresponding to 12-mer, 10-mer and 7-mer respectively.

A key difference between our observations and those previously reported for the SARS-Cov-1 complex relate to the ability of nsp14-nsp10 to process ssRNA substrates; for SARS-CoV-1 nsp14-nsp10, a strong preference for dsRNA over ssRNA substrates was reported^3^. To examine if the Cov-2 complex also exhibits a preference for ds-over ssRNA, we tested its activity on sequence-identical ssRNA, dsRNA, and hybrid RNA:DNA substrates (where the RNA strand is 5′-radiolabelled) (Fig. 2c). We did not observe a preference for dsRNA over ssRNA, in fact the converse was observed since higher concentrations of nsp14-nsp10 were required to produce the same extent of digestion for dsRNA when compared to ssRNA (Fig. 2c and quantified in Fig. 2d). Moreover, the enzyme appears to be agnostic to the nature of the strand annealed to the RNA substrate strand, since the RNA component of the RNA:DNA hybrid was digested with efficiency equal to the ssRNA substrate (Fig. 2c). Previous studies of SARS-CoV-1 nsp14-nsp10 concluded the complex possesses 3′-exonuclease activity, since activity was reduced when the 3′-hydroxyl terminus of the substrates examined was replaced by a phosphate group^3^. To examine this point for the SARS-CoV-2 complex, we compared the activity of nsp14-nsp10 on three sequence-identical substrates bearing either a 3′-hydroxyl, 3′-phosphate, or 3′-biotin moiety (Fig. 3a). Interestingly, the characteristic ladder of products extending from the full-length substrate (at the top of the gel) was strongly reduced when the 3′-hydroxyl was replaced by a biotin group only. However, the other major products (7-mer and 10-mer) observed on the substrate with the hydroxyl terminus were variably affected when the 3′-end was replaced with a phosphate or biotin. In the presence of the 3′-phosphate, the 10-mer product was still observed, being produced in approximately equal amounts as for the 3′-hydroxyl substrate, while the 7-mer product was observed at slightly higher concentrations (above 250 nM) compared to the 3′-hydroxyl substrate. With a 3′-biotin substrate, higher enzyme concentrations were required to observe cleavage with predominantly the 10-mer product being observed at concentrations above 250 nM. The 12-mer product was still observed at concentrations above 125 nM, albeit very faintly, whilst the 7-mer product was only observed at the highest concentration (500 nM). Together, these observations strongly suggest that SARS-CoV-2 not only has a 3′-exonuclease activity, which is dependent upon the presence of an unblocked 3′-RNA terminus (leading to the laddering products we characteristically observe at the top of the gels), but also an endonucleolytic activity, cleaving at positions further from the 3′-end. These incisions occur independently of the presence of a 3′-hydroxyl or phosphate group, and therefore do not require engagement with a 3′-terminus prior to initial exonucleolytic processing at the 3′-terminus for their production. Interestingly, in the presence of manganese the main cleavage product is the 7-mer, suggesting that this ion enhances the endonuclease activity (Supplementary Fig. 2a).

**Figure 3:**
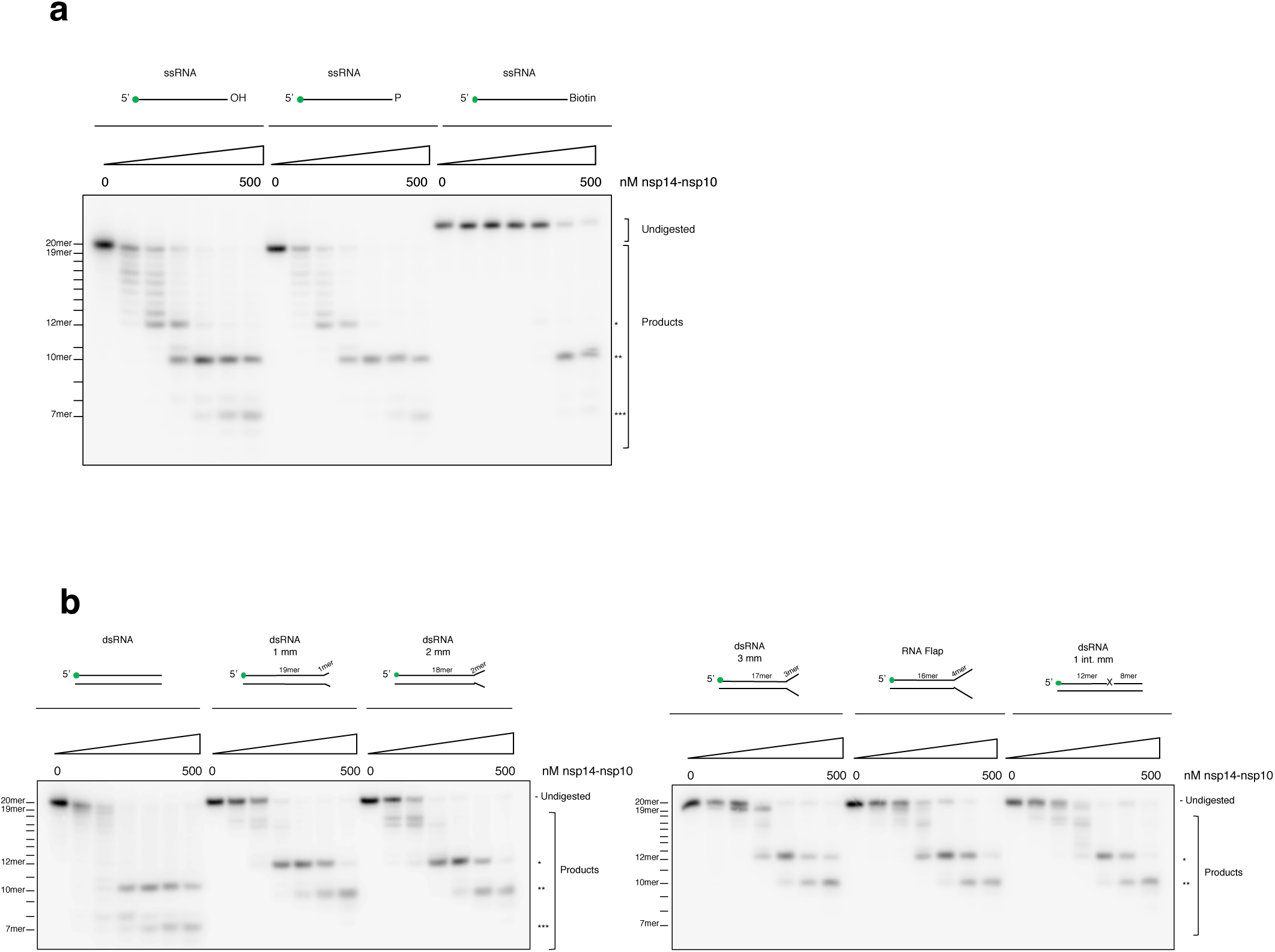
Nsp14-nsp10 of Sars-CoV-2 exhibits both 3’-exonuclease activity and a newly-described endonucleolytic activity that reaches beyond the classical role of a proofreading nuclease. (a) Nsp14-nsp10 is an RNA exo- and endo-nuclease. With a substrate containing a 3’-biotin group, the characteristic laddering of the substrate is lost and only endonucleolytic cleavage at the positions furthest from the 3’-end is observed. Substrates with a 3’-hydroxyl or phosphate group, exhibit near identical product profiles. (b) Nsp14-nsp10 is an exo- and endo-nuclease able to incise a variety of RNA substrates, including RNA substrates with mismatched termini and flaps with no preference for mismatched ribonucleotides. Quantification in Suppl. Fig. 3B (mm: mismatch, int. mm: internal mismatch). Increasing concentrations of protein (as indicated) were incubated with substrate at 37°C for 45 min; reactions were subsequently analysed by 20% denaturing PAGE to visualize product formation. Size of products was determined as shown in Suppl. Fig 1D. Main products are labelled *, ** and *** corresponding to 12-mer, 10-mer and 7-mer respectively. All oligos used are indicated in Suppl. table 1A and B.

To date, CoV nsp14-nsp10 complexes have been reported to act exclusively on RNA. To confirm that this is the case for the Sar-CoV-2 complex, we compared the activity of nsp14-nsp10 on our 20-mer ssRNA substrate with its activity on sequence-related 20-mer ssDNA and DNA:RNA hybrid substrates (where the DNA strand is 5′-radiolabelled) (Supplementary Fig. 3a). As expected, no activity was observed on either DNA substrate. We also examined the ability of nsp14-nsp10 to act on substrate containing a single ribonucleotide embedded within an 18-mer DNA duplex (3-ribonucleotides form the 3′-end), analogous to the types of substrate processed by ribonucleotide excision repair (RER) enzymes (Supplementary Fig. 3a)^18^. Some activity was observed at concentrations above 125 nM with the two products released being consistent with incisions being introduced 3′- and 5′-to the site of the embedded ribonucleotide. While not necessarily physiologically relevant, this activity is informative regarding the proposed endonucleolytic activity of this complex, since the 3′-DNA terminus of this substrate is completely resistant to nsp14-nsp10, but nonetheless nsp14-nsp10 can cleave adjacent to the single embedded ribonucleotide.

### Impact of mismatches, flaps and chemical modification on nsp14-nsp10 activity

It has been reported that the SARS-CoV-1 nsp14-nsp10 complex preferentially degrades substrates containing mismatches of up to 4-ribonucleotides at their 3′-termini^3, 10^. To address this possibility for the SARS-CoV-2 complex, we generated substrates containing 1-, 2-, 3- or 4-ribonucleotide mismatches at their 3′-termini, where the 4-ribonucleotide mismatch effectively introduces a small 3′-flap allowing us to examine the impact of this secondary structure on activity. We observed no substantial difference in digestion efficiency across any of these four structures (Fig. 3b and quantified in Supplementary Fig. 3b), although there were qualitative differences in the major products released, likely reflecting differential modes of association of the complex due to the structural variations in the substrates. We also examined a substrate containing a single mismatch near its centre (8-ribonucleotides from the 3′-end). This was also processed with an efficiency indistinguishable from the other substrates, exhibiting a qualitative pattern of digestion similar to the substrates containing the terminal mismatches (Fig. 3b and quantified in Supplementary Fig. 3b). Taken together, our results show that mismatched termini are not preferentially processed by the SARS-Co-V-2 nsp14-nsp10 complex, but rather the complex is an exo- and endo-nuclease able to act on a wide variety of RNA substrates, both single- and double-stranded.

Finally, we examined whether some common modifications of viral RNA impact the RNase activity of nsp14-nsp10. We investigated the impact of 6-methyladenine, one of the most common modifications found in cytoplasmic RNA, along with an artificial base modification, 2-methyladenine. Importantly, 6-methyladenine modification has been associated with viral evasion by the host innate immune system^19^. We also examined the variant RNA base inosine, that can be generated by A-to-I editing and has been reported to enhance viral recognition by the innate immune sensors^20^. When any of these modified bases was placed two ribonucleotides from the 3′-terminus of the 20-mer substrate, none affected, either qualitatively or quantitatively, activity of nsp14-nsp10 when compared with a native (unmodified) RNA substrate (Supplementary Fig. 3c). We conclude that some common chemical modifications of RNA do not substantially affect nsp14-nsp10 RNase activity.

### Identification of nsp14-nsp10 inhibitors

The urgent need to identify therapeutics to treat COVID-19 then led us to scope potential nsp14-nsp10 inhibitors, with a focus on compounds identified as potential COVID-19 treatments and known nuclease inhibitors. Previous studies suggest that inactivation of nsp14-nsp10 can lead to increased genomic instability and therefore a potentially lethal mutagenesis and concomitant reduction in viral propagation. While this alone might be insufficient to treat COVID-19 effectively, the possibility that nsp14-nsp10 inhibitors could be combined with inhibitors of other key factors (for example, the major proteases 3CL^pro^/M^pro^ or replicative RNA polymerase catalytic subunit nsp12) merits exploration.

Based on the efforts of ourselves and others to identify inhibitors of RNA and DNA nucleases, we first performed *in silico* docking experiments focusing on chemotypes known to inhibit nucleases. We employed the AutoDock Vina^21^ platform to dock in-house compounds on the entire exposed surface (blind docking) and the surface centred on the active site of a structure of SARS-CoV-1 nsp14-10 (PDB:5NFY)^10^. Use of a homology model of SARS-CoV-2 nsp14-nsp10 created using PDB:5NFY was considered inappropriate due to the low resolution of this structure. The docking analyses revealed a range of pharmacophores that may interact with the enzyme (Supplementary Fig. 4a), and one of these, an *N*-hydroxyimide compound (denoted A-1) was identified as potentially interacting with the nsp14-nsp10 active site (Fig. 5a; Supplementary Fig. 4a).

**Figure 5:**
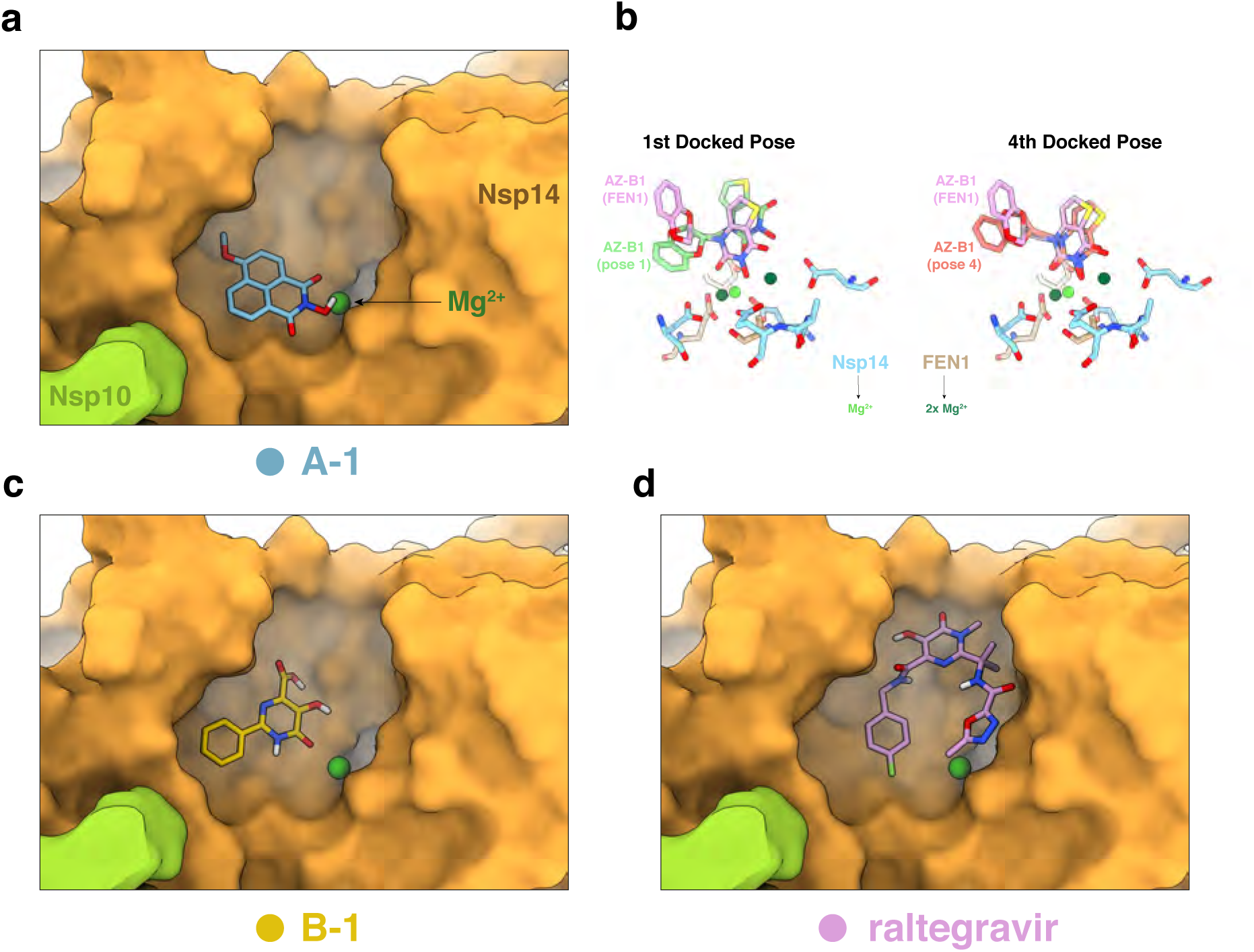
Docking of potential inhibitors SARS-CoV nsp14-nsp10. Nsp14-nsp10 was docked with compounds within grid boxes encompassing a surface focussed on the active site surface using the reported Nsp14-nsp10 structure (PDB 5NFY) and Autodock (see Materials and Methods for details). (a) The highest-affinity docking pose of A1 overlaid on the surface of SARS-CoV nsp14-nsp10. (b) Comparison of docked interaction between and AZ-B1 and nsp14 Mg^2+^ ion with a crystal structure of AZ-B1 in complex with the nuclease FEN1.Mg^2+^ (PDB: 5FV7). (c)&(d) The highest-affinity docking poses of B1 and raltegravir overlaid on the surface of SARS-CoV nsp14-nsp10. Views of the nsp14 surface in panels b, d–e are matched; nsp14 is in yellow-orange, nsp10 is in light green, Mg^2+^ is in dark green.

Efforts to establish a quantitative fluorogenic assay for nsp14-nsp10 activity, analogous to those we and others have devised for endo- and exonucleases previously^22^, have as yet been unsuccessful due to the exquisite sensitivity of the nsp14-nsp10 complex to substrate modification by all fluor and quench moieties tested. We therefore utilised a gel-based nuclease assay employing the 20-mer ssRNA described above to examine the potential inhibitory characteristics of the compound A-1, and its positional isomer A-2, available to us from existing chemical libraries. The chemical aurintricarboxylic acid (ATA), which is a promiscuous ribonuclease inhibitor used during nucleic acid extraction protocols was used as a positive control^23^; as expected, inhibited nsp14-nsp10 with an IC_50_ of 7.6 ± 1.1 μM (Fig. 4; Supplementary Fig. 5). Fig. 4a shows the inhibitory characteristics of A-1 and A-2 *N*-hydroxyimides in the gel-based assay, each of which exhibit qualitatively complete inhibition of substrate digestion (Fig. 4a). Quantification of the gel-based nuclease assay data did not yield an IC_50_ value for A-1, however, for A-2 this was determined to be 20 ± 0.5 μM (Fig. 4b, Supplementary Fig. 5). Based on the docking data of A-1 and potency of A-2 in the gel-based assay, we synthesised three structural variants of these compounds. We decided to remove one of the rings, and use a bicyclic hydroxyimide scaffold, as has previously been used against the nucleases FEN1 and XPF-ERCC1^24, 25^; these compounds were denoted A-3 and A-4. We also attempted to introduce a thiocarbonyl (A-5) to examine the potential effects on inhibition (Supplementary Fig. 6). However, none of these exhibited a lower IC_50_, and therefore greater potency than A-2 (Fig. 4; Supplementary Fig. 6). It is important to note that due to nature of the gel-based assay, and the quality of the curves extracted, that the absolute IC_50_ values should be regarded as preliminary, though complete inhibition of nuclease activity is observed for A-1, A-2, A-3, B-1 and raltegravir at concentrations of 100 μM.

**Figure 4:**
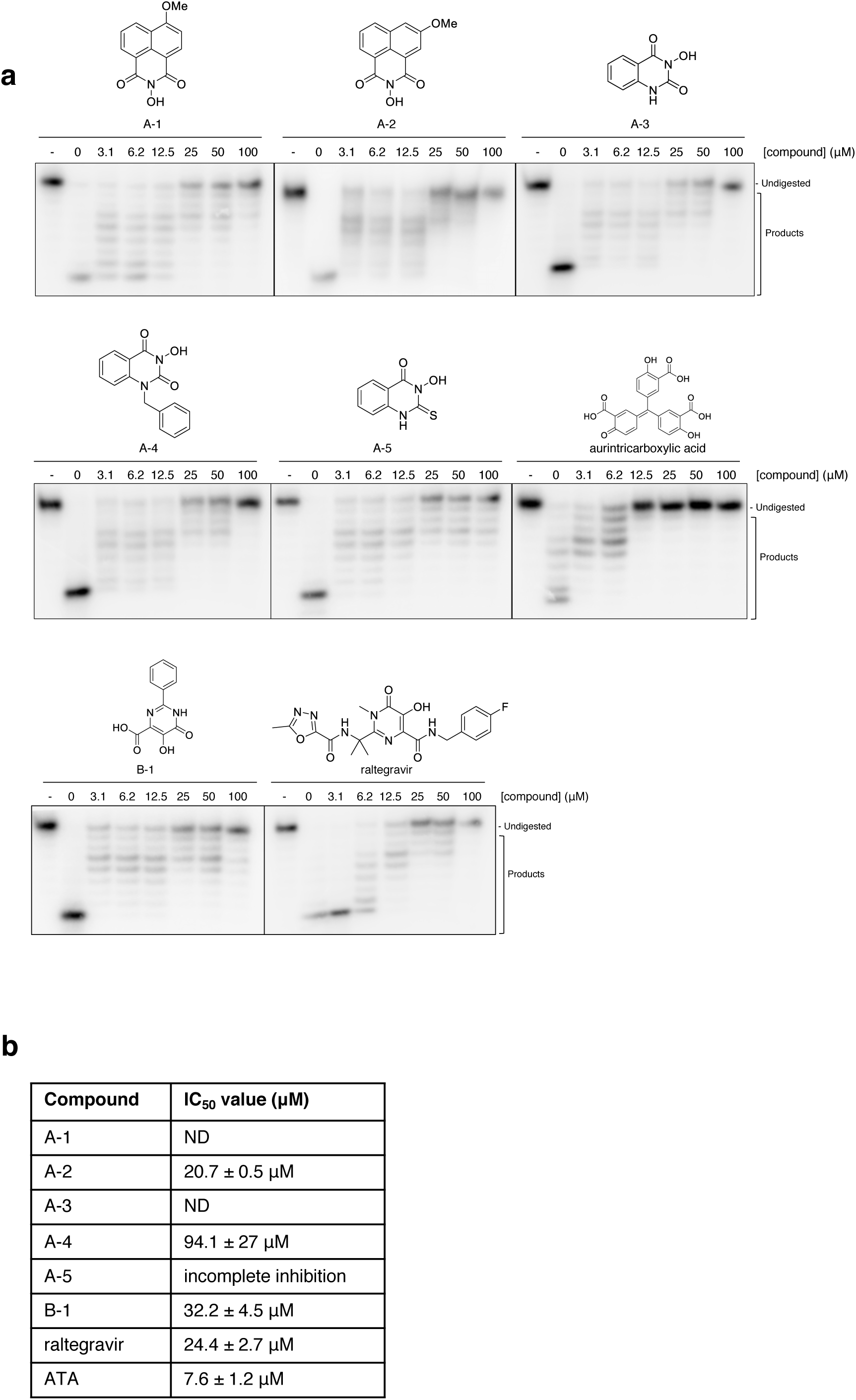
The exonuclease activity of nsp14-nsp10 is inhibited by *N*-hydroxyimide and hydroxypyrimidinone containing compounds. (a) Increasing concentrations (as indicated, in μM) of the indicated compounds were incubated with 100 nM nsp14-nsp10 (10 minutes, room temperature), before initiating a standard nuclease assay by the addition of ssRNA and incubating at 37°C for 45 min. Products were analysed by 20% denaturing PAGE. A decrease in the generation of nucleolytic reaction products and a concomitant increase in undigested substrate indicates inhibition of nuclease activity at increasing inhibitor concentrations. - indicates no enzyme. Compounds A-1–A-4 are based on a *N-*hydroxyimide scaffold, B-1 is a hydroxypyrimidinone. (b) IC50 values as calculated by quantification of gel digestion products and dose-response curves were determined by nonlinear regression. The mean ± s.e.m. were calculated from ≥ 3 biological repeats.

The *N*-hydroxyimide pharmacophore based is reported to inhibit structure-selective DNA nucleases, *via* chelation of an active site metal ion(s) (particularly Mg^2+^)^24, 25^. Our *in silico* docking shows potential for *N*-hydroxyimide compound binding at the nsp14-10 active site, with polar inhibitor atoms positioned close to the catalytic Mg^2+^; wherein for A-1 two of the hydroxyimide oxygen atoms are a distances of 2.4 Å and 2.6 Å from the Mg^2+^ (Fig. 5A).The crystal structure of the human FEN1 nuclease with an *N*-hydroxyimide-based compound, AZ13623940 (denoted AZ-B1), bound to the active site has previously been reported (PDB: 5FV7)^26^. To allow possible mechanistic insight, the docking pose of AZ-B1 and its interaction with the catalytic Mg^2+^ in the nsp14-nsp10 model was compared with a structure of FEN1 in complex with AZ-B1 (Fig. 5b). While there was little overlap of the highest predicted affinity active site-docked pose of AZ-B1 with the AZ-B1 binding mode in the FEN1 structure, the fourth pose did align well with the Mg^2+^ of nsp14-nsp10, close to a FEN1 Mg^2+^ (Fig. 5b), also validating the ability of AutoDock Vina to place compounds near divalent metal ions.

Taking advantage of previous identification and detailed characterisation of a series of FEN-1 inhibitors^26^, we determined whether this molecule (AZ-B1), and AZ1353160 (denoted AZ-A1) are also capable of inhibiting SARS-CoV-2 nsp14-nsp10. While some inhibition was observed (Supplementary Fig. 7), these were also inferior to A-2.

The N-hydroxypyrimidinone ring is structurally similar to N-hydroxyimide and hydroxypyrimidinones inhibit nucleases through binding to the active site metal ion(s), typically Mg^2+^^27, 28^. Therefore, we tested the commercially available compound 5,6-dihydroxyl-2-phenylpyrimidine-4-carboxylic acid (denoted B-1). B-1 exhibited a comparable inhibition profile to A-2 with an IC_50_ value of 32.2 ± 4.5 μM (Fig. 4; Supplementary Fig. 5). We also tested raltegravir, an HIV integrase inhibitor, which contains a hydroxypyrimidinone ring^27, 28^. Similar to B-1, raltegravir exhibited clear inhibition of the exonuclease activity of nsp14-nsp10 with an IC_50_ value of 24.4 ± 2.7 μM (Fig. 4; Supplementary Fig. 5). Our *in silico* approach modelled B-1 docked proximal to the active site of nsp14-nsp10, with an oxygen atom of the hydroxypyrimidinone coordinating the catalytic Mg^2+^, at a distance of 2.7 Å (Fig. 5c). Docking shows B-1 can bind at the active site of nsp14-10, with an oxygen atom of the hydroxypyrimidinone coordinating the catalytic Mg^2+^(2.7 Å) (Fig. 5c). The docked pose of raltegravir differed from that of B-1, however, it was again positioned close to the active site Mg^2+^, making additional contacts within the putative substrate binding pocket (Fig. 5D).

These experiments provided important insight into pharmacophores that may be further developed and tested as COVID-19 treatments. However, the urgency of addressing COVID-19 requires that experimental or approved drugs can be repurposed for treatment. We therefore undertook at low-throughput (limited by the lack of a scalable fluorescent assay) screen of candidate drugs that might be predicted, on the basis of their mechanism of action, to inhibit the nuclease activity of nsp14-nsp10. These included nucleoside analogues, topoisomerase poisons, candidate DNA repair inhibitors, compounds reported to inhibit other COVID-19 targets and antivirals believed to interfere with nucleic acid metabolism. The compounds tested and the results from the gel-based nuclease assay screening are summarised in Supplementary Table 2). Of these, ebselen (an organoselenium molecule with broad pharmacological properties^29, 30^ and a known potent inhibitor of the main CoV protease 3CL^pro^) and disulfuram (a carbothioamide used to treat alcohol dependence^31^) appeared most active in our screen. Detailed analysis allowed us to determine that the IC_50_ for ebselen and disulfuram are 3.3 ± 0.09 μM and 89 ± 33 μM, respectively (Fig. 6 and Supplementary Fig. 5). We also included thiram, a non-drug compound (used as a fungicide) which is also a carbothioamide related to disulfuram to determine whether this chemotype more generally acts to inhibit nsp14-nsp10. The IC_50_ for thiram was 48.2 ± 1.8 μM, suggesting that further surveys of carbothioamides are warranted in the search for nsp14-nsp10 inhibitors.

**Figure 6:**
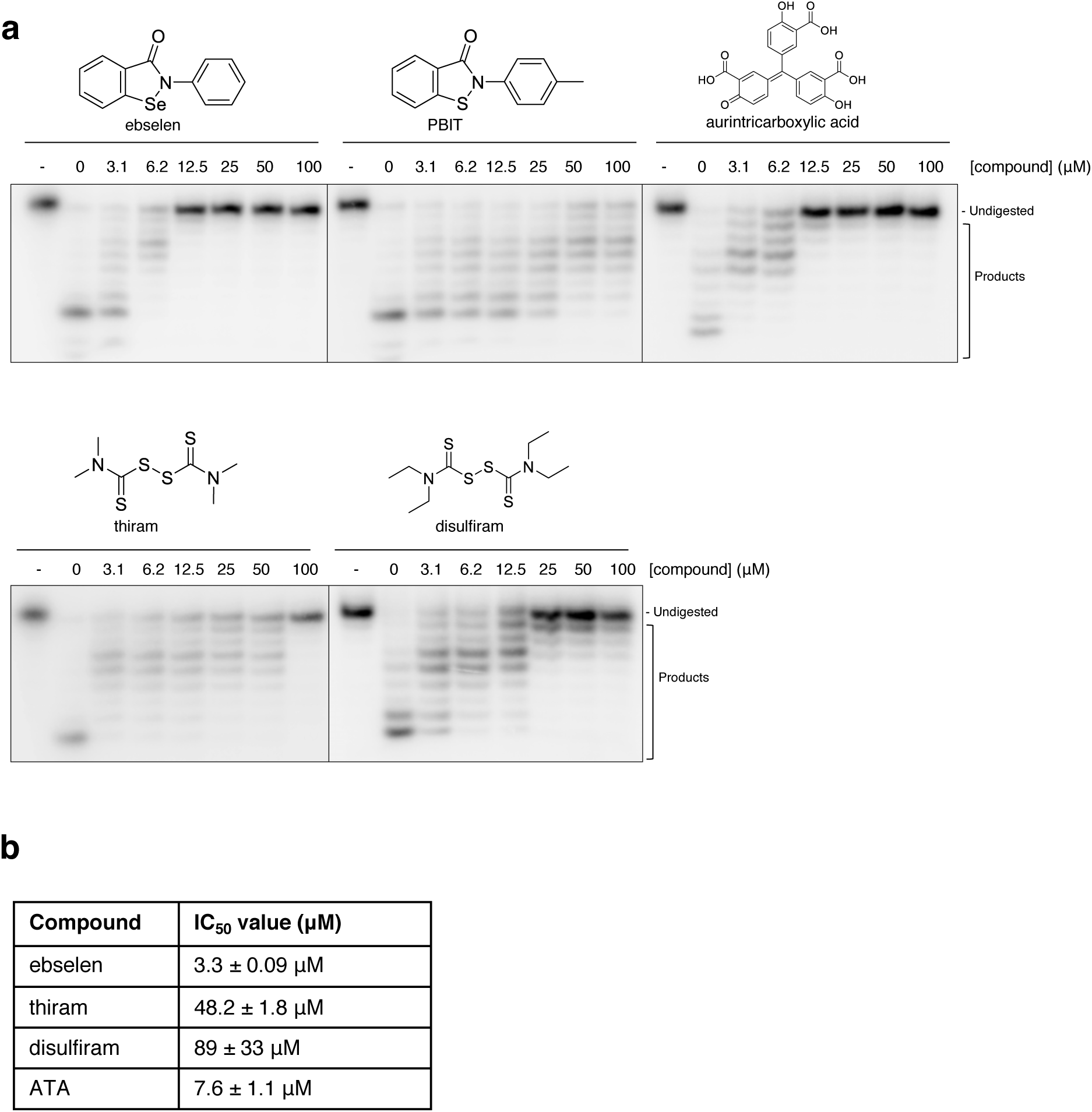
The exonuclease activity of nsp14-nsp10 is inhibited by the presence of drug and drug-like compounds. (a) Increasing concentrations (as indicated, in μM) of drug and drug-like compounds were incubated with 100 nM nsp14-nsp10 for 10 min at room temperature, before starting a standard nuclease assay reaction by the addition of ssRNA and incubating at 37°C for 45 min. Reaction products were analysed by 20% denaturing PAGE. A decrease in the generation of nucleolytic reaction products and a concomitant increase in undigested substrate indicates inhibition of nuclease activity. - indicates no enzyme. Gels are representative of at least three biological repeats. (b) IC50 values as calculated by quantification of gel digestion products and dose-response curves were determined by nonlinear regression. The mean ± s.e.m. were calculated from ≥ 3 biological repeats.

To ensure the compounds that exhibited inhibitory effects were not exerting these through aggregation or precipitation of nsp14-nsp10, we employed differential scanning fluorimetry (DSF) under identical reaction conditions to the inhibitor nuclease assays. None of the inhibitors showed any destabilising effect on thermal stability of nsp14-nsp10, with the exception of ATA at higher concentrations (Supplementary Fig. 8).

Ebselen has generally well-tolerated in clinical studies investigating its effectiveness for treating a range of conditions from stroke and hearing loss to bipolar disorder^29^. One mechanism of action proposed for ebselen relates to the potential capacity of the selenium to react with the thiolate ligands of zinc clusters in proteins and to release zinc^32–35^. To test whether the selenium is critical for the inhibition of nsp14-nsp10 nuclease activity, we investigated an analogue of this molecule, PBIT, where a sulphur atom replaces the selenium. Consistent with an important role for the selenium in nsp14-nsp10 inhibition, PBIT demonstrated a dramatically reduced potency of inhibition (Fig. 6a). We conclude that the best, potentially clinically useful nsp14-nsp10 inhibitor we have identified is the M^pro^ inhibitor ebselen, where the selenium moiety plays an important role mediating its inhibitory effects.

In conjunction with our low-throughput nuclease assay inhibitor screening, all compounds tested were screened for binding to SARS-CoV-1 nsp14-nsp10 using AutoDock Vina. There was no clear relationship between the calculated docking affinities and the extent of exonuclease inhibition *in vitro,* either for the active site or full surface docking (Supplementary Fig. 4 and Supplementary Table 2), in part reflecting the likely covalent inhibition by compounds such as ebselen, underlining the need for experimental validation of hits proposed by computational docking simulations. Interestingly, the majority of primary docked poses were located in the nsp14 ExoN-MTase domain boundary, within a pore that is otherwise occupied by substrates of the methyltransferase reaction, G^5′^ppp^5′^A and *S*-adenosyl methionine (PDB: 5C8S) (Supplementary Fig. 4c–e)^8^. While compounds docked at this site may be unlikely to have an impact on competitive inhibition of nsp14-nsp10 nuclease activity, this suggests that the MTase active site may also be an attractive target, whereby compounds may hinder the binding of MTase cognate substrates.

## DISCUSSION

Here we present the first biochemical analysis of the nsp14-nsp10 protein complex, that plays a key role in the genome duplication of the SARS-CoV-2 virus, the causative agent for COVID-19. Previous studies of the related SARS-CoV-1 complex have primarily focused on structural aspects^8, 10^. The 3′-exonuclease activity of the CoV-1 was complex was reported, but it was not subject to detailed biochemical analysis.

Our study confirms the presence of a 3′-exonuclease activity in the Cov-2 nsp14-nsp10 complex, which is curtailed when the 3′-hydroxyl terminus of the RNA substrate is changed to a biotin group. Altering the nature of the 3′-terminus, however, revealed a further feature of nsp14-nsp10 activity, i.e. the capacity of the enzyme complex to catalyse incisions further into (closer to the 5′-end) of the substrate. We observed a robust and almost equivalent activity of nsp14-nsp10 on both ssRNA and dsRNA. Related, we did not observe preferential excision of terminally mismatched nucleotides by the CoV-2 nsp14-nsp10 complex, as has been reported for the analogue CoV-1 complex^3^. Instead, therefore, we observed that nsp14-nsp10 is capable of degrading (approximately) the first eight ribonucleotides from the 3′-end of all substrates tested (either ssRNA or dsRNA) in an exonucleolytic fashion. When a 3′-hydroxyl group was not present the nsp14-nsp10 nuclease acted further downstream (5′-to) these sites to endonucleolytically release fragments of 7- to 10-ribonucleotides endonucleolytically. We therefore propose that the role of nsp14-nsp10 is more than that of a proof-reader, by the analogy to the 3′-exonuclease present in replicative DNA polymerase complexes^11^. We propose nsp14-nsp10 might represent a simple replication-repair system potentially capable of removing ribonucleotides from the 3′-end of elongating replicating RNA strand when extension is blocked. It is likely that the activity of the nsp14-nsp10 is constitutively active during viral replication. It is possible that only when replication stalls due to mismatch incorporation or the presence of altered ribonucleotides that cannot be further extended by the polymerase, the nuclease degrades and/or cleaves the elongating strand, so removing the aberration and allowing for re-engagement of the RNA polymerase and resumption of synthesis. It will be important to reconstitute this replication-repair reaction in future studies to test this hypothesis. We note that another key feature of nsp14-nsp10 nuclease activity is its insensitivity to common RNA modifications that are associated with viral evasion (6-methyladenine) or induction (inosine) of innate immune pathways.

An important question pertains to why we have observed catalytic activities not previously reported in the analyses of CoV-1 nsp14-nsp10. We tentatively propose an explanation based on our experience of working with nsp14-nsp10, namely that the purified complex is exceptionally unstable under frozen storage, even after a period as short as one week. The assays presented here were all performed on nsp14-nsp10 that was less than one week old; after this time elapsed, we found that its activity diminished dramatically, and its behaviour altered. It is possible that degradation and/or aggregation both reduces nsp14-nsp10 activity and alters its substrate selectivity. It is notable that the protein concentrations we employed in our assays, and which supported good activity on ssRNA and dsRNA substrates (15 nM), were far lower than those employed in previous studies that reported on the activity of the CoV-1 complex (several hundred nM), consistent with this suggestion^3, 8^.

Due to the urgency of identifying new drugs and candidate agents that can treat COVID-19, we also performed both an *in silico* docking screen to identify chemotypes capable of interacting with the metal-containing active site of nsp14, and a focused screen of drugs (approved and in development) that might inhibit nsp14-nsp10. Our docking simulations, employing structural data for the highly homologous Co-V nsp14-nsp10 complex, revealed that both *N-*hydroxyimide based compounds could potentially interact with the active site magnesium ion. Testing supported this possibility, revealing one compound from a library available to us, A-2 with an IC_50_ of 20.7 ± 0.5 μM. In the limited time available, we were able to generate several chemical variants around this pharmacophore, but none to date out-performed A-2, nor did two related compounds previously developed to inhibit the FEN-1 DNA nuclease^26^. We expanded our search to explore the putative inhibitory potential of a hydroxypyrimidone scaffold. Compound B-1 exhibited comparable levels of inhibition to the *N-*hydroxyimide based compounds, as did the HIV integrase inhibitor, raltegravir, with IC_50_ values of 32.3 ± 4.5 μM and 24.4 ± 2.7 μM, respectively.

Using docking we were able to model binding of A-1,3–5, B-1, and raltegravir to nsp14-nsp10 using with the structure of SARS-Cov-1. In accordance with previous literature, the *N-*hydroxyimide and hydroxypyrimidone scaffolds chelated the active site magnesium ion of the SARS-Cov nsp14-nsp10^24, 26^, thus providing insight into the possible mode of inhibition. Interestingly, the *in silico* modelling with raltegravir suggests the larger compound could be accommodated well in the active site pocket, potentially making additional contacts, which has the possibility of being explored and/or exploited in future approaches to generate targeted inhibitors of nsp14-nsp10.

Our focused screens of drug and drug candidates with predicted potential to interact with or inhibit nucleic acids processing enzymes revealed that the selenium-containing drug ebselen was an effective inhibitor of nsp14-nsp10. It is interesting to note that ebselen was the most potent inhibitor experimentally confirmed inhibitor of the major SARS-Cov-1 protease (M^pro^), and the authors demonstrated antiviral activity in cell-based assays^30^. The potency of ebselen for nsp14-nsp10 and M^pro^ inhibition are similar *in vitro*; the IC_50_ for M^pro^ was 4.67 μM, and for nsp14-nsp10 we determined an IC_50_ of 3.3 μM. The mechanism(s) of action of ebselen in nsp14-nsp10 inhibition remains unknown, but given the known role for the selenium in this compound to lead to co-ordination or ejection of zinc atoms in these proteins^33–35^, we propose this is a possible mode of action. Indeed, zinc fingers are highly represented in the proteome of SARS-CoV-2^30^, where the M^pro^ possesses zinc binding motifs and the SARS-CoV-1 (and by extension, SARS-CoV-2, due to very high level of sequence conservation) nsp10 possesses at least one zinc finger which is likely to be important for mediating interactions with the RNA viral genome^8, 10^. Interference with these zinc atoms from these binding motifs might induce conformational changes that are not compatible with robust activity. Given that ebselen is capable of targeting two key enzymes required for the propagation of SARS-CoV-2, it is reasonable that further studies assessing the utility of this agent in combatting COVID-19, as a stand-alone agent or in combination, are warranted. More generally, given that many important antimicrobial medicines act by binding to multiple targets (e.g. β-lactams such as penicillin and cephalosporins), we suggest that polypharmacology be pursued as a matter of priority for COVID-19 treatment.

## METHODS

### Expression and purification of wildtype and nuclease dead (NSP14^D113A/E115A^) NSP14-10 complex

The N-terminally His-tagged nsp14-10 protein complex was expressed as a bi-cistronic mRNA in which nsp14 bearing a N-terminal Histidine tag is followed by a ribosome binding site and the untagged nsp10 ORF. The protein was generated by transformation into *E. coli* BL21 Rosetta2 cells and expression in Terrific Broth media supplemented with 30 μg mL^-1^ kanamycin and 10 mM ZnCl_2_. The culture was incubated with shaking (200 rpm) at 37 °C until an OD_600_ of 2.0 was reached. Cultures were then transferred to an incubator at 18 °C for 30 min before induction with 1 mM Isopropyl β-D-1-thiogalactoyranoside (IPTG) and incubated for 18 hours with shaking (200 rpm).

Cells were harvested by centrifugation at 6000 x g for 30 min at 4 °C. The cell pellet was resuspended in lysis buffer (50 mM Tris-HCl pH 8.0, 500 mM NaCl, 10 mM imidazole, 5% v/v glycerol and 1 mM tris(2-carboxyethyl)phosphine (TCEP) and lysed by sonication. The lysate was clarified by centrifugation at 40,000 g for 30 min, and the supernatant was loaded onto an equilibrated (lysis buffer) immobilised metal affinity chromatography column (IMAC) (Ni-IDA Sepharose, GE healthcare).

The immobilised protein was washed with lysis buffer and eluted with elution buffer (50 mM Tris-HCl pH 8.0, 500 mM NaCl, 300 mM imidazole, 5% v/v glycerol and 1 mM TCEP). The protein-containing fractions were pooled and dialysed overnight at 4 °C in dialysis buffer (25 mM Tris-HCl pH 8.0, 150 mM NaCl, 5% v/v glycerol, and 1 mM TCEP) supplemented with recombinant tobacco etch virus (rTEV) protease for cleavage of the N-terminal 6xHis tag. The protein was then subjected to a second IMAC step to remove His-tagged rTEV protease, cleaved His tag, and uncleaved His-tagged nsp14-10 complex.

The cleaved nsp14-10 complex was concentrated to 1 mL using a 50 kDa MWCO centrifugal concentrator. The protein was then further purified on to a Superdex 200 increase 10/300 GL column equilibrated with SEC buffer (25 mM HEPES pH 7.5, 150 mM NaCl, 5% v/v glycerol, 2 mM TCEP) at 0.8 mL/min. The purified protein was concentrated to 0.5 mg mL^-1^ and was snap frozen in liquid nitrogen for later use. Monomeric nsp14 was purified from the same vector under different induction conditions (overnight growth without adding IPTG).

The presence of the nsp14-10 complex was confirmed via SDS-PAGE, and ESI-Q-TOF (electrospray-ionisation quadrupole time-of-flight) mass spectroscopy (MS).

### Generation of 5′ radiolabelled substrates

10 pmol of single-stranded RNA or DNA (Eurofins MWG Operon, Germany) was incubated with 6.8 pmol γ-32P-dATP (Perkin Elmer), and 10 U T4 PNK (ThermoFisher Scientific) at 37 °C for 1 hour. This solution was passed through a P6 Micro Bio-Spin chromatography column (BioRad) to remove unbound label and diluted accordingly in nuclease-free ultrapure H_2_O.

For double-stranded structures, radiolabelled RNA or DNA was annealed to unlabeled complementary molecules at a 1:1.5 ratio to give a final concentration of 100 nM in 10 mM Tris-HCl (pH 7.5), 50 mM NaCl and 1 mM EDTA by heating the sample to 95 °C for 3 min before cooling to room temperature. A detailed list of all oligonucleotide sequences can be found in Supplementary Table 1.

### Generation of a single nucleotide RNA ladder

10 pmol of 5′ radiolabelled single-stranded RNA was incubated in 50 mM potassium acetate (pH 7.9), 20 mM Tris-acetate (pH 7.9), 10 mM magnesium acetate (pH 7.9) and 1 mM DTT for 2, 5, 10, and 15 min at 95 °C respectively. 5 μL of stop solution (95% formamide, 10 mM EDTA, 0.25% xylene cyanol, 0.25% bromophenol blue) was added to each sample, samples were combined and run on a 20% denaturing polyacrylamide gel (40% solution of 19:1 acrylamide:bis-acrylamide, BioRad) and 7 M urea (Sigma Aldrich)) in 1 x TBE (Tris-borate EDTA) buffer. Electrophoresis was carried out at 525 V for 1.5 hours; gels were subsequently fixed for 60 min in fixing solution (50% methanol, 10% acetic acid), and dried at 80 °C for two hours under a vacuum. The dried gels were exposed to a Kodak phosphorimager screen and scanned using a Typhoon 9400 instrument (GE).

### Nuclease assays

Nuclease assays were carried out in 10 μL reactions containing 25 mM HEPES-KOH, pH 7.5, 50 mM NaCl, 5 mM MgCl_2_, 5% glycerol, 1.0 mM DTT, 10 nM RNA or DNA substrate and serially diluted two-fold nsp14-nsp10 complex or nsp14 from 500 nM. Reactions were incubated at 37 °C for 45 min, and quenched by the addition of 5 μL stop solution (95% formamide (v/v), 10 mM EDTA, 0.25% xylene cyanol, 0.25% bromophenol blue) and boiling at 95 °C for 3 min.

Reactions were then analysed by 20% denaturing polyacrylamide gel electrophoresis (40% solution of 19:1 acrylamide:bis-acrylamide, BioRad) and 7 M urea (Sigma Aldrich)) in 1 x TBE (Tris-borate EDTA) buffer. Electrophoresis was carried out at 525 V for 1.5 hours; gels were subsequently fixed for 60 min in fixing solution (50% methanol, 10% acetic acid), and dried at 80 °C for two hours under a vacuum. The dried gels were exposed to a Kodak phosphorimager screen and scanned using a Typhoon 9400 instrument (GE).

Where applicable, gel-based nuclease assays were quantified by analysing the product formation (digestion) of the radiolabelled RNA substrate(s). Gel images collected on a Typhoon scanner were analysed in Image J to determine the amount of substrate remaining in the well (undigested) in comparison to the amount of substrate that entered the gel (product) reported as a percent digested. This was plotted versus protein concentration to indicate digestion rate. Error is standard error. Three or more gels were analysed for each substrate.

### Nuclease inhibition assays

Inhibitor exonuclease assays were carried out in 10 μL volumes containing 25 mM HEPES-KOH, pH 7.5, 50 mM NaCl, 5 mM MgCl_2_, 5% glycerol, 1.0 mM DTT and 100 nM nsp14-10. Inhibitors were solubilised in DMSO and diluted immediately prior to the reaction in the above buffer. Inhibitors were serially diluted two-fold from 100 μM and the DMSO concentration was keep constant at 0.1% in the final reaction. Inhibitors were incubated for 10 min at room temperature with the nsp14-10 complex, before the reactions were initiated by the addition of RNA substrate (10 nM), incubated at 37 °C for 45 min, then quenched by the addition of 5 μL stop solution (95% formamide, 10 mM EDTA, 0.25% xylene cyanol, 0.25% bromophenol blue) and boiling at 95 °C for 3 min. Reactions were analysed by 20% denaturing PAGE, fixed, dried, then imaged as described above.

### Molecular docking

The structure of the best available SARS-CoV-1 nsp14-nsp10 entry in the Protein Data Bank (PDB), PDB ID 5NFY with a resolution of 3.38 Å^10^, were aligned with another SARS-CoV-1 nsp14-nsp10 structure (PDB: 5C8U, resolution 3.40 Å^8^) to incorporate the missing active site Mg^2+^ in 5NFY from that in 5C8U. The coordinates were then translated such that the centre of mass was closer to the origin, (0,0,0), and prepared for docking with AutoDockTools. Blind docking was carried out using AutoDock Vina using a grid box of 64 Å x 70 Å x 126 Å centred on (-0.476, 5.300, 8.298), encompassing the entire protein surface. A second set of more focused dockings was performed with AutoDock Vina focused on the active site, with the grid box specified to be 30 Å x 24 Å x 24 Å centred on (-5.196, -9.373, -7.837). Predicted AutoDock Vina affinities of each docked binding mode for each ligand were extracted, along with the distance from the centre of the binding mode to the catalytic Mg^2+^, and to the centre of mass of GpppA in the MTase binding site. Docked poses were analysed using PyMOL and rendered with ChimeraX.

## ACNOWLEDGEMENTS

Our work was supported by the University of Oxford COVID-19 Research Response Fund. Related work on nucleases in the laboratory of PM is supported by Cancer Research UK, MRC, and AstraZeneca. Related work in CJS and OG laboratories is supported by Cancer Research UK, and the Wellcome Trust.

## AUTHOR CONTRIBUTIONS

OG, CJS and PM conceived the project and helped design the experiments. YY designed and executed the protein purification protocols and MS analysis. YY and AC-M purified the proteins. HTB and SB performed the biochemical analysis of nsp14-nsp10, with technical help from LS. MB performed the organic synthesis. MR performed the docking studies under the guidance of GMM. PJM wrote the paper with help from all the authors.

---

**Suppl. Figure 1:**
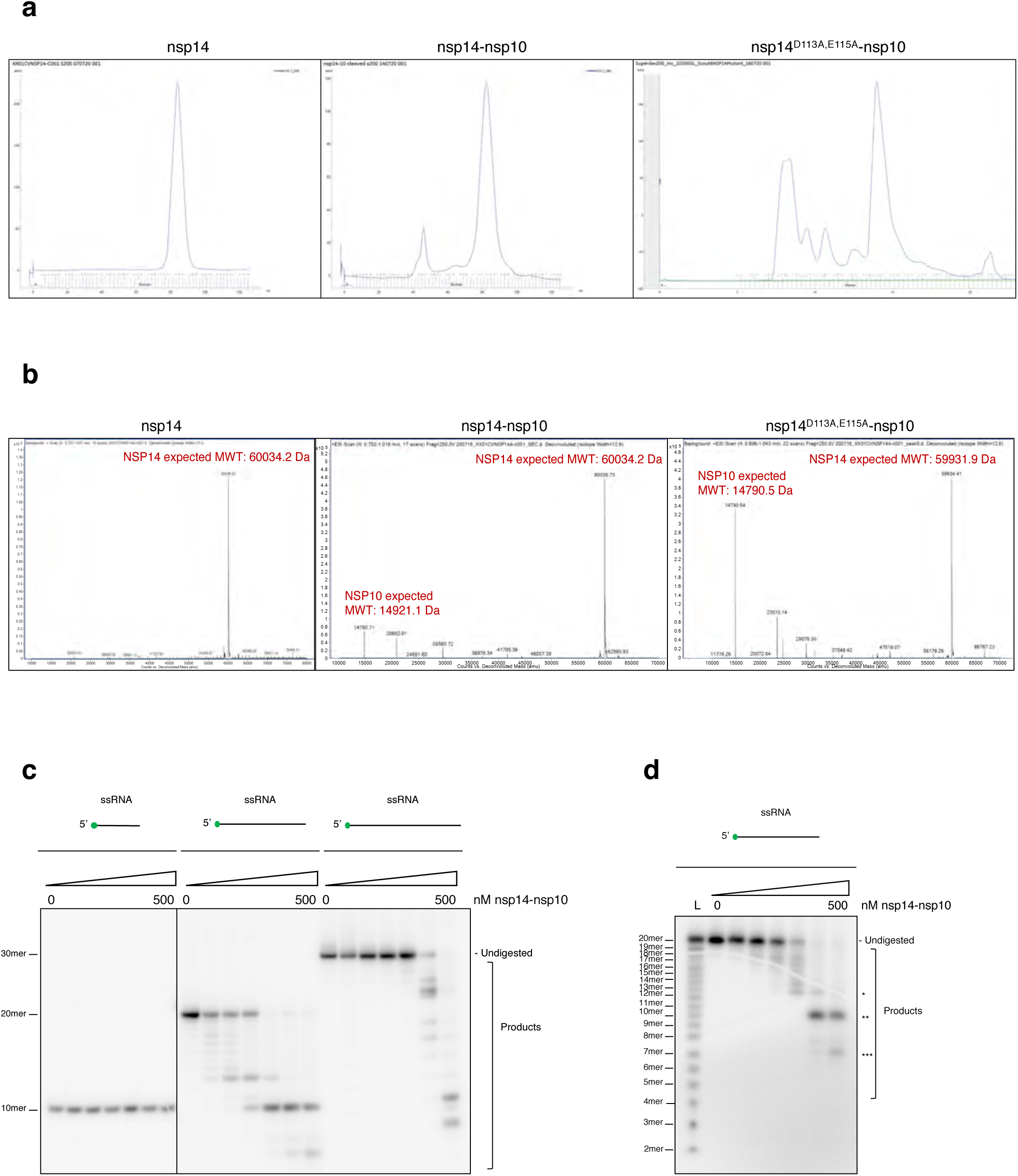
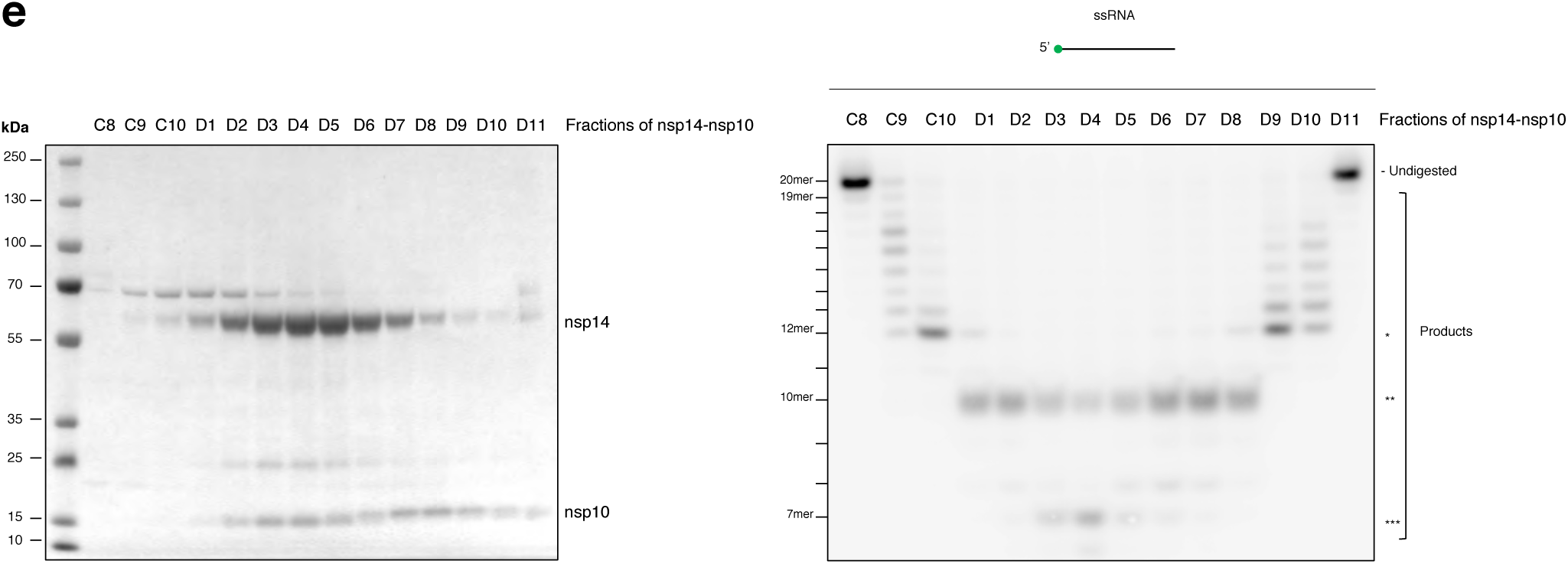
Nsp14-nsp10 is an RNA nuclease with a complex digestion pattern. (a) Size exclusion chromatogram for nsp14 alone, wild-type nsp14-nsp10 (designated nsp14-nsp10), and a control ‘nuclease-dead’ complex bearing alanine substitutions at residues D113 and E115 (nsp14^D113A,E115A^-nsp10). Traces of Superdex 200 16/60 runs are shown for Nsp14 alone and wild-type nsp14-nsp10 and trace of Superdex increase 10/300 run for the ‘nuclease-dead’ complex. Wild-type nsp14-nsp10 complex elutes in a single peak at ∼82.5 mL, Nsp14 alone at ∼84.3 mL and ‘nuclease-dead’ complex at ∼14.6 mL . SDS-PAGE analysis of the fraction across the peak confirms the presence of both nsp14 and nsp10 in fractions D1–D8 (see Suppl. Fig. 1e). (b) Intact mass spectrometry analysis of nsp14 alone, wild-type nsp14-nsp10 (designated nsp14-nsp10) and a control ‘nuclease-dead’ complex bearing substitutions at residues D113 and E115 (nsp14^D113A,E115A^-nsp10). LC-MS chromatogram shows the observed mass of all proteins is within range of the calculated mass. (c) Nsp14-nsp10 digests 20-mer and a 30-mer ssRNA substrates, but not a 10-mer ssRNA. (d) Nsp14-nsp10 is an RNA nuclease manifests a complex digestion pattern, digesting from the 3’ end in a single-nucleotide fashion until the 8^th^ ribonucleotide, then cleaving at the 10^th^ and 13^th^ ribonucleotide. A single nucleotide ladder was used to determine the size of the released products. (e) SDS-PAGE of the fractions before, during and after the peak used from gel filtration for wild-type nsp14-nsp10. The elution peak (D1–D8) coincides precisely with the characteristic RNAse activity that we observe in our activity gel. The predicted molecular weight is 59 463 Da for nsp14 and 15 281 Da for nsp10. Increasing concentrations of protein (as indicated) were incubated with substrate at 37°C for 45 min, reactions were subsequently analysed by 20% denaturing PAGE to visualize product formation. The size of products was determined as shown in Suppl. Fig 1D. Main products are labelled *, ** and *** corresponding to 12-mer, 10-mer and 7-mer respectively. All oligos used are indicated in Suppl. table 1A and B. For Suppl. Figs. 1D and E, oligo 2 was used.

**Suppl. Figure 2:**
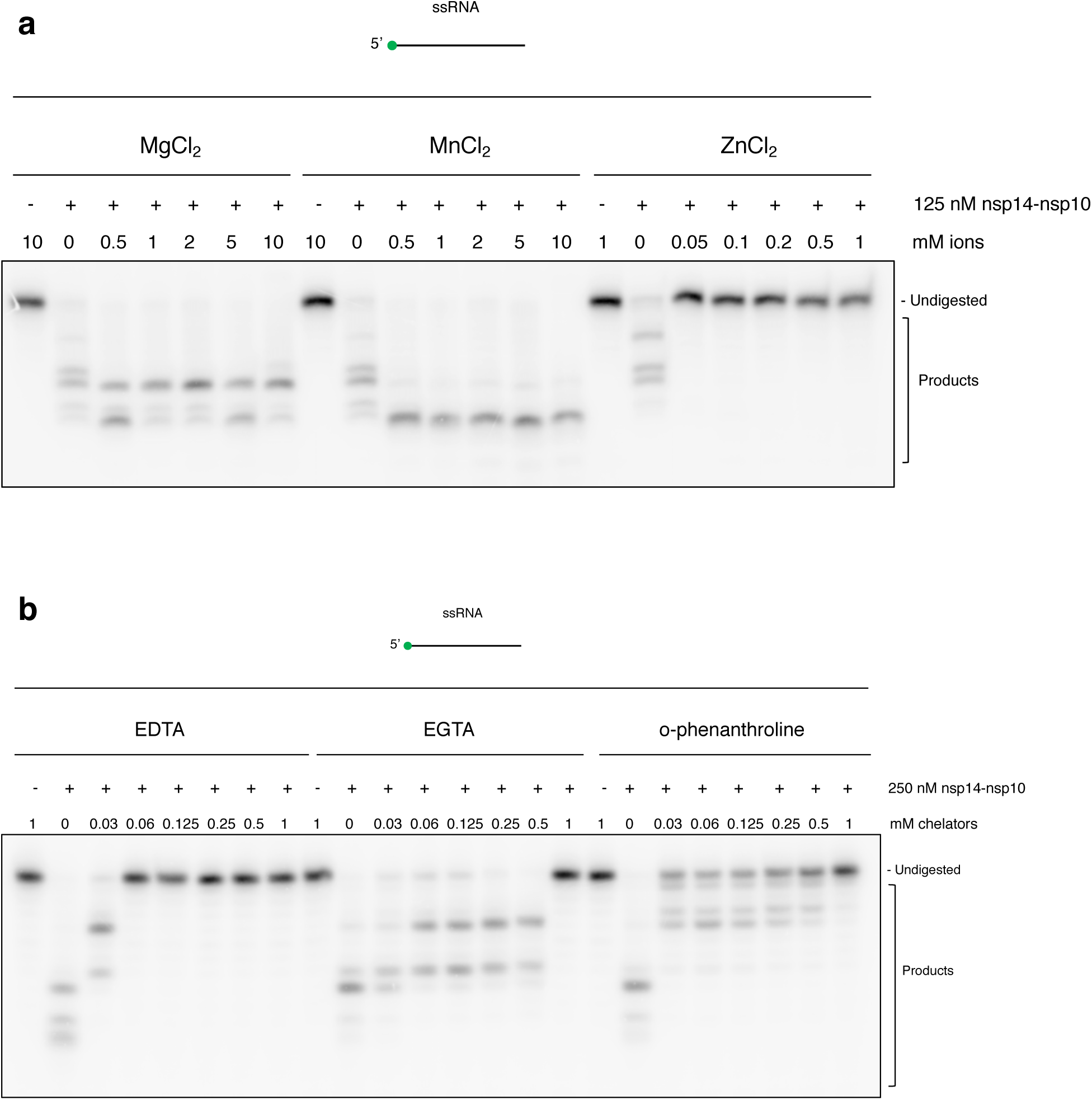
Nsp14-nsp10 is metal-ion dependent RNA nuclease. (a) Nsp14-nsp10 activity is enhanced by the addition of both MgCl2 and MnCl2, but is inhibited by ZnCl2. We determined 5 mM MgCl2 as optimal for activity. (b) Consistent with its requirement for metal ions, nsp14-nsp10 activity is inhibited by the addition of metal ion chelators EDTA, EGTA, and ο-phenanthroline, with a particular sensitivity towards EDTA. 125 nM or 250 nM of protein was incubated with increasing concentrations of ions or chelators (as indicated) on ice for 15 min before adding substrate and incubating the reaction at 37°C for 45 min. Reactions were subsequently analysed by 20% denaturing PAGE to visualize product formation. For Suppl. Figs. 2A and B, oligo 2 was used.

**Suppl. Figure 3:**
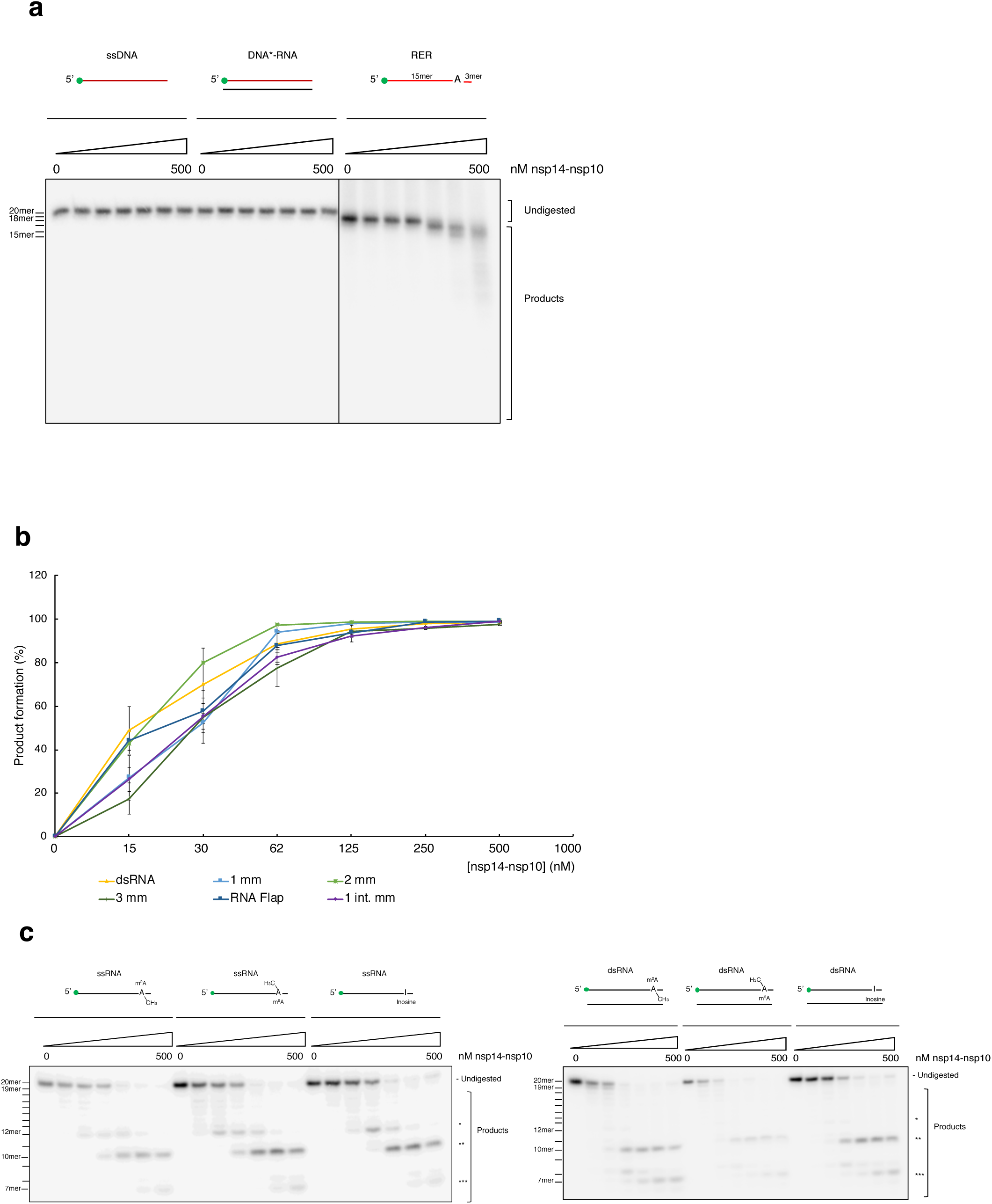
Nsp14-nsp10 activity is specific to RNA only and is able to process common chemical modifications on RNA. (a) Nsp14-nsp10 has no discernible nuclease activity on ssDNA or the DNA strand of an DNA:RNA hybrid. Interestingly, the complex is able to incise around an embedded ribonucleotide in a ssDNA oligo at concentrations above 125 nM. (b) Product formation (%) was quantified for Fig. 3 B comparing nsp14-nsp10 nuclease activity on dsRNA and RNA substrates containing termini and internal mismatches as outlined in Methods and Materials. All data are shown as mean ± s.e.m, and at least three biological replicates were used for each substrate. Nsp14-nsp10 shows no preference for mismatched oligonucleotides compared to dsRNA. mm: mismatch int. mm: internal mismatch (c) Nsp14-nsp10 is able process common chemical modifications of RNA, namely 2-methyladenine (m^2^A), 6-methyladenine (m^6^A) and inosine (I), both ssRNA and dsRNA with no apparent change compared to ssRNA or dsRNA. Increasing concentrations of protein (as indicated) were incubated with substrate at 37°C for 45 min, reactions were subsequently analysed by 20% denaturing PAGE to visualize product formation. The size of products was determined as shown in Suppl. Fig 1D. Main products are labelled *, ** and *** corresponding to 12-mer, 10-mer and 7-mer respectively. Oligomer substrates are in Suppl. Table 1A and B.

**Suppl. Figure 4:**
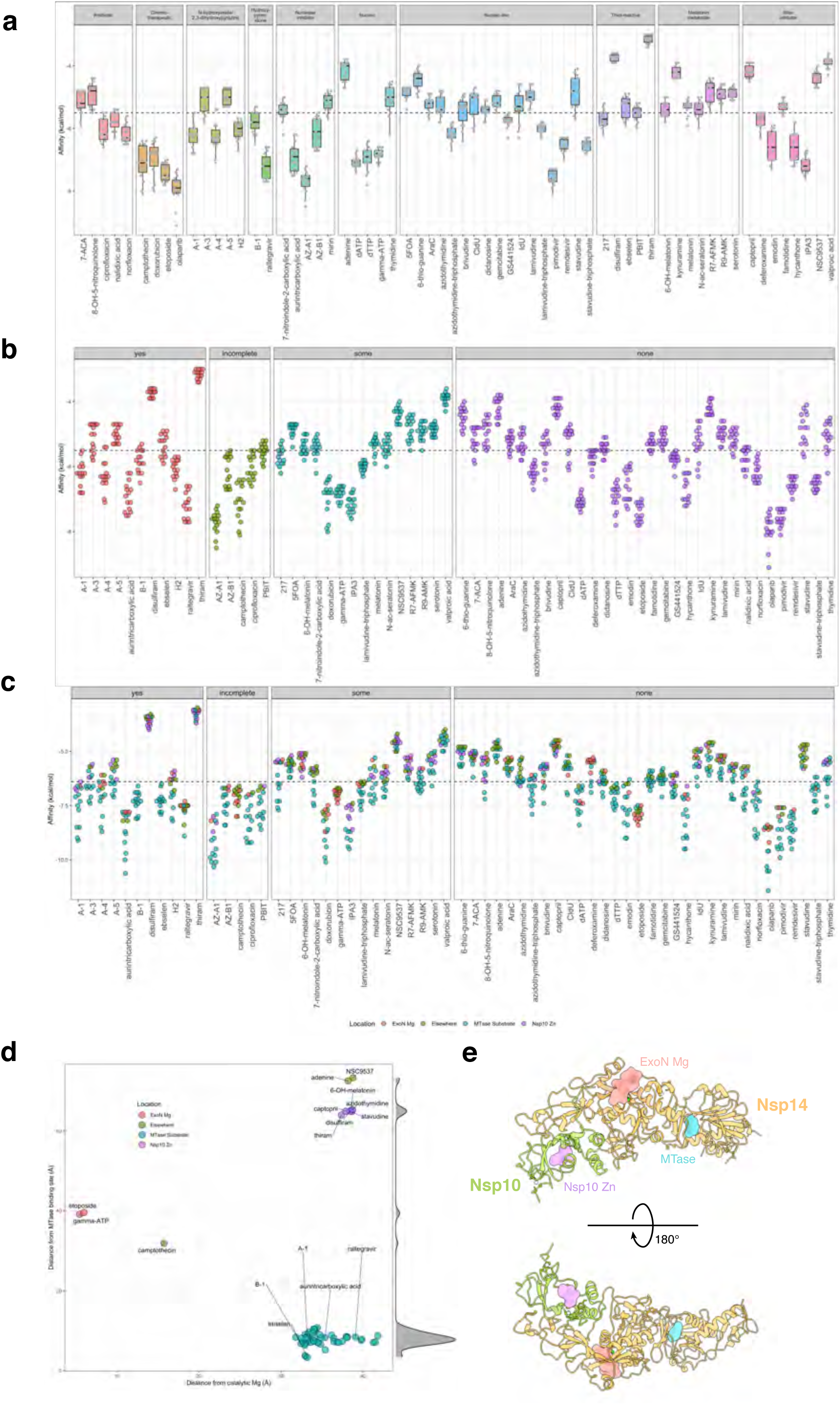
Molecular docking of SARS-CoV nsp14-nsp10 using Autodock. (a)&(b) Nsp14-nsp10 was docked with compounds within a grid box encompassing a surface focussed on the active. The calculated affinities of all binding modes are shown for each compound. Compounds are grouped based on their function groups (a), and by predicted inhibition activity (b); the dashed line describes the median of affinities for all compounds. (c) Nsp14-nsp10 was docked with compounds within a grid box encompassing the full surface. The affinities of binding modes were calculated for each compound and grouped based on predicted ???activities. The colouring reflects the location of binding mode on the structure of nsp14-nsp10; dashed line describes the median of affinities in this cohort. (d) The positions of highest-affinity docked poses as a function of the distance from the active site ExoN Mg^2+^ centre and the distance from the MTase substrate (GpppA) binding site. (e) Three representatives of predicted inhibitor binding locations on nsp14-nsp10 are shown: ^+^ (i) nsp-10 Zn site (disulfiram, purple), ExoN Mg^2)^ (etoposide, salmon), and the MTase substrate binding site (A-1, cyan).

**Suppl. Figure 5:**
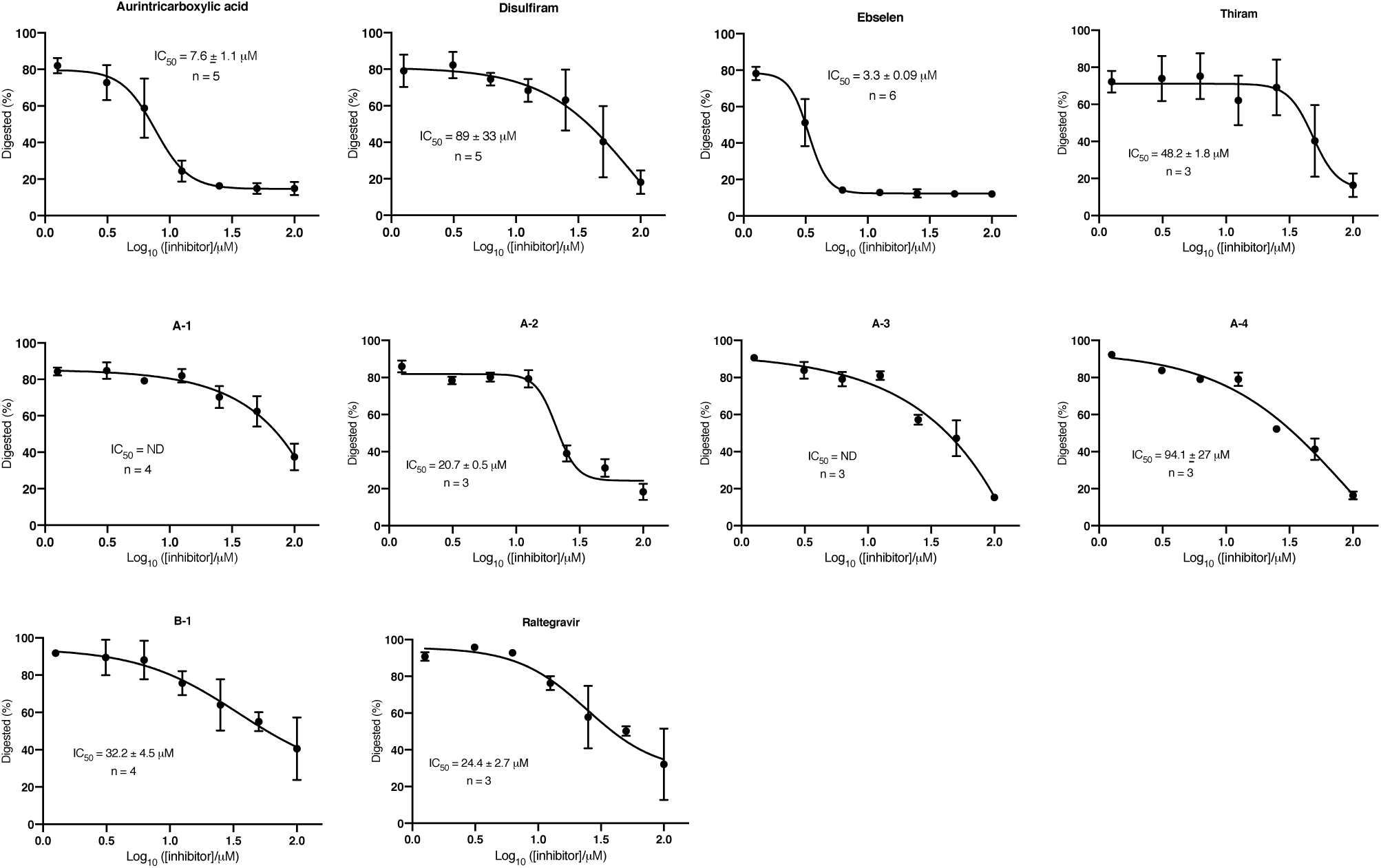
Inhibition profile curves generated from gel-based nuclease assay data with the indicated compounds. Inhibition profile curves were calculated by quantification of gel-based nuclease assay data as outlined in Methods and Materials. A decrease in the ‘% digested’ of the substrate represents an increase in inhibition. Data were plotted against log10 of [inhibitor] and dose response curves were generated using non-linear regression. Where possible IC50 values were obtained. All data are shown as mean ± sem; t least three biological replicates were used for each compound. In some cases an IC50 value was unable to be calculated, either due to incomplete inhibition, or the inability to fit a sigmoidal curve to the data.

**Suppl. Figure 6:**
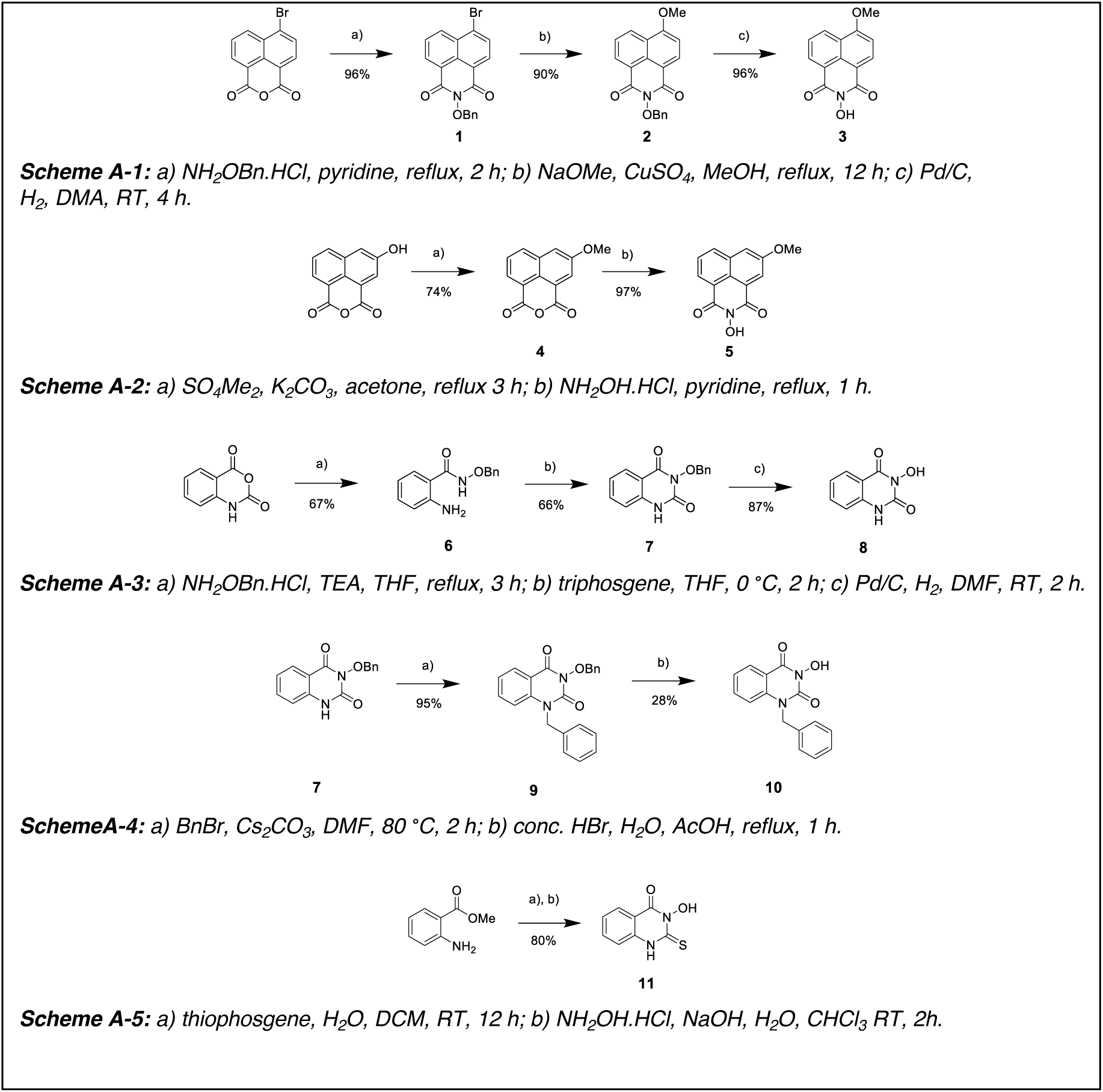
Synthesis scheme for N-hydroxyimide compounds A-1 to A-5. See Suppl. Materials and Methods for detailed synthesis protocols.

**Suppl. Figure 7:**
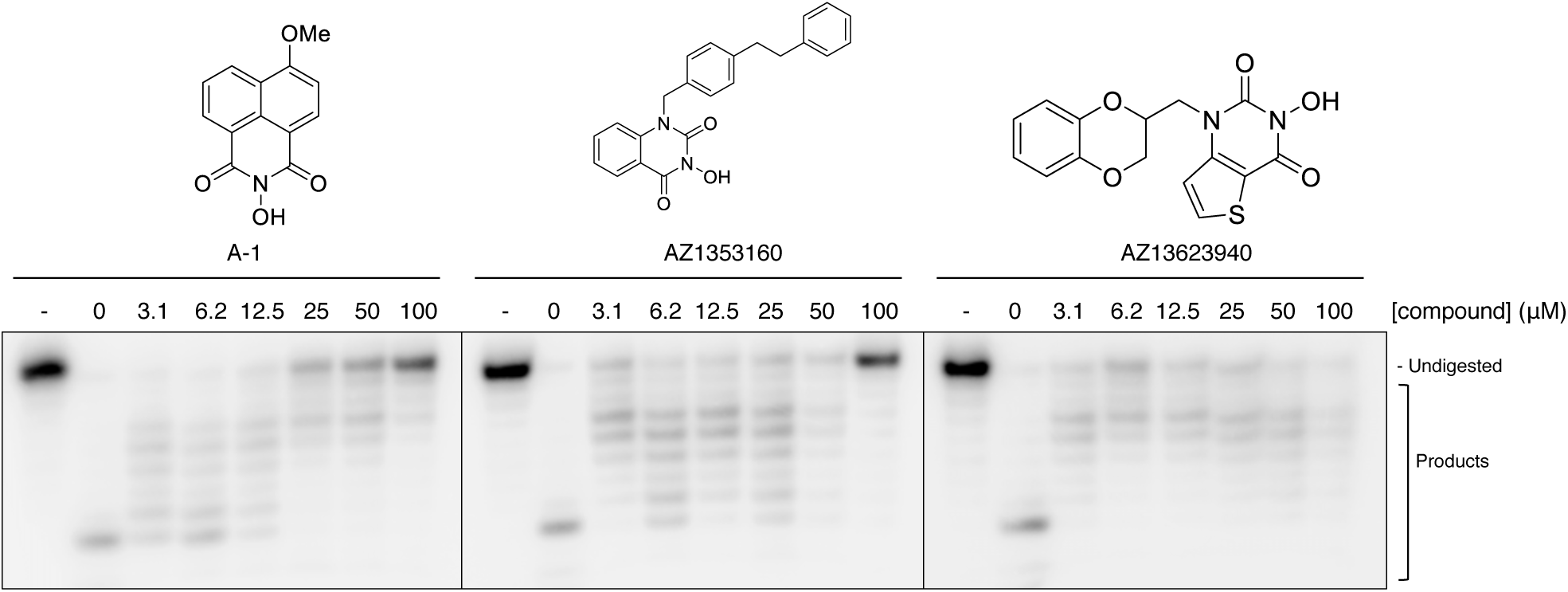
The *N*-hydroxyimide-based compounds, A-1, AZ1353160 and AZ13623940 (AZ-B1), exhibit limited inhibition of nsp14-nsp10. Increasing concentrations (as indicated, in μM) of indicated compounds were incubated with 100 nM nsp14-nsp10 for 10 minutes at room temperature, before starting a standard nuclease assay reaction by the addition of ssRNA, incubating at 37°C for 45 min. Reaction products were analysed by 20% denaturing PAGE. A decrease in the generation of nucleolytic reaction products and a concomitant increase in undigested substrate indicates inhibition of nuclease activity at increasing inhibitor concentrations. - indicates no enzyme.

**Suppl. Figure 8:**
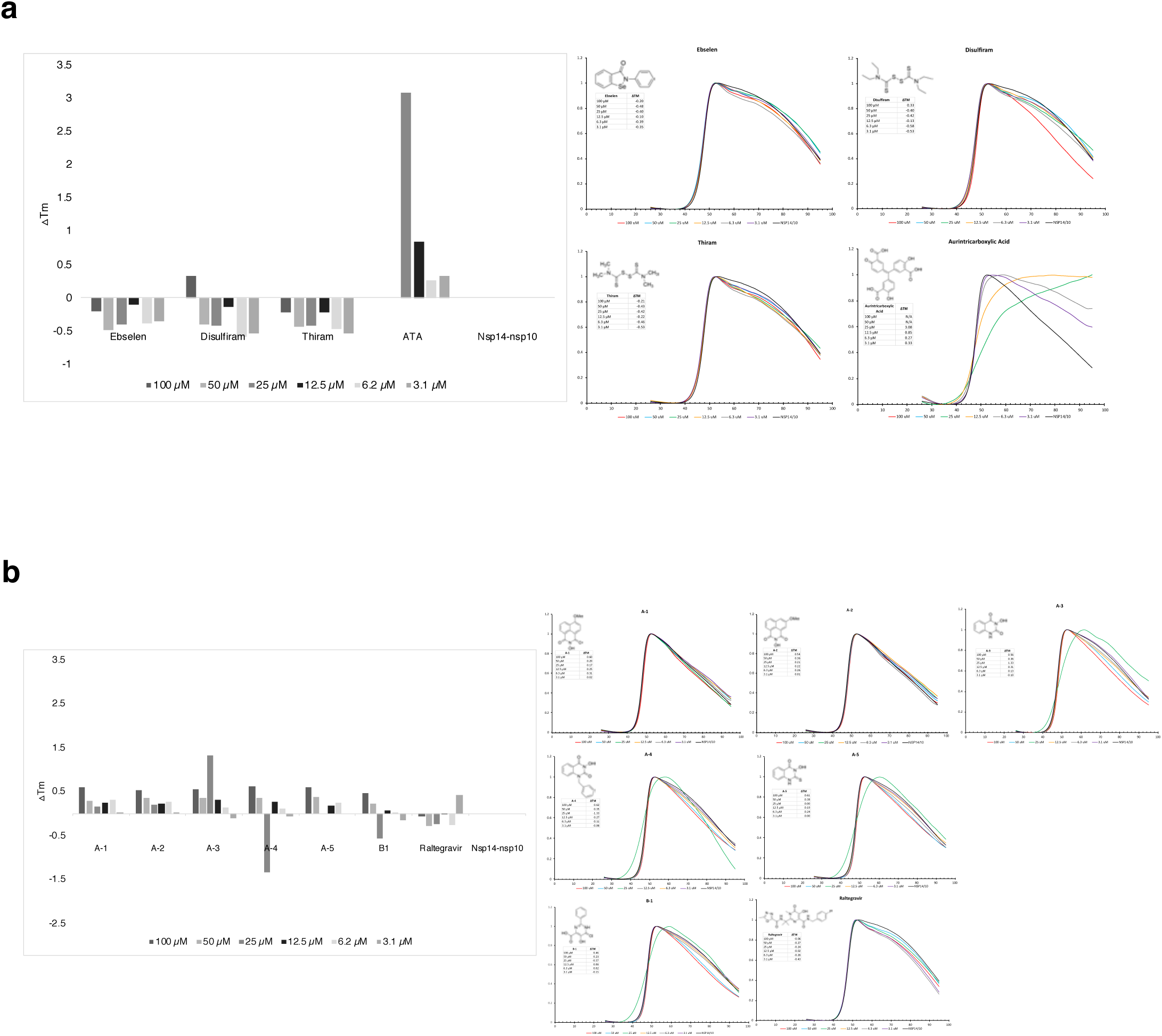
Differential scanning spectrometry (DSF) of nsp14-nsp10 with drug and drug-like compounds and *N*-hydroxyimide and hydroxypyrimidinone containing compounds. (a) ΔTm of nsp14-nsp10 with drugs and drug-like compounds. Of all the compounds tested, only aurintricarboxylic acid (ATA), a known pan-nuclease inhibitor appears to aggregate the protein at high concentration (100 µM and 50 µM). (b) ΔTm of nsp14-nsp10 with *N*-hydroxyimide compounds Samples were heated from 25–95°C in 1°C per minute increments. Fluorescence intensities were plotted as a function of temperature to generate a sigmoidal curve with the inflection point of the transition curve indicating the melting temperature (Tm) of the protein. The Tm of nsp14-nsp10 is around 47°C. The ΔTm for each of the compound dilutions are shown as inset table. See Suppl. Materials and Methods for more detail.

**Suppl. Table 1:**
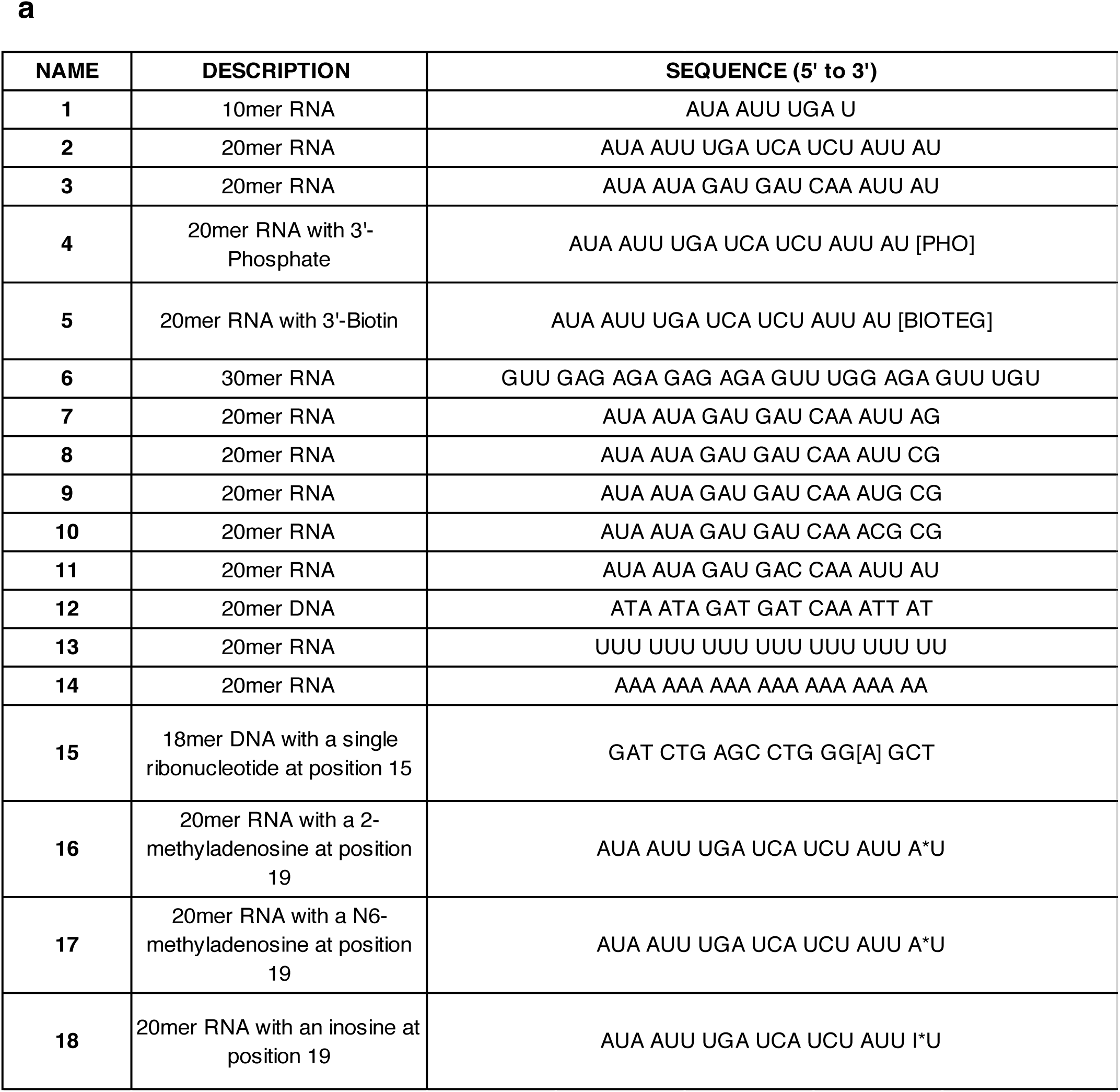
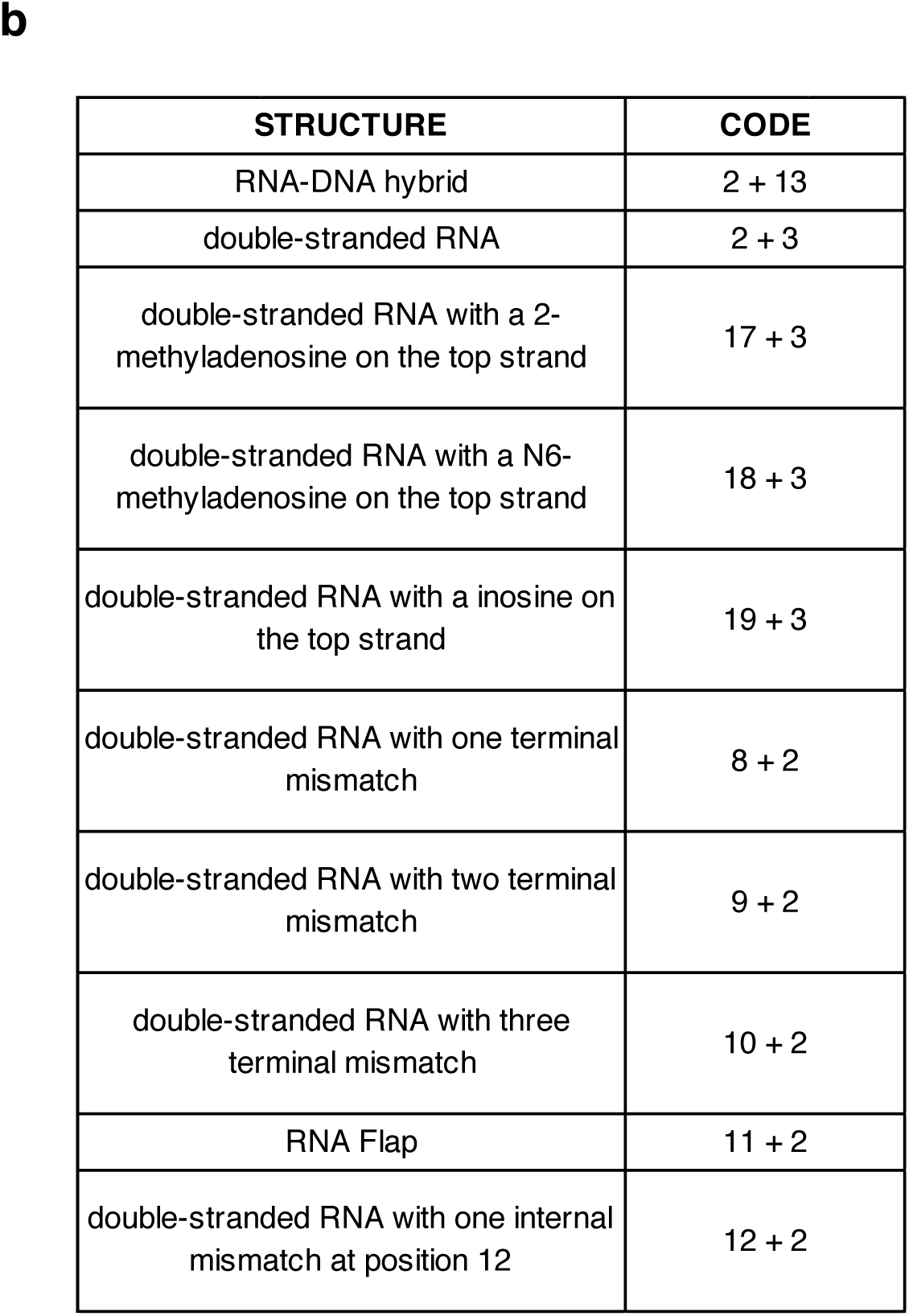
List of RNA and DNA oligonucleotide sequences used to generate simple and complex RNA- and DNA-substrates. (a) Each oligonucleotide sequence is numbered and a general description included. (b) To generate complex RNA and DNA structures, single-stranded oligos were annealed as designated by the numbers using the protocol described in the Materials and Methods.

**Supplementary Table 2.**
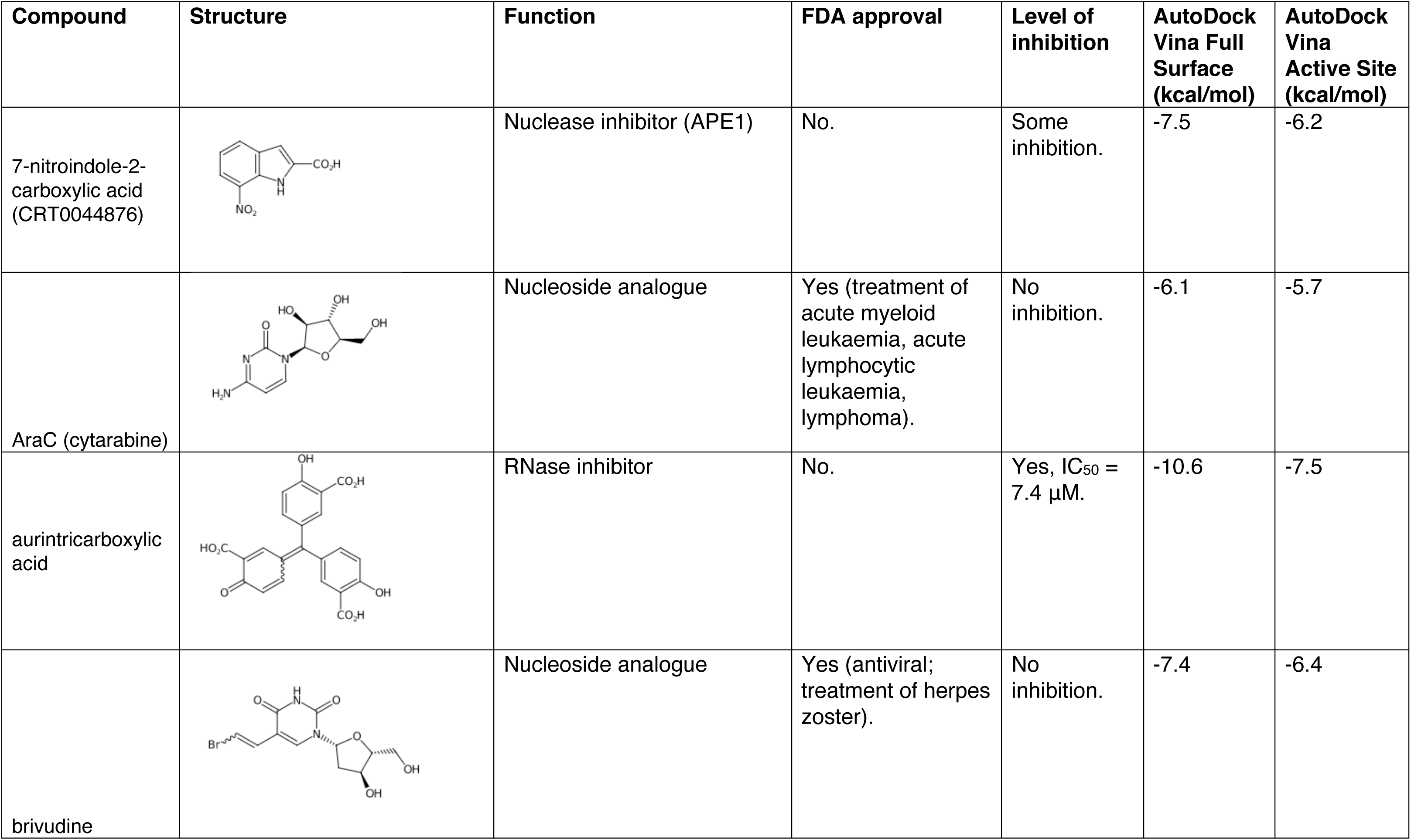
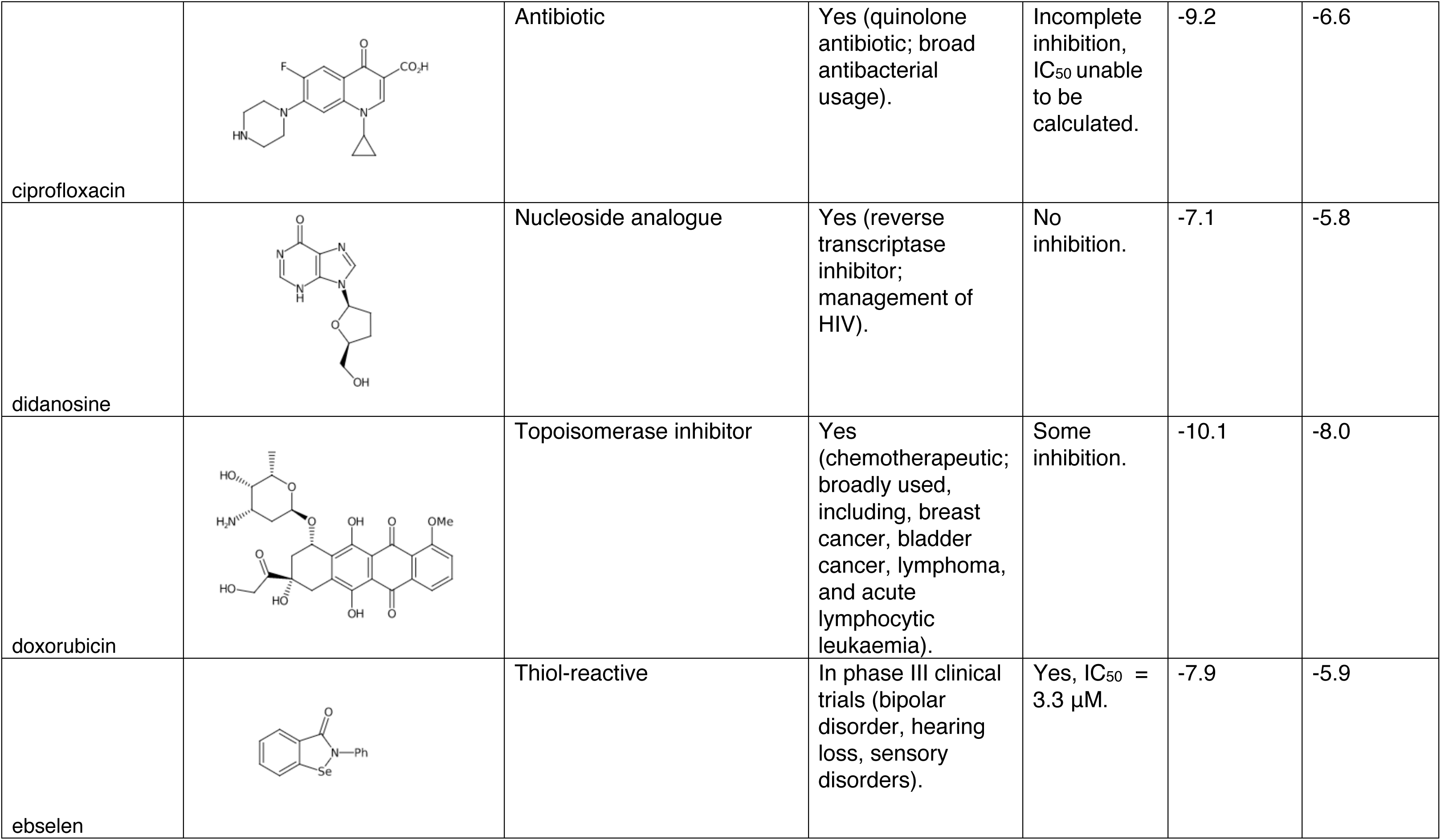
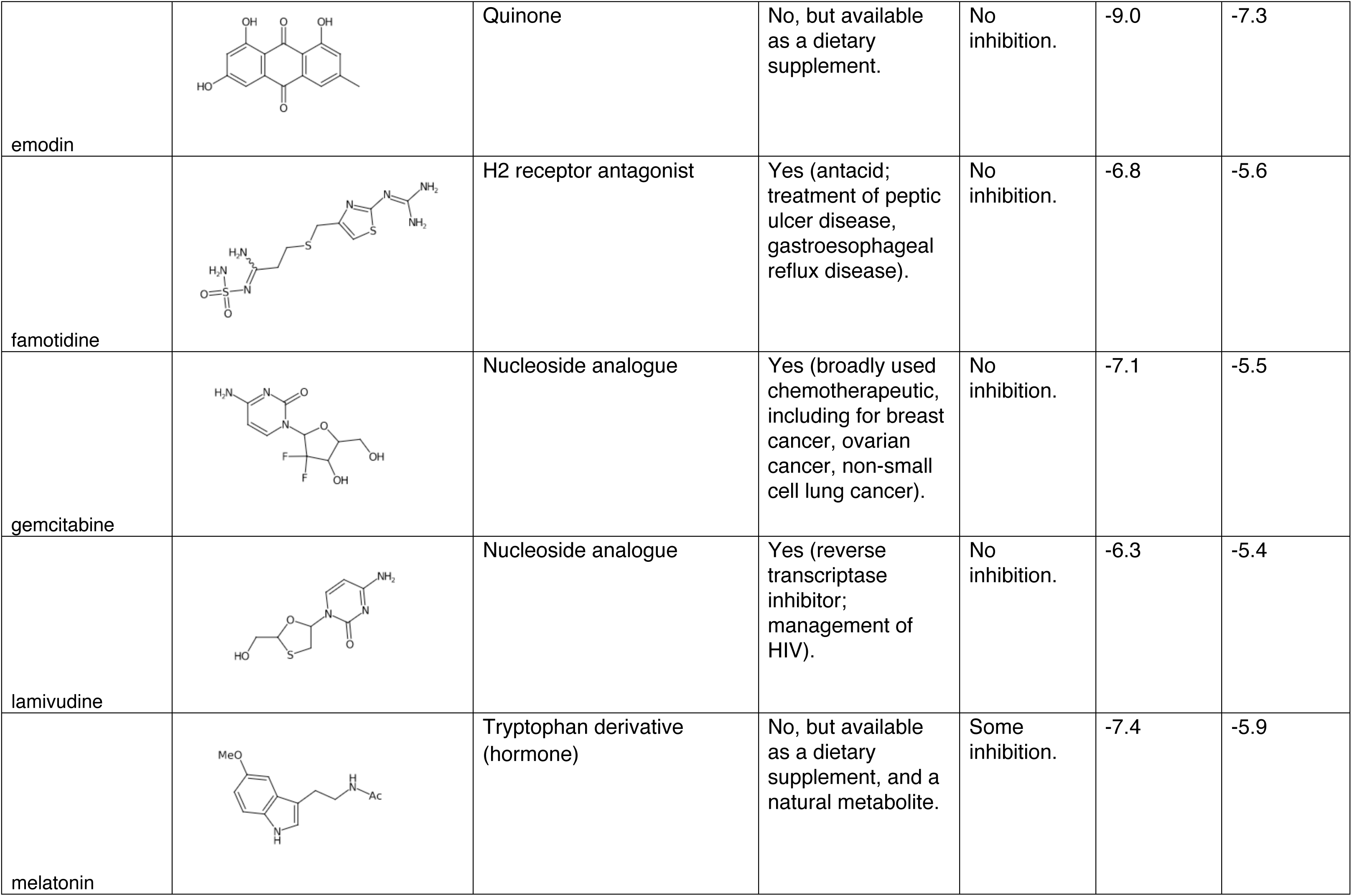
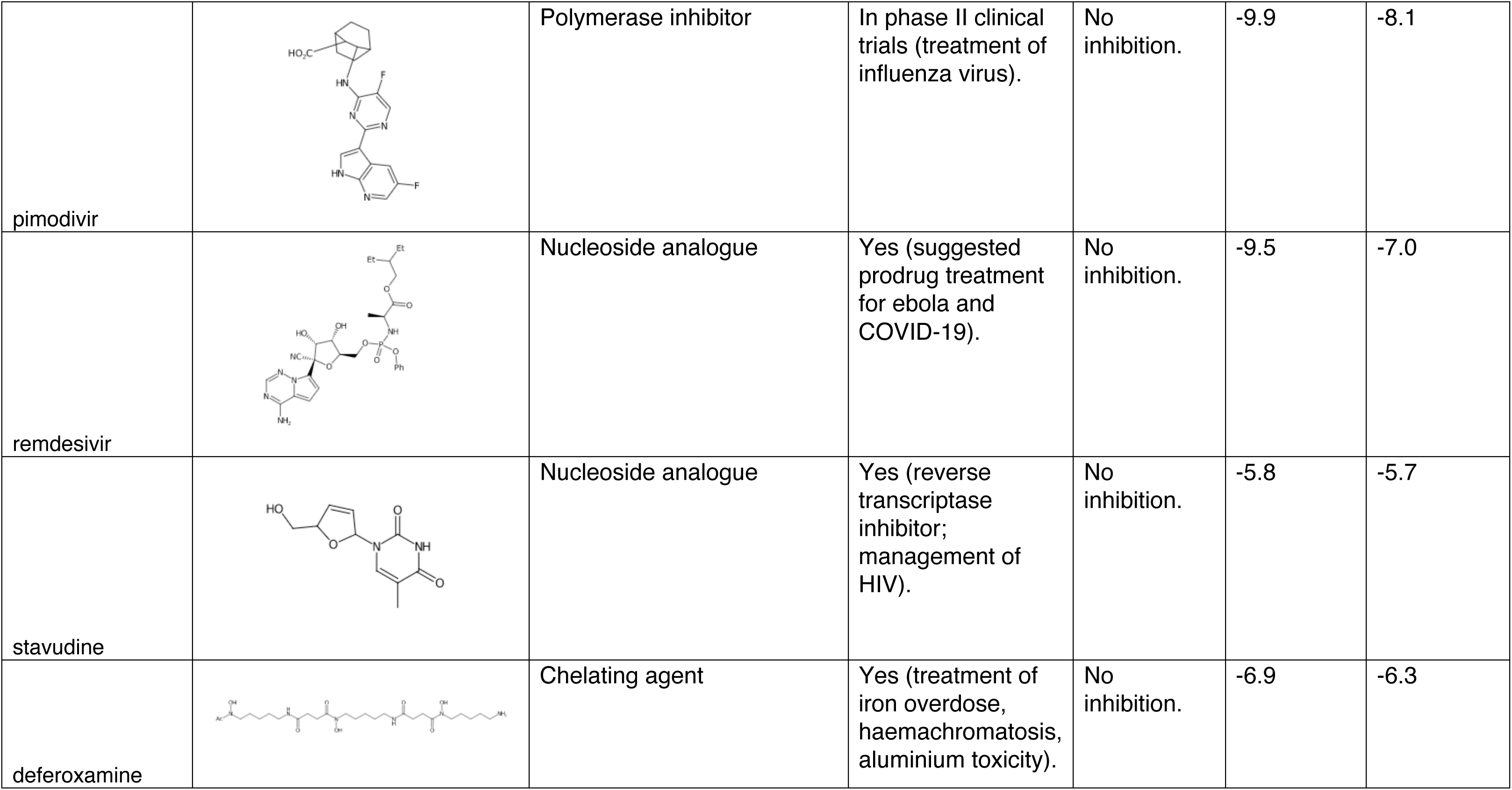
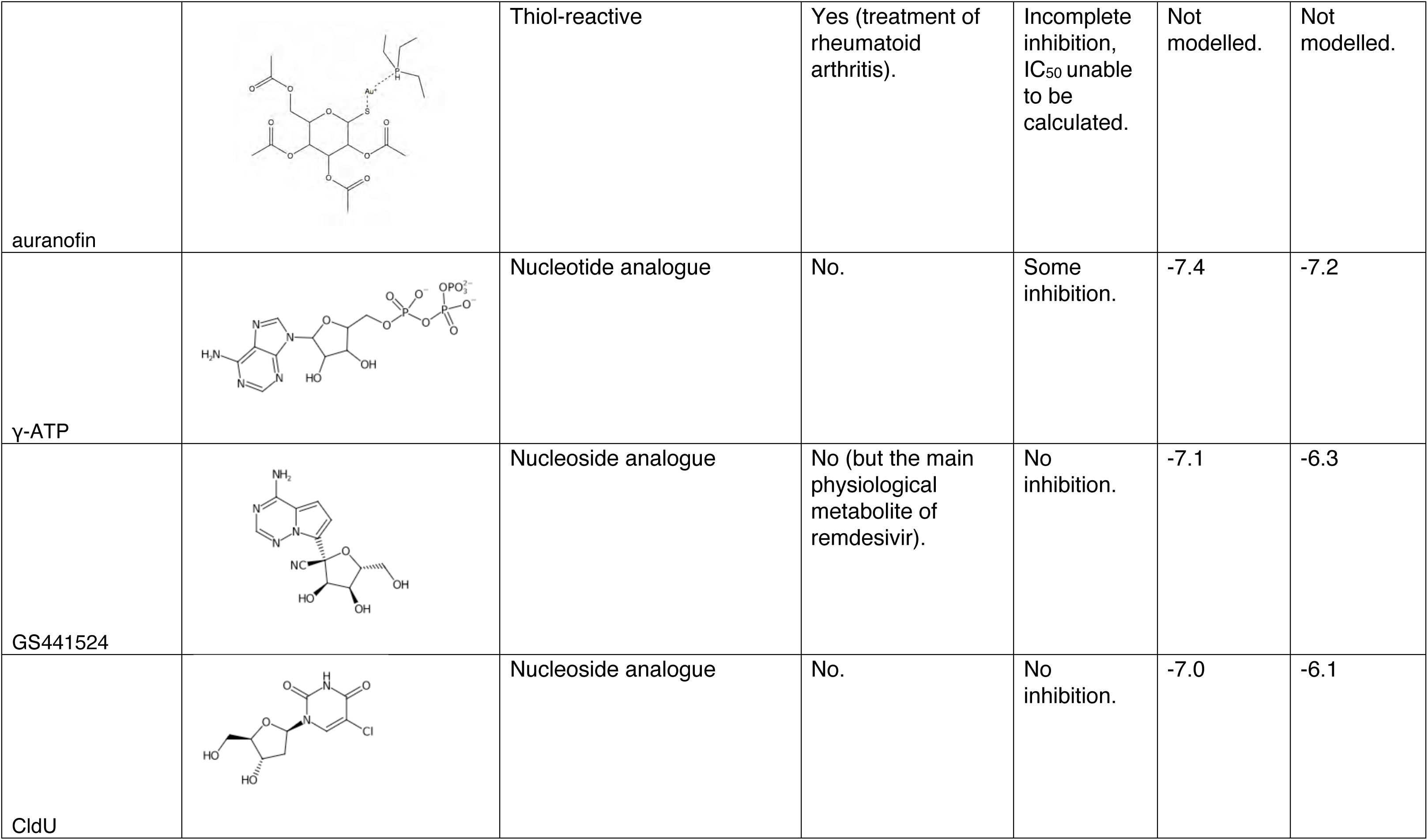
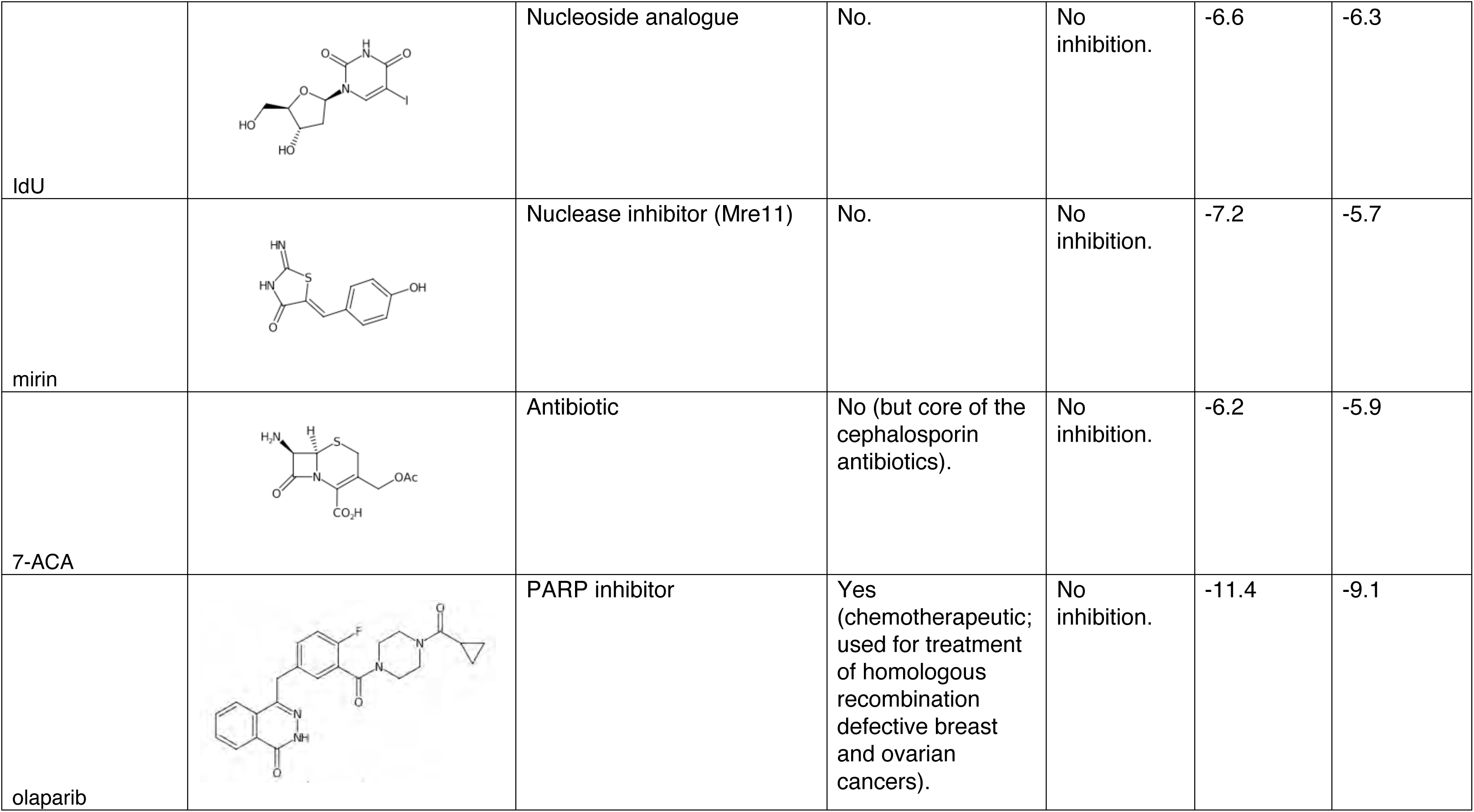
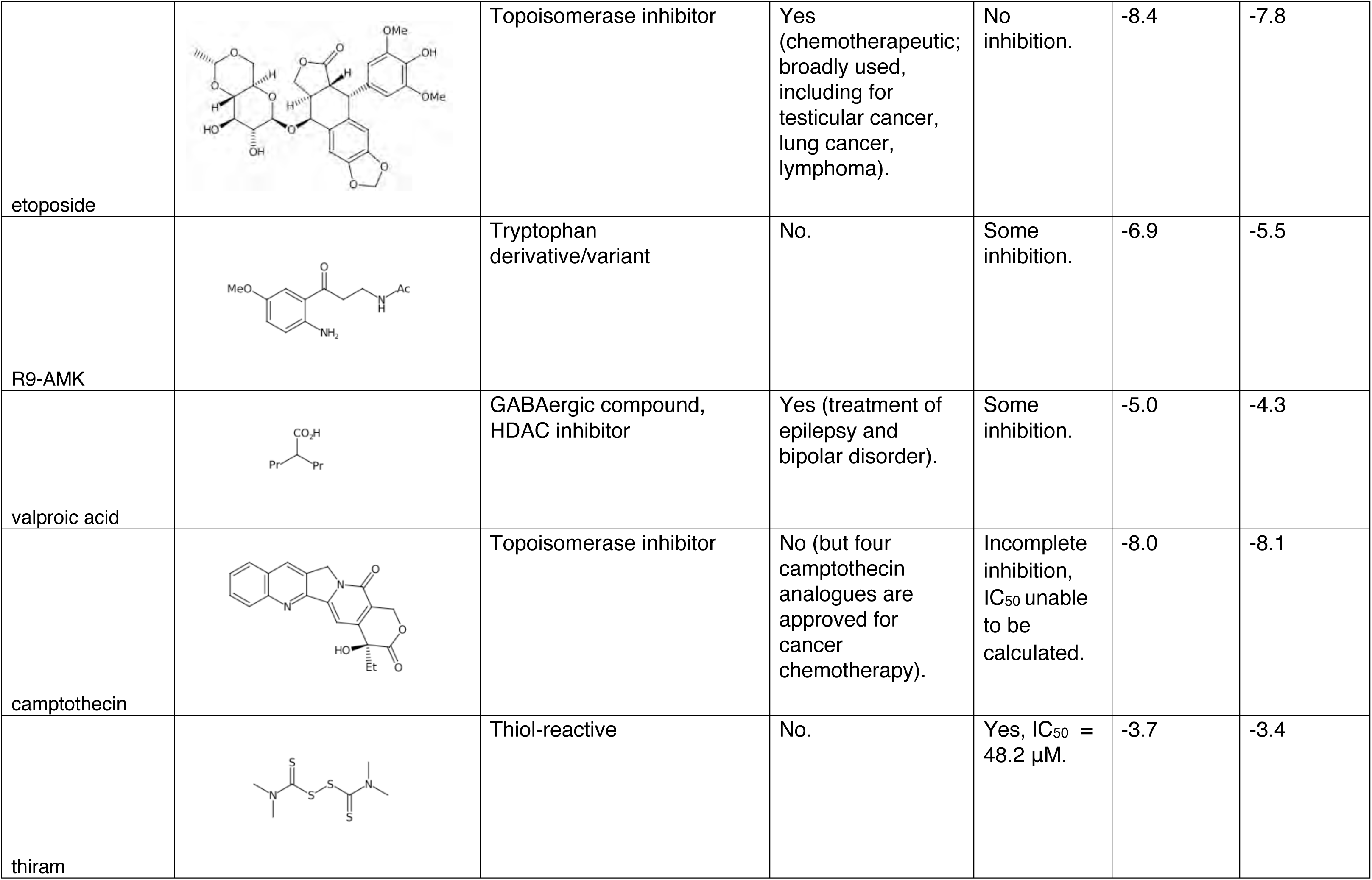
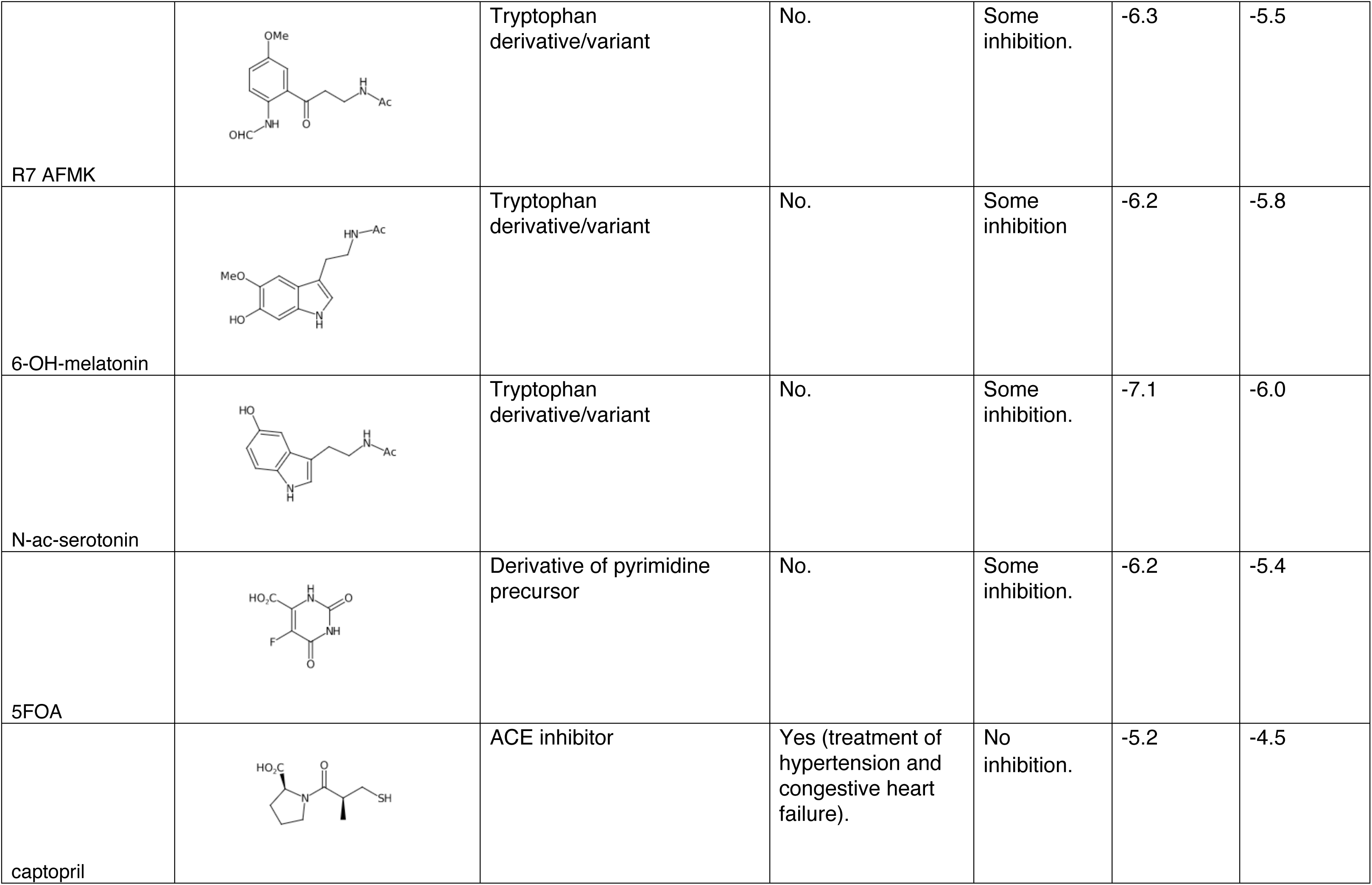
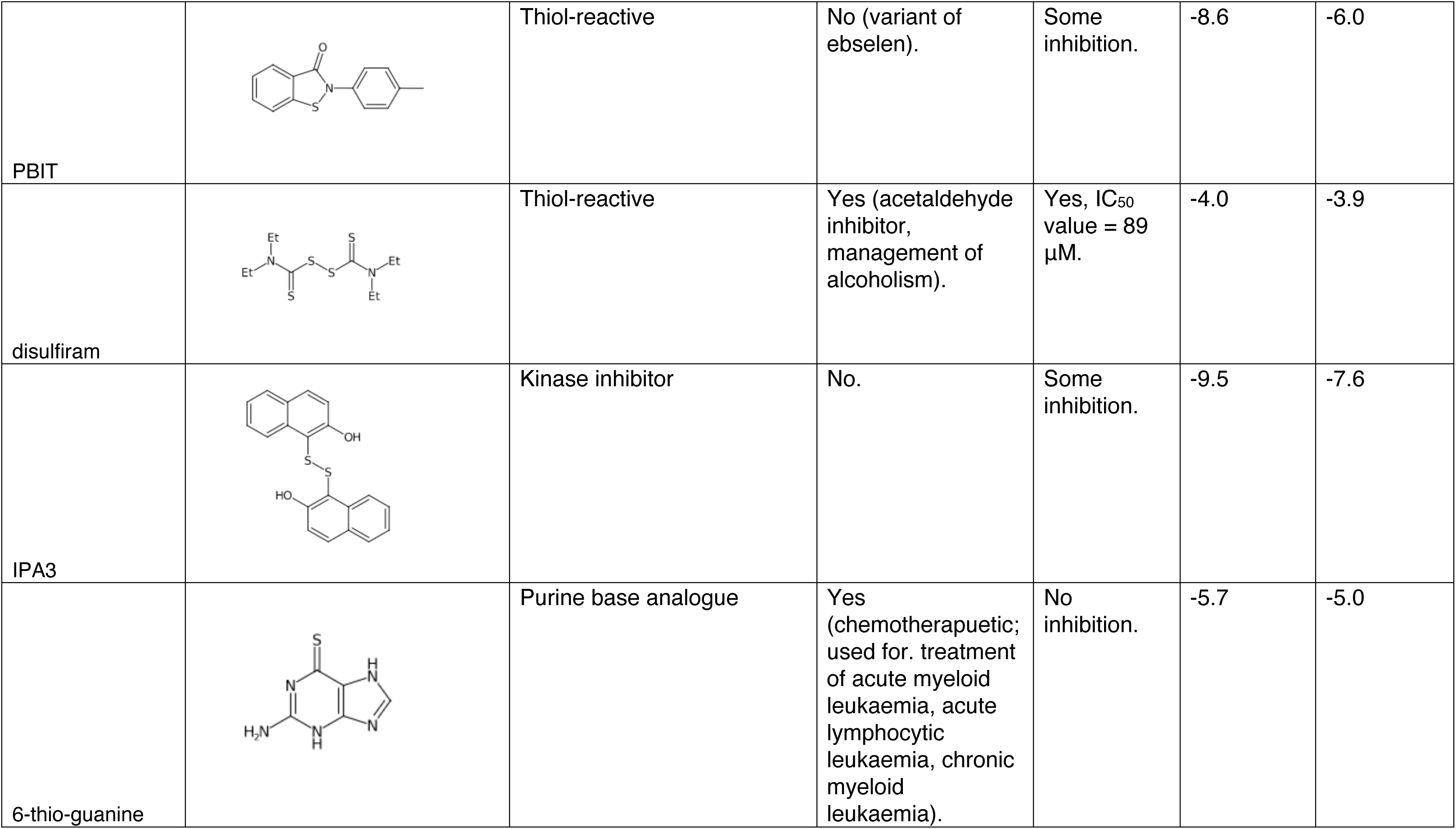
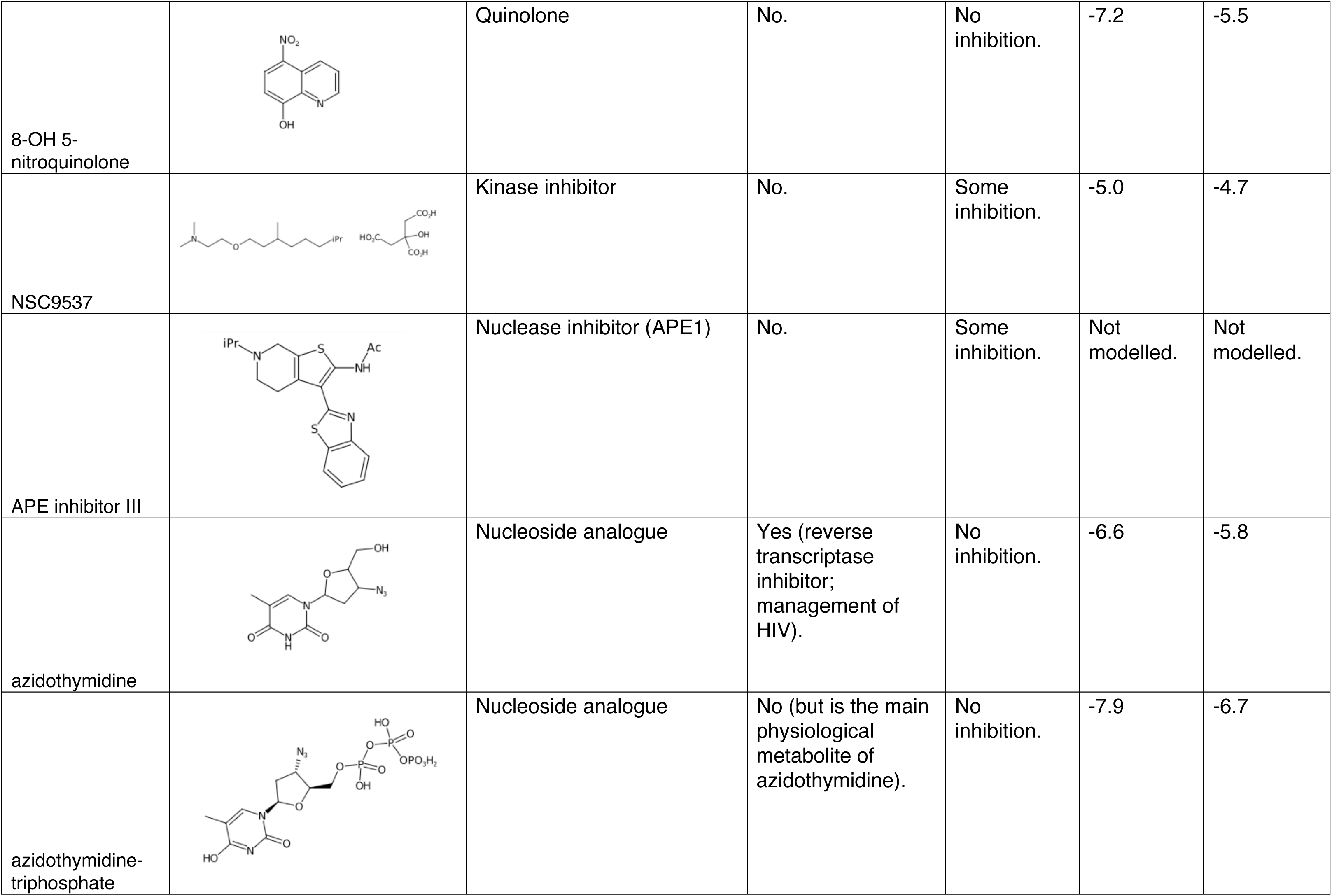
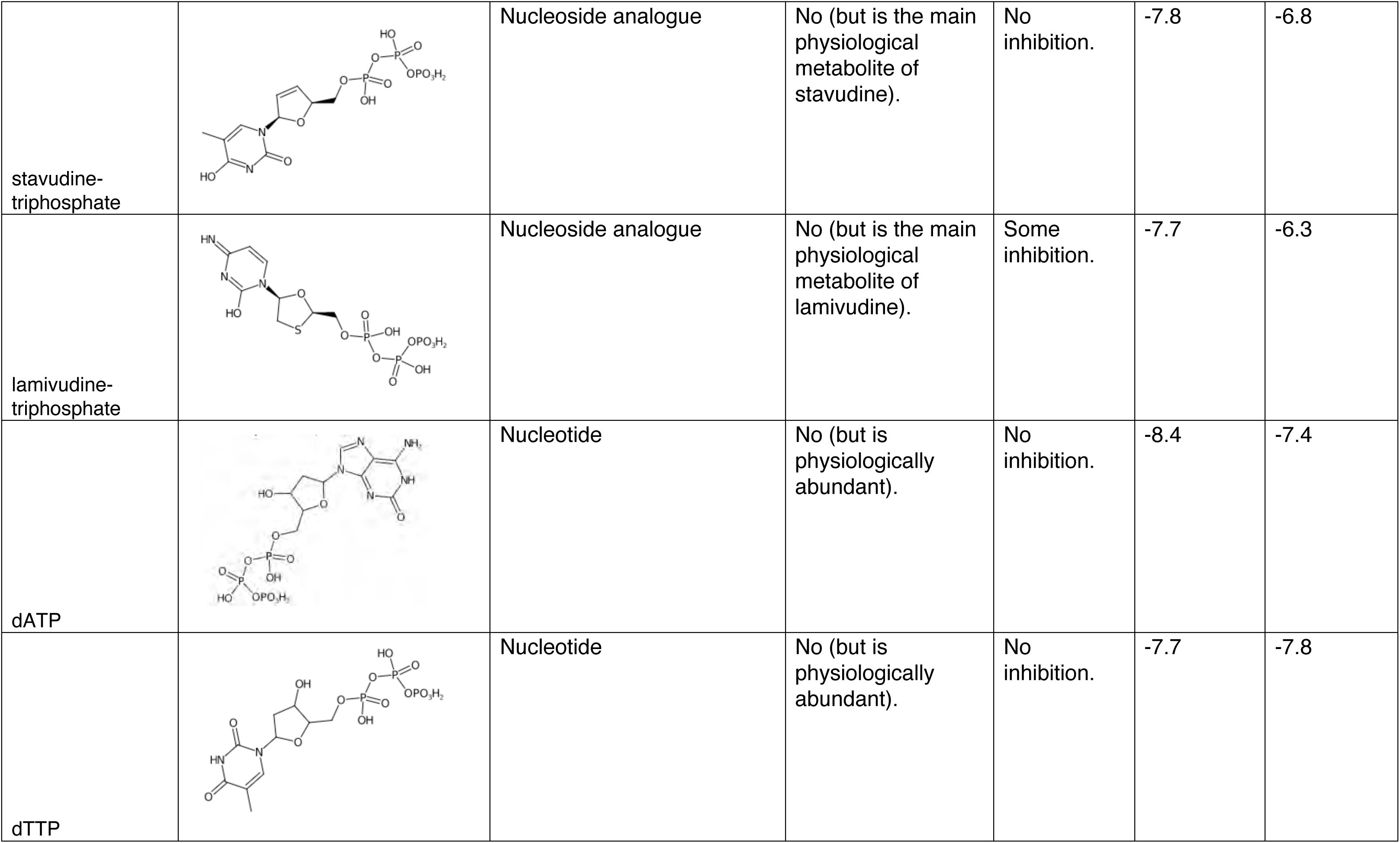
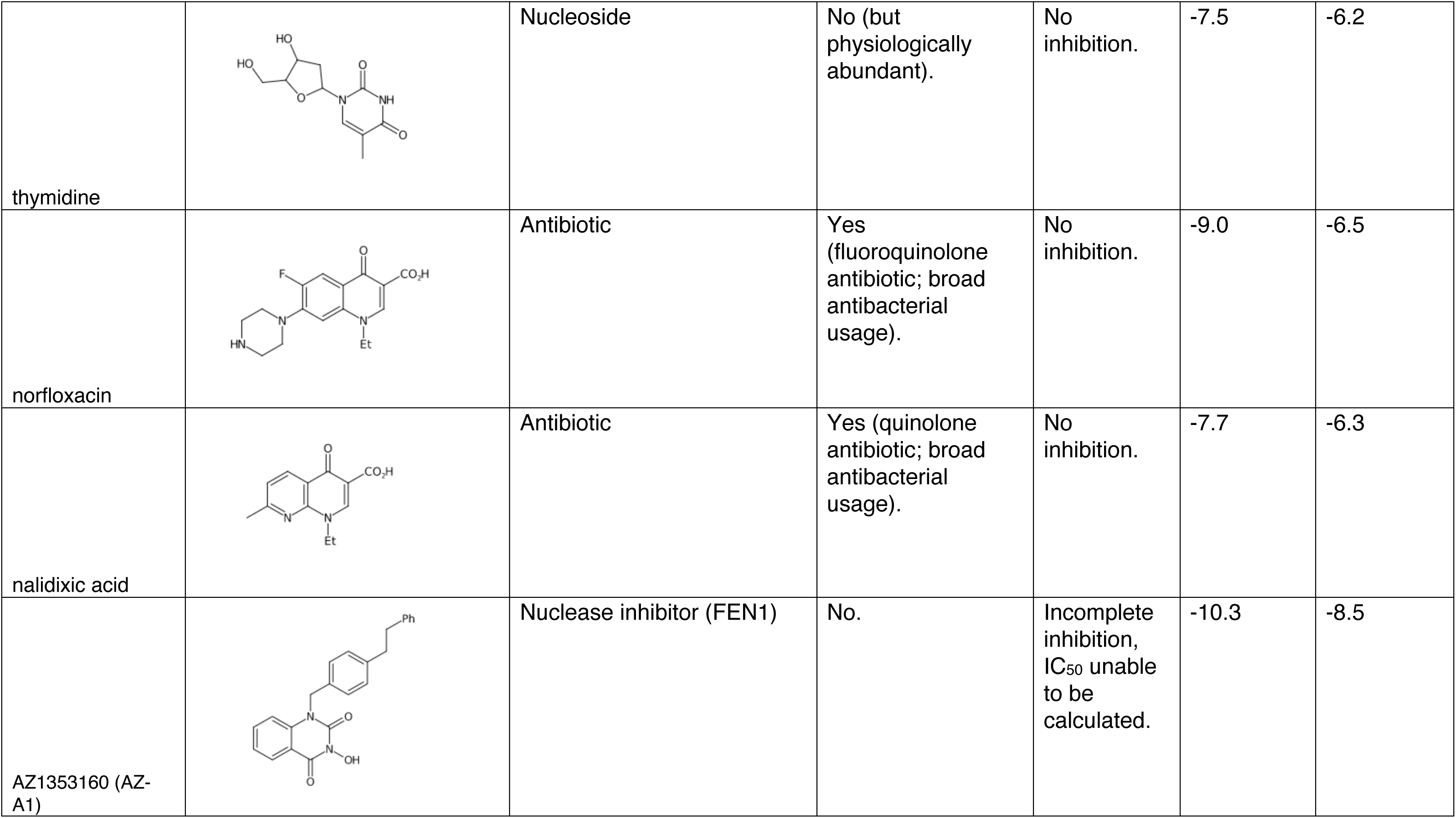
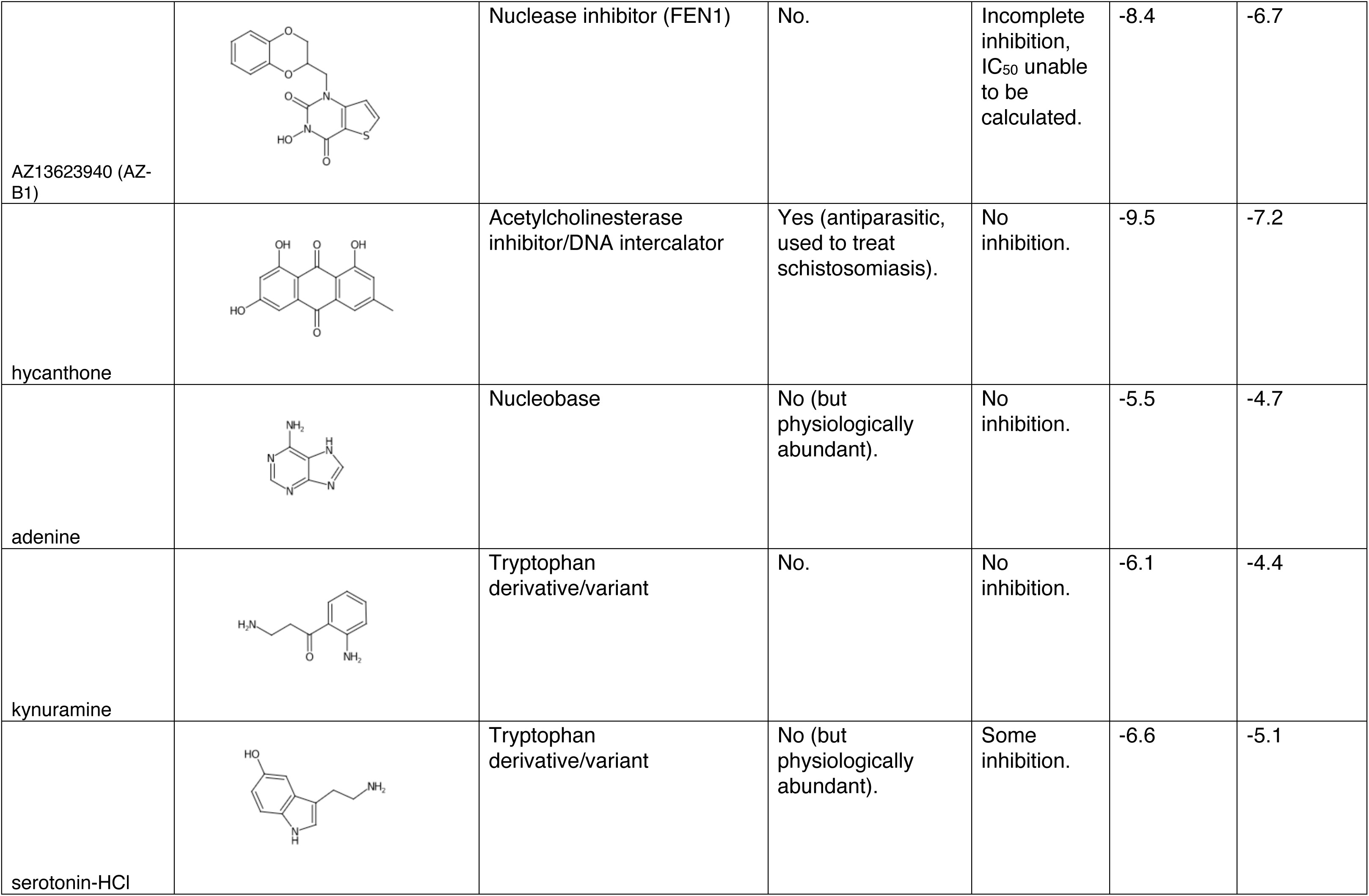
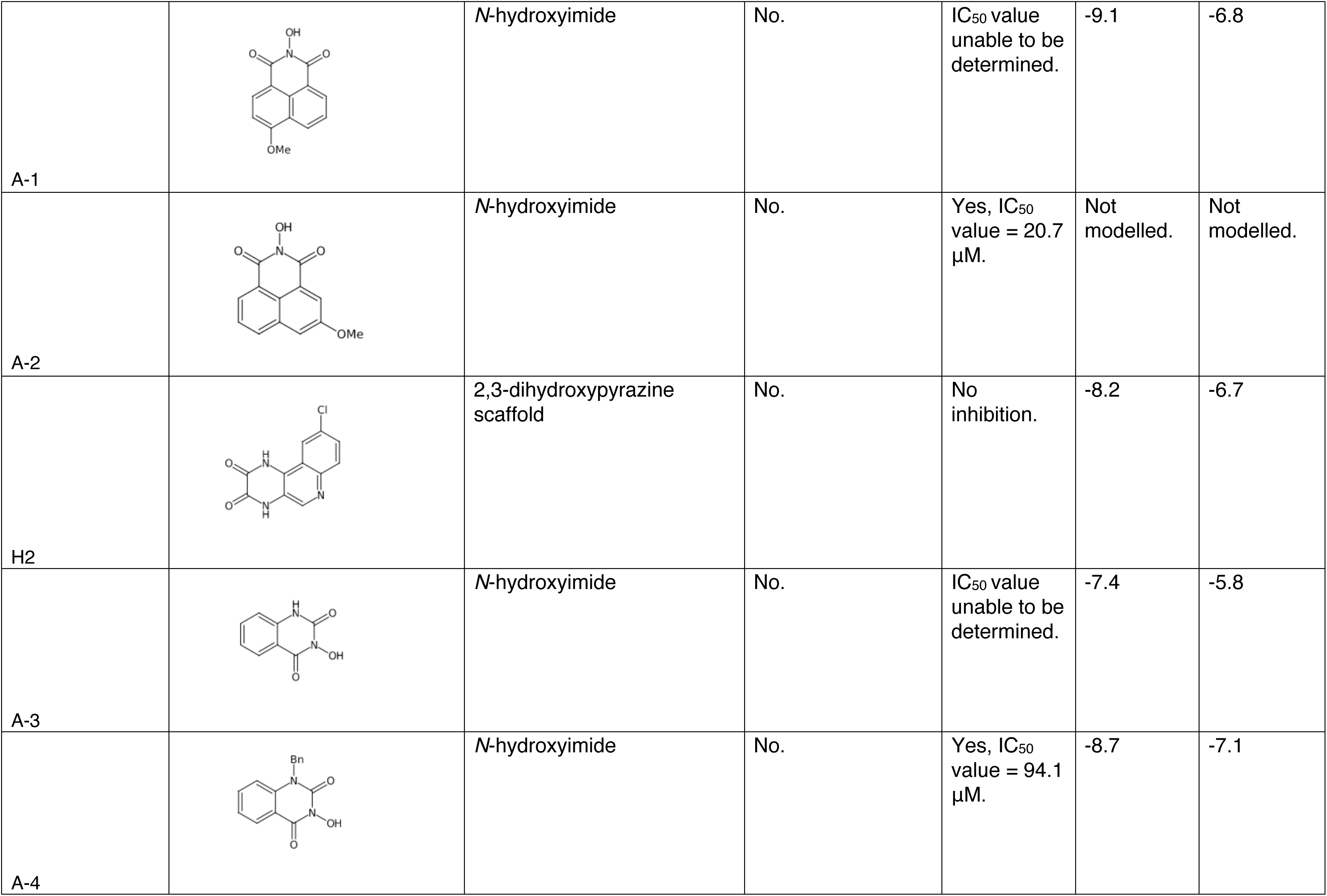
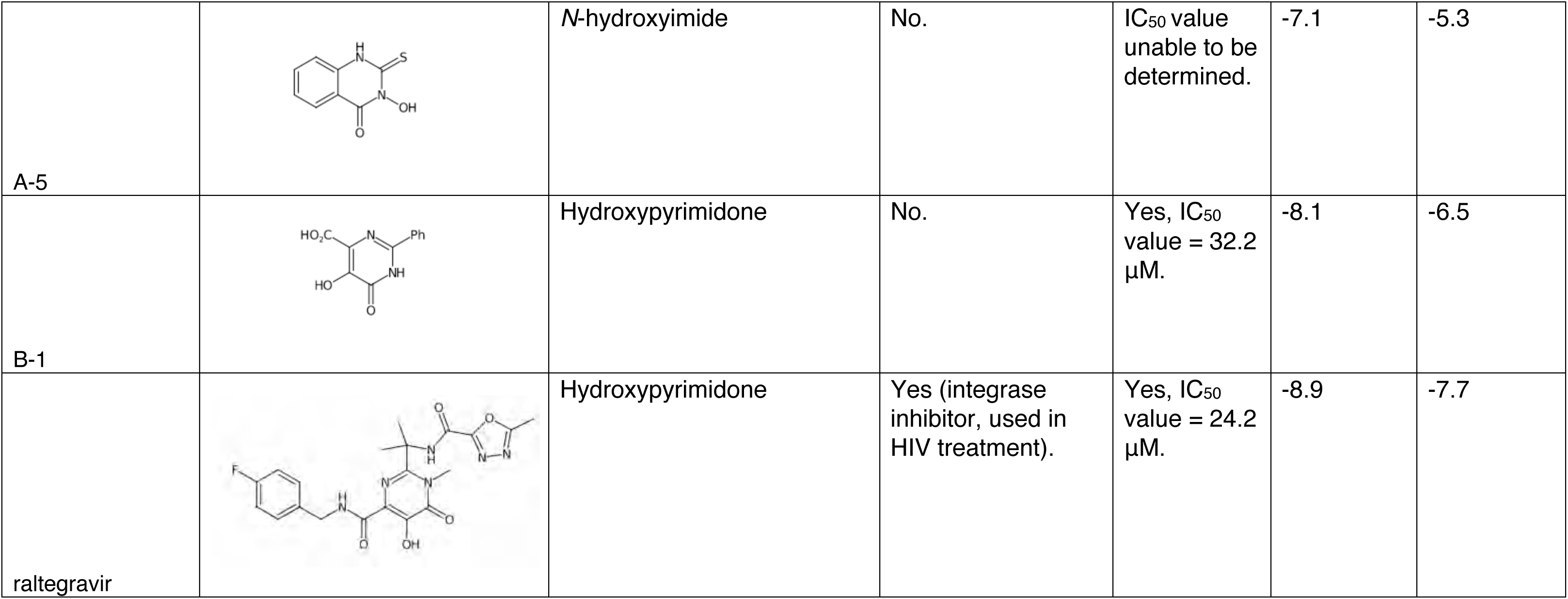
List of compounds tested for *in vitro* inhibition on nsp14-nsp10 nuclease activity. Compound structures, functional groupings, and FDA approval status are shown alongside *in vitro* inhibition activity and AutoDock Vina scores of the highest-affinity binding modes for each compound. See Fig. 4, Fig. 5, Fig. 6, Suppl. Fig. 4, Suppl. Fig. 5, Suppl. Fig. 7, Suppl. Fig. 8 for relevant data.

| Compound | Structure | Function | FDA approval | Level of inhibition | AutoDock Vina Full Surface (kcal/mol) | AutoDock Vina Active Site (kcal/mol) |
| --- | --- | --- | --- | --- | --- | --- |
| 7-nitroindole-2-carboxylic acid (CRT0044876) |  | Nuclease inhibitor (APE1) | No. | Some inhibition. | -7.5 | -6.2 |
| AraC (cytarabine) |  | Nucleoside analogue | Yes (treatment of acute myeloid leukaemia, acute lymphocytic leukaemia, lymphoma). | No inhibition. | -6.1 | -5.7 |
| aurintricarboxylic acid |  | RNase inhibitor | No. | Yes, IC<sub>50</sub> = 7.4 μM. | -10.6 | -7.5 |
| brivudine |  | Nucleoside analogue | Yes (antiviral; treatment of herpes zoster). | No inhibition. | -7.4 | -6.4 |
| ciprofloxacin |  | Antibiotic | Yes (quinolone antibiotic; broad antibacterial usage). | Incomplete inhibition, IC<sub>50</sub> unable to be calculated. | -9.2 | -6.6 |
| didanosine |  | Nucleoside analogue | Yes (reverse transcriptase inhibitor; management of HIV). | No inhibition. | -7.1 | -5.8 |
| doxorubicin |  | Topoisomerase inhibitor | Yes (chemotherapeutic; broadly used, including, breast cancer, bladder cancer, lymphoma, and acute lymphocytic leukaemia). | Some inhibition. | -10.1 | -8.0 |
| ebselen |  | Thiol-reactive | In phase III clinical trials (bipolar disorder, hearing loss, sensory disorders). | Yes, IC<sub>50</sub> = 3.3 μM. | -7.9 | -5.9 |
| emodin |  | Quinone | No, but available as a dietary supplement. | No inhibition. | -9.0 | -7.3 |
| famotidine |  | H2 receptor antagonist | Yes (antacid; treatment of peptic ulcer disease, gastroesophageal reflux disease). | No inhibition. | -6.8 | -5.6 |
| gemcitabine |  | Nucleoside analogue | Yes (broadly used chemotherapeutic, including for breast cancer, ovarian cancer, non-small cell lung cancer). | No inhibition. | -7.1 | -5.5 |
| lamivudine |  | Nucleoside analogue | Yes (reverse transcriptase inhibitor; management of HIV). | No inhibition. | -6.3 | -5.4 |
| melatonin |  | Tryptophan derivative (hormone) | No, but available as a dietary supplement, and a natural metabolite. | Some inhibition. | -7.4 | -5.9 |
| pimodivir |  | Polymerase inhibitor | In phase II clinical trials (treatment of influenza virus). | No inhibition. | -9.9 | -8.1 |
| remdesivir |  | Nucleoside analogue | Yes (suggested prodrug treatment for ebola and COVID-19). | No inhibition. | -9.5 | -7.0 |
| stavudine |  | Nucleoside analogue | Yes (reverse transcriptase inhibitor; management of HIV). | No inhibition. | -5.8 | -5.7 |
| deferoxamine |  | Chelating agent | Yes (treatment of iron overdose, haemachromatosis, aluminium toxicity). | No inhibition. | -6.9 | -6.3 |
| auranofin |  | Thiol-reactive | Yes (treatment of rheumatoid arthritis). | Incomplete inhibition, IC<sub>50</sub> unable to be calculated. | Not modelled. | Not modelled. |
| γ-ATP |  | Nucleotide analogue | No. | Some inhibition. | -7.4 | -7.2 |
| GS441524 |  | Nucleoside analogue | No (but the main physiological metabolite of remdesivir). | No inhibition. | -7.1 | -6.3 |
| CldU |  | Nucleoside analogue | No. | No inhibition. | -7.0 | -6.1 |
| IdU |  | Nucleoside analogue | No. | No inhibition. | -6.6 | -6.3 |
| mirin |  | Nuclease inhibitor (Mre11) | No. | No inhibition. | -7.2 | -5.7 |
| 7-ACA |  | Antibiotic | No (but core of the cephalosporin antibiotics). | No inhibition. | -6.2 | -5.9 |
| olaparib |  | PARP inhibitor | Yes (chemotherapeutic; used for treatment of homologous recombination defective breast and ovarian cancers). | No inhibition. | -11.4 | -9.1 |
| etoposide |  | Topoisomerase inhibitor | Yes<br>(chemotherapeutic;<br>broadly used,<br>including for<br>testicular cancer,<br>lung cancer,<br>lymphoma). | No<br>inhibition. | -8.4 | -7.8 |
| R9-AMK |  | Tryptophan<br>derivative/variant | No. | Some<br>inhibition. | -6.9 | -5.5 |
| valproic acid |  | GABAergic compound,<br>HDAC inhibitor | Yes (treatment of<br>epilepsy and<br>bipolar disorder). | Some<br>inhibition. | -5.0 | -4.3 |
| camptothecin |  | Topoisomerase inhibitor | No (but four<br>camptothecin<br>analogues are<br>approved for<br>cancer<br>chemotherapy). | Incomplete<br>inhibition,<br>IC<sub>50</sub> unable<br>to be<br>calculated. | -8.0 | -8.1 |
| thiram |  | Thiol-reactive | No. | Yes, IC<sub>50</sub> =<br>48.2 μM. | -3.7 | -3.4 |
| R7 AFMK |  | Tryptophan derivative/variant | No. | Some inhibition. | -6.3 | -5.5 |
| 6-OH-melatonin |  | Tryptophan derivative/variant | No. | Some inhibition | -6.2 | -5.8 |
| N-ac-serotonin |  | Tryptophan derivative/variant | No. | Some inhibition. | -7.1 | -6.0 |
| 5FOA |  | Derivative of pyrimidine precursor | No. | Some inhibition. | -6.2 | -5.4 |
| captopril |  | ACE inhibitor | Yes (treatment of hypertension and congestive heart failure). | No inhibition. | -5.2 | -4.5 |
| PBIT |  | Thiol-reactive | No (variant of ebselen). | Some inhibition. | -8.6 | -6.0 |
| disulfiram |  | Thiol-reactive | Yes (acetaldehyde inhibitor, management of alcoholism). | Yes, IC<sub>50</sub> value = 89 μM. | -4.0 | -3.9 |
| IPA3 |  | Kinase inhibitor | No. | Some inhibition. | -9.5 | -7.6 |
| 6-thio-guanine |  | Purine base analogue | Yes (chemotherapeutic; used for. treatment of acute myeloid leukaemia, acute lymphocytic leukaemia, chronic myeloid leukaemia). | No inhibition. | -5.7 | -5.0 |
| 8-OH 5-nitroquinolone |  | Quinolone | No. | No inhibition. | -7.2 | -5.5 |
| NSC9537 |  | Kinase inhibitor | No. | Some inhibition. | -5.0 | -4.7 |
| APE inhibitor III |  | Nuclease inhibitor (APE1) | No. | Some inhibition. | Not modelled. | Not modelled. |
| azidothymidine |  | Nucleoside analogue | Yes (reverse transcriptase inhibitor; management of HIV). | No inhibition. | -6.6 | -5.8 |
| azidothymidine-triphosphate |  | Nucleoside analogue | No (but is the main physiological metabolite of azidothymidine). | No inhibition. | -7.9 | -6.7 |
| stavudine-triphosphate |  | Nucleoside analogue | No (but is the main physiological metabolite of stavudine). | No inhibition. | -7.8 | -6.8 |
| lamivudine-triphosphate |  | Nucleoside analogue | No (but is the main physiological metabolite of lamivudine). | Some inhibition. | -7.7 | -6.3 |
| dATP |  | Nucleotide | No (but is physiologically abundant). | No inhibition. | -8.4 | -7.4 |
| dTTP |  | Nucleotide | No (but is physiologically abundant). | No inhibition. | -7.7 | -7.8 |
| thymidine |  | Nucleoside | No (but physiologically abundant). | No inhibition. | -7.5 | -6.2 |
| norfloxacin |  | Antibiotic | Yes (fluoroquinolone antibiotic; broad antibacterial usage). | No inhibition. | -9.0 | -6.5 |
| nalidixic acid |  | Antibiotic | Yes (quinolone antibiotic; broad antibacterial usage). | No inhibition. | -7.7 | -6.3 |
| AZ1353160 (AZ-A1) |  | Nuclease inhibitor (FEN1) | No. | Incomplete inhibition, IC<sub>50</sub> unable to be calculated. | -10.3 | -8.5 |
| AZ13623940 (AZ-B1) |  | Nuclease inhibitor (FEN1) | No. | Incomplete inhibition, IC<sub>50</sub> unable to be calculated. | -8.4 | -6.7 |
| hycanthone |  | Acetylcholinesterase inhibitor/DNA intercalator | Yes (antiparasitic, used to treat schistosomiasis). | No inhibition. | -9.5 | -7.2 |
| adenine |  | Nucleobase | No (but physiologically abundant). | No inhibition. | -5.5 | -4.7 |
| kynuramine |  | Tryptophan derivative/variant | No. | No inhibition. | -6.1 | -4.4 |
| serotonin-HCl |  | Tryptophan derivative/variant | No (but physiologically abundant). | Some inhibition. | -6.6 | -5.1 |
| A-1 |  | *N*-hydroxyimide | No. | IC<sub>50</sub> value<br>unable to be<br>determined. | -9.1 | -6.8 |
| A-2 |  | *N*-hydroxyimide | No. | Yes, IC<sub>50</sub><br>value = 20.7<br>μM. | Not<br>modelled. | Not<br>modelled. |
| H2 |  | 2,3-dihydroxypyrazine<br>scaffold | No. | No<br>inhibition. | -8.2 | -6.7 |
| A-3 |  | *N*-hydroxyimide | No. | IC<sub>50</sub> value<br>unable to be<br>determined. | -7.4 | -5.8 |
| A-4 |  | *N*-hydroxyimide | No. | Yes, IC<sub>50</sub><br>value = 94.1<br>μM. | -8.7 | -7.1 |
| A-5 |  | N-hydroxyimide | No. | IC<sub>50</sub> value unable to be determined. | -7.1 | -5.3 |
| B-1 |  | Hydroxypyrimidone | No. | Yes, IC<sub>50</sub> value = 32.2 μM. | -8.1 | -6.5 |
| raltegravir |  | Hydroxypyrimidone | Yes (integrase inhibitor, used in HIV treatment). | Yes, IC<sub>50</sub> value = 24.2 μM. | -8.9 | -7.7 |

## SUPPLEMENTAL MATERIALS AND METHODS

### Cloning and site directed mutagenesis of wildtype and nuclease dead (NSP14^D113A/E115A^) NSP14-10 complex

Wildtype (WT) NSP14 and NSP10 encoding DNA were codon-optimised for *Escherichia coli (E. coli)* expression and were synthesised by Twist Bioscience (California, USA). Both the NSP14 and NSP10 genes were subcloned into the pNIC28-BSA4 vector^1^ to enable bicistronic expression and appropriately stoichiometric production of NSP14 and NSP10 proteins.

Site-directed-mutagenesis (SDM) was carried out using an ‘inverse’ PCR experiment, whereby an entire plasmid is amplified using complementary mutagenic primers with minimal cloning steps^2^. The Herculase II Fusion DNA Polymerase (Agilent) was utilised and SDM PCR was performed to amplify the whole plasmid, according to the manufacturers instructions. The PCR product was then added to a standard KLD enzyme mix (NEB) reaction and was incubated at room temperature for 1 hour, prior to transformation into chemically competent *E. coli* cells.

### Compound synthesis

All reagents were from Sigma-Aldrich, Acros Organics, Fluka, Fluorochem, Abcr, or Fisher Scientific and used without further purification. Flash column chromatography was performed using a Biotage Isolera automated flash column chromatography platform using Biotage KP-SNAP-Sil or SNAP Ultra columns. IR spectra were recorded using a Bruker Tensor 27 FT-IR spectrometer in the solid state or as a thin film. Selected characteristic peaks are reported in cm^−1^. NMR spectra were recorded using Bruker Avance spectrometers in the deuterated solvent stated. Assignments given correspond with the numbering system as drawn; where assignments are not given it was not possible to assign the resonances due to overlapping signals. Low-resolution mass spectra were recorded using an Agilent Technologies 1260 Infinity LC-MS system fitted with a 6120 Quadrupole mass spectrometer.

High-resolution mass spectra (HRMS) were recorded in HPLC grade methanol using electrospray ionisation (ESI+) on a Bruker APEX III FT-ICR mass spectrometer.

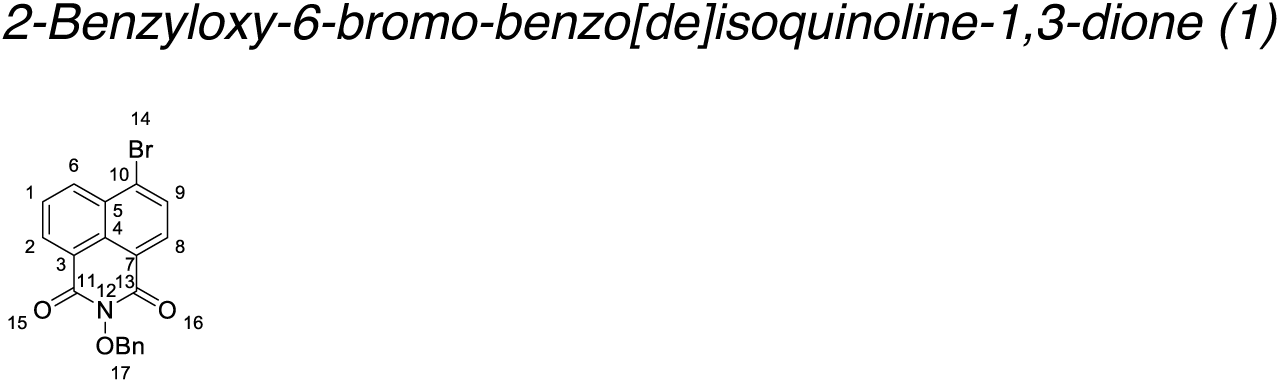

To stirred solution of 6-bromo-benzo[de]isochromene-1,3-dione (300 mg, 1.08 mmol) in anhydrous pyridine (10 ml) was added O-benzylhydroxylamine hydrochloride (346 mg, 2.16 mmol). The reaction was stirred under reflux for 2 h under an inert atmosphere. After the reaction was complete, it was cooled to room temperature and concentrated *in vacuo*. To the resultant solid residue 20 mL of EtOH was added; the precipitate was filtered, washed with cold EtOH and dried to give 2-benzyloxy-6-bromo-benzo[de]isoquinoline-1,3-dione (395 mg, 96%) as an orange solid. ^1^H NMR (400 MHz, CDCl_3_) δ_H_ 8.63 (dd, *J* = 7.3, 1.2 Hz, 1H, C(6)H), 8.55 (dd, *J* = 8.5, 1.2 Hz, 1H, C(2)H), 8.38 (d, *J* = 7.9 Hz, 1H, C(9)H, 8.00 (d, *J* = 7.9 Hz, 1H, C(8)H), 7.81 (dd, *J* = 8.5, 7.3 Hz, 1H. C(1)H), 7.66 – 7.57 (m, 2H, OBn(17) ArH_2_), 7.39 – 7.28 (m, 3H, OBn(17) ArH_3_), 5.20 (s, 2H, OBn(17) -CH_2_-); ^13^C NMR (101 MHz, CDCl_3_) δ_C_ 160.53, 133.99, 133.94, 132.51, 131.60, 131.35, 131.12, 130.90, 130.02, 129.21, 128.54, 128.32, 128.30, 123.31, 122.42, 78.70; LRMS: *m/z*(%)= 382.2 (100), 384.2 (98) [M + H]^+^.

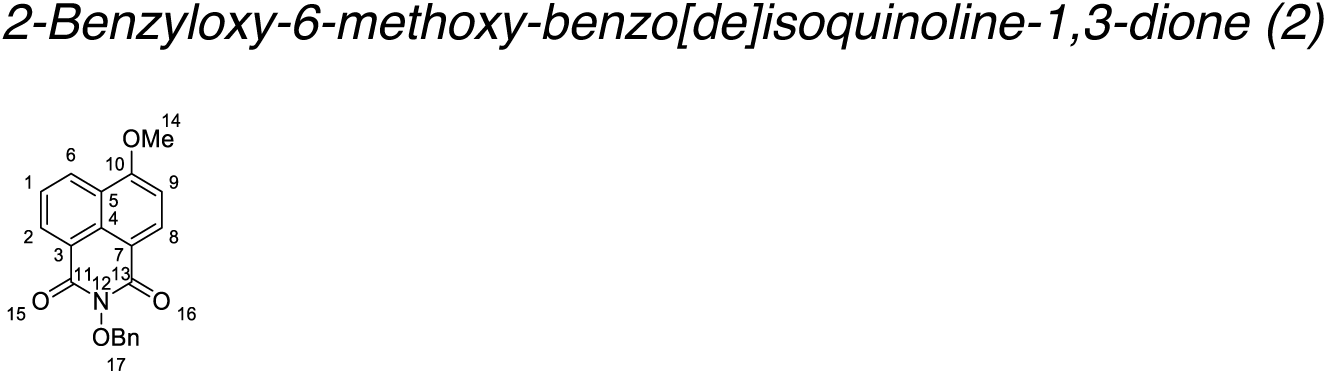

To stirred solution of **1** (360 mg, 0.94 mmol) and CuSO_4_ (10 mg) in anhydrous MeOH (8 mL), NaOMe solution 25 (wt/wt %) in MeOH (0.27 mL, 1.43mmol) were added. The reaction was stirred under reflux for 12 h under an inert atmosphere. After the reaction was complete it was cooled to room temperature; the resultant precipitate was filtered, washed with H_2_O and dried to give 2-benzyloxy-6-methoxy-benzo[de]isoquinoline-1,3-dione (286 mg, 90%) as a yellow solid. ^1^H NMR (400 MHz, CDCl_3_) δ_H_ 8.64 (dd, *J* = 7.3, 1.3 Hz, 1H, C(6)H), δ 8.60 (dd, *J* = 8.4, 1.2 Hz, 1H, C(2)H), 8.59 (d, *J* = 8.3 Hz, 1H, C(8)H), 7.72 (dd, *J* = 8.4, 7.3 Hz, 1H, C(1)H), 7.69 (dd, *J* = 7.8, 1.7 Hz, 2H, OBn(17) ArH_2_), 7.46 – 7.33 (m, 3H, OBn(17) ArH_3_), 7.07 (d, *J* = 8.3 Hz, 1H, C(9)H), 5.26 (s, 2H, OBn(17) -CH_2_-).), 4.14 (s, 3H, OMe(14) -CH_3_). ^13^C NMR (101 MHz, CDCl_3_) δ_C_ 161.49, 161.47, 161.00, 134.40, 134.09, 132.09, 130.12, 129.42, 129.16, 128.82, 128.61, 126.24, 123.91, 122.70, 115.19, 105.58, 78.64, 56.48. LRMS: *m/z*(%)= 334.2 (100) [M + H]^+^.

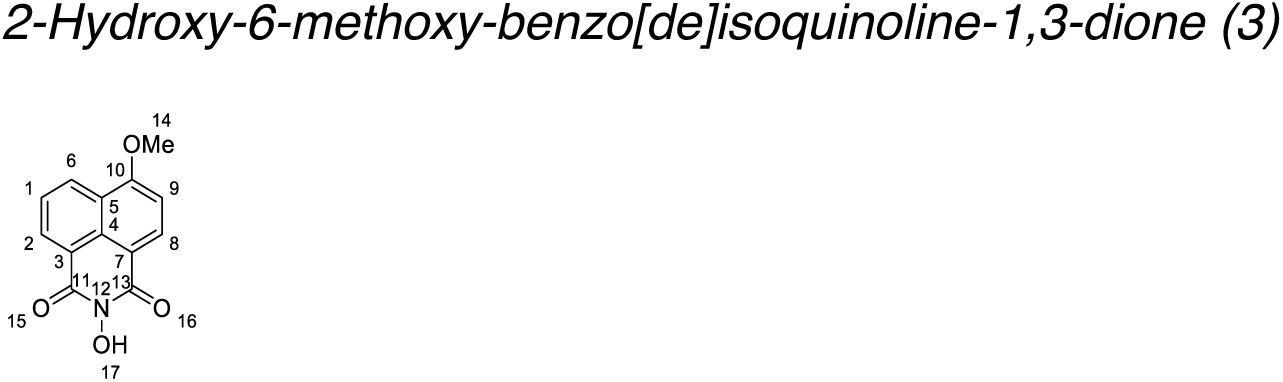

To a stirred solution of **2** (275 mg, 0.83 mmol) in dimethylacetamide (10 mL), palladium on carbon (35 mg, 10 wt. %) was added. The reaction was then stirred for 4 h at room temperature under a hydrogen atmosphere. The reaction was filtered through Celite^®^, washed with dimethylacetamide, then concentrated *in vacuo*. The resultant solid residue was washed with MeO, then dried to give 2-hydroxy-6-methoxy-benzo[de]isoquinoline-1,3-dione (194 mg, 96%). mp 259-261°C. ^1^H NMR (400 MHz, DMSO-*d*_6_) δ_H_ 10.63 (brs, 1H, O(17)H), 8.51 (app. ddd, *J* = 8.5, 7.8, 1.2 Hz, 2H, C(6)H, C(2)H), 8.46 (d, *J* = 8.3 Hz, 1H, C(8)H), 7.81 (dd, *J* = 8.4, 7.3 Hz, 1H, C(1)H), 7.32 (d, *J* = 8.4 Hz, 1H, C(9)H), 4.13 (s, 3H, OMe(14) -CH_3_). ^13^C NMR (101 MHz, DMSO-*d*_6_) δ_C_ 160.91, 160.55, 160.48, 133.42, 131.13, 128.47, 127.44, 126.51, 122.90, 122.20, 114.40, 106.43, 56.70; IR: 3659, 3183, 2981, 1703, 1650, 1589, 1576, 1459, 1382, 1366, 1268, 1233, 1187, 1158, 1141, 1081, 1049, 1026, 947, 810, 773, 746, 650; . LRMS: *m/z*(%)= 244.1 (100) [M + H]^+^; HRMS: calcd. C_13_H_9_O_4_N^23^Na [M + Na]^+^: 266.04238 ; found: 266.04261.

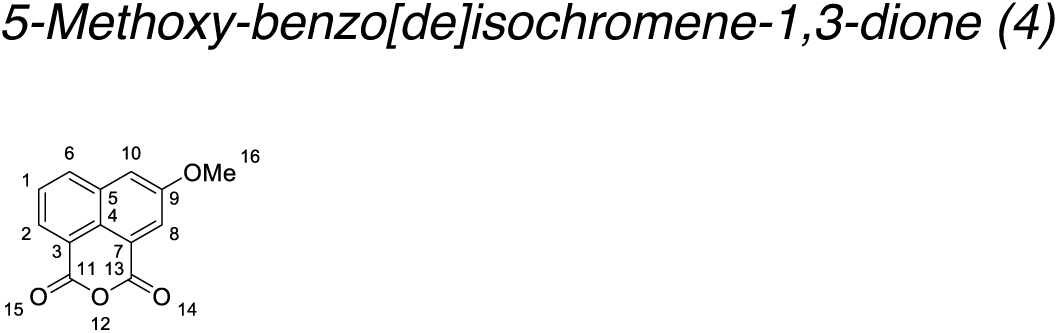

To a solution 5-hydroxy-benzo[de]isochromene-1,3-dione (100 mg. 0.46 mmol) in anhydrous acetone (5mL), SO_4_Me_2_ (0.8 ml, 0.93 mmol) and K_2_CO_3_ (200 mg, 1.45 mmol) were added. The reaction mixture was stirred under reflux for 3 h under an inert atmosphere and concentrated *in vacuo*. 1M HCl (20 mL) was added. The resultant precipitate was filtered, washed with H_2_O and dried to give 5-methoxy-benzo[de]isochromene-1,3-dione (78 mg, 74%) as a cream solid^3^. ^1^H NMR (400 MHz, DMSO-*d*_6_) δ_H_ 8.42 (dd, *J* = 8.3, 1.1 Hz, 1H, C(2)H), 8.35 (dd, *J* = 7.3, 1.1 Hz, 1H, C(6)H), 8.06 (d, *J* = 2.5 Hz, 1H, C(8)H), 8.02 (d, *J* = 2.6 Hz, 1H, C(10)H), 7.86 (dd, *J* = 8.3, 7.3 Hz, 1H, C(1)H), 4.01 (s, 3H, OMe(16) -CH_3_). ^13^C NMR (101 MHz, DMSO-*d*_6_) δ_C_ 161.1, 160.8, 158.3, 134.5, 133.7, 130.2, 128.5, 125.6, 123.7, 121.0, 119.4, 114.8, 56.6. LRMS: *m/z*(%)= 229.1 (100) [M + H]^+^.

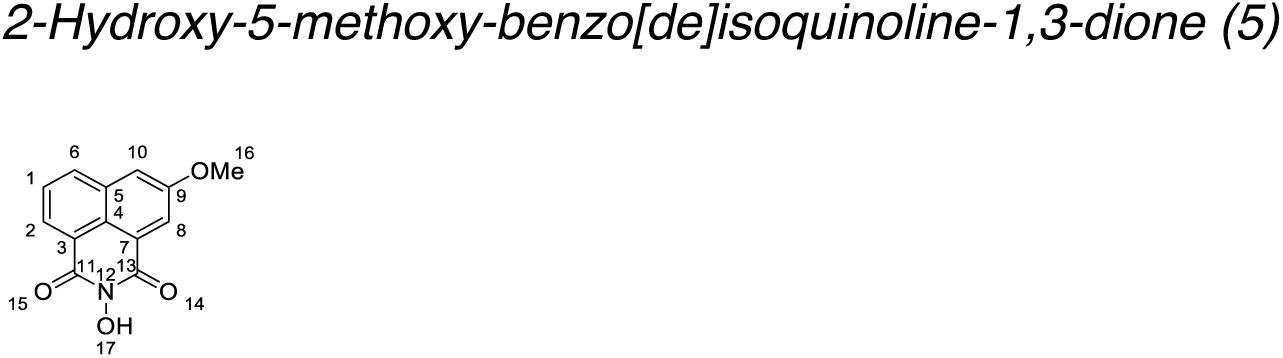

To solution of **4** (78 mg, 0.33 mmol) in anhydrous pyridine (3 mL), hydroxylamine hydrochloride (45 mg, 0.66 mmol) was added. The reaction mixture was stirred under reflux for 1 h under inert atmosphere. The reaction mixture was then diluted with cold H_2_O (75 mL); the precipitate was then filtered, washed with H_2_O and dried to give 2-hydroxy-5-methoxy-benzo[de]isoquinoline-1,3-dione (81 mg, 97%) as a yellow solid^3^. ν_max_: 3658, 3409, 2981, 2161, 2032, 1711, 1628, 1597, 1581, 1472, 1431, 1382, 1337, 1274, 1241, 1160, 1060, 1034, 1016, 959, 872, 831, 776, 739, 723; ^1^H NMR (400 MHz, DMSO-*d*_6_) δ_H_ 10.68 (brs, 1H, O(17)H) 8.31 (d, *J* = 7.5 Hz, 2H, C(2)H), 8.01 (d, *J* = 2.4 Hz, 1H, C(8)), 7.89 (d, *J* = 2.6 Hz, 1H, C(10)H), 7.80 (app. t, *J* = 7.7 Hz, 1H, C(1)H), 3.97 (s, 3H, OMe(16) -CH_3_); ^13^C NMR (101 MHz, DMSO-*d*_6_) δ_C_ 161.36, 160.91, 158.20, 133.75, 133.67, 128.66, 128.17, 124.29, 122.61, 122.20, 122.11, 113.86, 56.45; LRMS: *m/z*(%)= 244.1 (100) [M + H]^+^.

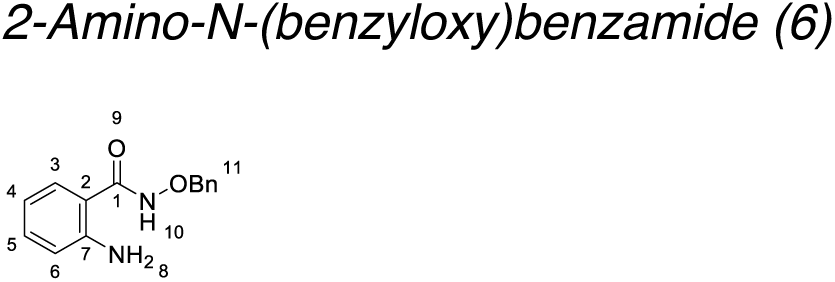

To solution of isatoic anhydride (6 g, 36.8 mmol) and O-benzylhydroxylamine hydrochloride (8.6 g, 54.0 mmol) in anhydrous THF (50 mL), was added triethylamine (7.5 mL, 54.0 mmol). The reaction mixture was stirred under reflux for 3 h under an inert atmosphere, then concentrated *in vacuo*. The crude product was purified using flash column chromatography (10-30% EtOAc in cyclohexane) to give 2-amino-N-(benzyloxy)benzamide (7.8g, 67%) as a white solid^4^. ^1^H NMR (400 MHz, DMSO-*d*_6_) δ_H_ 11.65 (s, 1H, N(10)H), 7.96 (dd, *J* = 7.9, 1.5 Hz, 1H, C(3)H), 7.68 (ddd, *J* = 8.5, 7.2, 1.6 Hz, 1H, C(5)H), 7.60-7.53 (m, 2H, OBn(11) ArH_2_), 7.48-7.34 (m, 3H,, OBn(11) ArH_3_), 7.30-7.19 (m, 2H, C(6)H, C(4)H) 5.10 (s, 2H, OBn(11) -CH_2_-); ^13^C NMR (101 MHz, DMSO-*d*_6_) δ_C_ 159.50, 148.53, 139.18, 135.56, 134.95, 129.93, 129.32, 128.84, 127.72, 123.17, 115.93, 114.91. 77.95; LRMS: *m/z*(%)= 243.1 (100) [M + H]^+^.

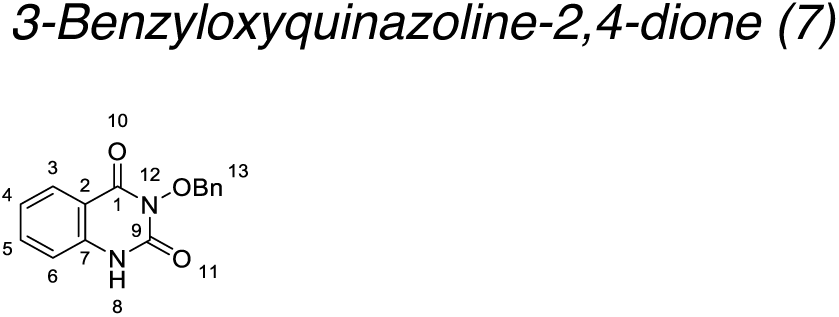

To solution of **6** (1 g, 4,1 mmol) and triphosgene (1.35 g, 4.5 mmol) in anhydrous THF (40 mL) at 0 °C, TEA (1.26 mL, 9.0 mmol) was added dropwise. The reaction mixture was stirred for 2 h at room temperature under an atmosphere, then concentrated *in vacuo*. The crude product was purified using flash column chromatography (10-60% EtOAc in cyclohexane). 3-Benzyloxyquinazoline-2,4-dione (726 mg, 66%) was obtained as a white solid^4^. ^1^H NMR (400 MHz, DMSO-*d*_6_) δ_H_ 11.44 (s, 1H, N(8)H), 7.59 – 7.35 (m, 5H, OBn(13) ArH_5_), 7.29 (dd, *J* = 8.0, 1.6 Hz, 1H, C(3)H), 7.15 (ddd, *J* = 8.6, 7.0, 1.6 Hz, 1H, C(5)H), 6.71 (d, *J* = 8.2 Hz, 1H, C(6)H), 6.49 (t, *J* = 7.5 Hz, 1H C(4)H), 4.90 (s, 2H, OBn(13) -CH_2_-); ^13^C NMR (101 MHz, DMSO-*d*_6_) δ_C_ 167.68, 150.01, 136.52, 132.57, 129.33, 128.76, 128.71, 128.21, 116.79, 115.11, 112.81, 77.41; LRMS: *m/z*(%)= 269.1 (100) [M + H]^+^.

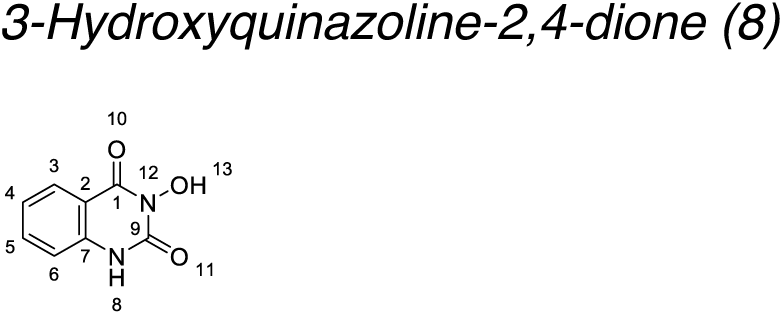

To a degassed solution of **7** (50 mg, 0.19 mmol) in DMF (2 mL), palladium on carbon (10 mg, 10 (wt/wt) %) was added. The reaction mixture was stirred for 2 h under H_2_ atmosphere, then filtered through Celite^®^, then concentrated *in vacuo* to give 3-hydroxyquinazoline-2,4-dione (28 mg, 87%) as a white solid^4^. ^1^H NMR (400 MHz, DMSO-*d*_6_) δ_H_ 11.05 (brs, 2H, N(8)H, O(13)H), 7.94 (d, *J* = 8.1 Hz, 1H, C(3) H), 7.66 (t, *J* = 7.7 Hz, 1H, C(5)H), 7.35-7.13 (m, 2H, C(4)H, C(5)H). ^13^C NMR (101 MHz, DMSO-*d*_6_) δ_C_ 159.87, 149.23, 138.84, 135.14, 127.53, 122.98, 115.75, 114.61.; LRMS: *m/z*(%)= 179.1 (100) [M + H]^+^;

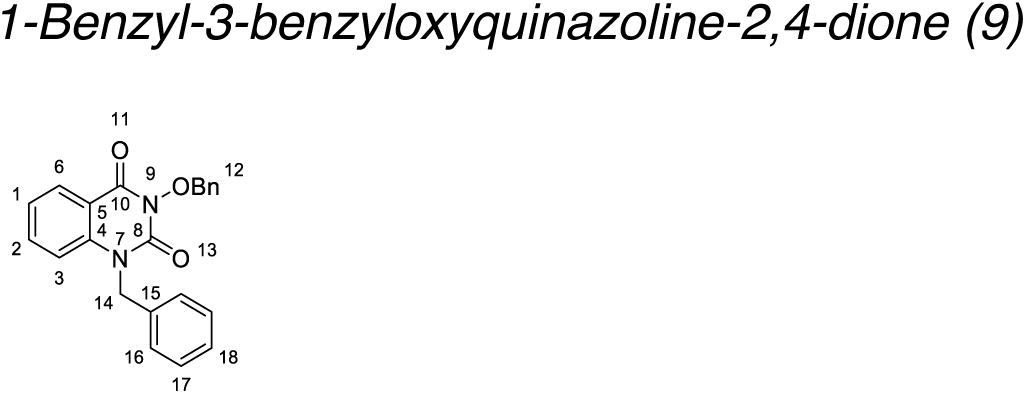

To solution of **7** (200 mg, 0.75 mmol) and Cs_2_CO_3_ (485 mg, 1.5 mmol) in anhydrous DMF (3 mL), benzyl bromide (106 µL, 089 mmol) was added. The reaction mixture was stirred at 80 °C for 2 h, then quenched with cold H_2_O (75mL); the resultant precipitate was collected by filtration, washed with water, and dried. 1-Benzyl-3-benzyloxyquinazoline-2,4-dione (256 mg, 95%) was obtained as a white solid (DOI: 10.1039/J39660000523, 523-527). ^1^H NMR (400 MHz, CDCl_3_) δ_H_ 8.26 (dd, *J* = 7.9, 1.7 Hz, 1H, C(6)H), 7.64 (dd, *J* = 6.5, 2.9 Hz, 2H, C(2)H), 7.55 (ddd, *J* = 8.8, 7.3, 1.7 Hz, 1H, C(1)H), 7.45-7.16 (m, 10H, OBn(12) ArH_5,_ (C(16)H)_2_, (C(17)H)_2_, (C(18)H)), 7.11 (d, *J* = 8.5 Hz, 1H, C(3)H), 5.37 (s, 2H, (C(14)H_2_), 5.29 (s, 2H, OBn(12) -CH_2_-). ^13^C NMR (101 MHz, CDCl_3_) δ_C_ 158.70, 149.68, 139.20, 135.24, 135.22, 130.24, 129.18, 129.03, 128.99, 128.49, 127.82, 126.50, 123.42, 114.72, 78.43, 47.47; LRMS: *m/z*(%)= 359.2 (100) [M + H]^+^.

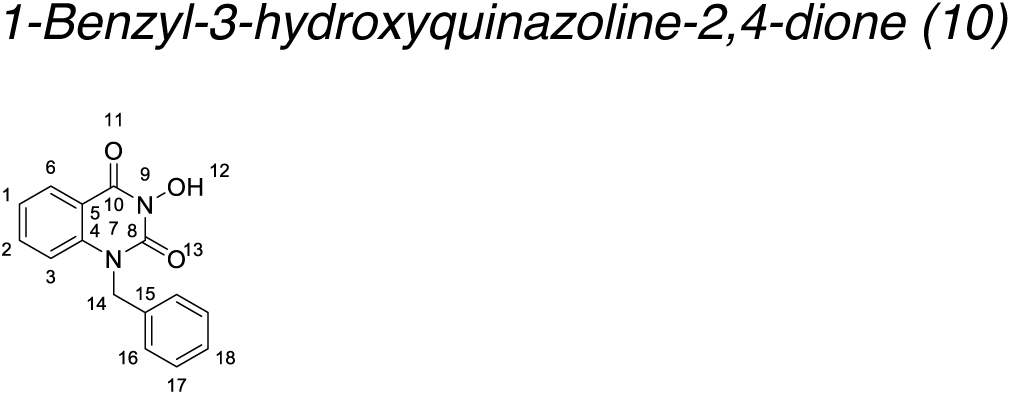

A solution of **8** (240 mg, 0.67 mmol) in conc. HBr (48% in H_2_O, 2mL) and AcOH (2mL) was stirred under reflux for 1 h. The reaction mixture was then concentrated *in vacuo*, dissolved in DMF (0.5 mL), poured onto to cold H_2_O (50 mL); precipitate was then filtered off and washed with minimal amount of cold CH_2_Cl_2_ to give 1-benzyl-3-hydroxyquinazoline-2,4-dione (51 mg, 28%)^5^. ^1^H NMR (400 MHz, DMSO-*d*_6_) δ_H_ 10.86 (s, 1H, O(12)H), 8.09 (dd, *J* = 7.8, 1.6 Hz, 1H, C6)H), 7.67 (ddd, *J* = 8.7, 7.2, 1.7 Hz, 1H, C(2)H), 7.39 – 7.22 (m, 7H, C(3)H, C(1)H, (C(16)H)_2_, (C(17)H)_2_, (C(18)H), 5.40 (s, 2H C(14)H_2_); ^13^C NMR (101 MHz, DMSO-*d*_6_) δ_C_ 158.96, 150.27, 139.13, 136.60, 135.44, 129.19, 128.23, 127.79, 126.91, 123.55, 115.70, 115.61, 46.87; LRMS: *m/z*(%)= 269.2 (100) [M + H]^+^.

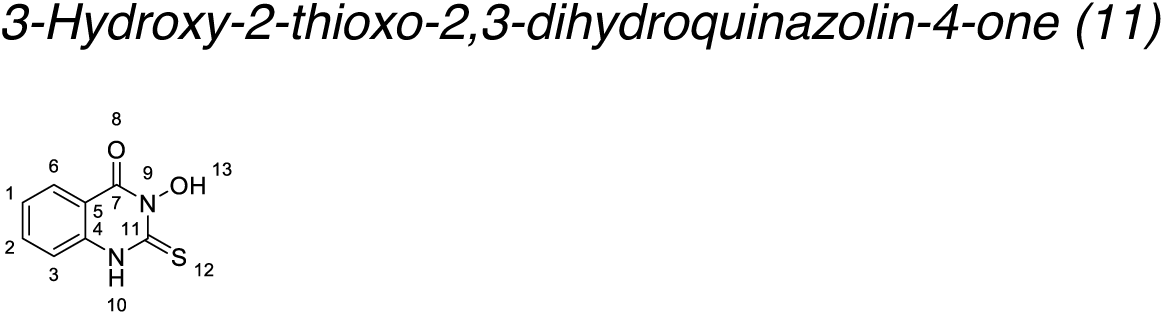

To solution of methyl anthranilate (150 µL, 1.16 mmol) in H_2_O (3 mL) and CH_2_Cl_2_ (10mL), thiophosgene (100 µL, 1.3 mmol) were added dropwise. The reaction mixture was stirred at room temperature overnight, then diluted with H_2_O (15 mL), extracted with CH_2_Cl_2_ (3 × 15 mL); the combined organic extracts were washed with brine (20 mL), dried over Na_2_SO_4_ then concentrated *in vacuo*. Crude methyl 2-isothiocyanatobenzoate was then added to solution of hydroxylamine hydroxyl (82 mg, 1.18 mmol) and sodium hydroxide (47 mg, 1.20 mmol) in mixture of CHCl_3_/H_2_O (1:1, 10 mL). The reaction mixture was stirred for 2h at room temperature, precipitate was filtered off, washed with CH_2_Cl_2_ then dried. 3-Hydroxy-2-thioxo-2,3-dihydroquinazolin-4-one (179 mg, 80%) was obtained as a white solid^6^. ^1^H NMR (400 MHz, DMSO-*d*_6_) δ_H_ 12.97 (s, 1H,, N(10)H), 11.24 (s, 1H, O(13)H), 7.98 (d, *J* = 8.0 Hz, 1H, C(6)H), 7.75 (t, *J* = 7.9 Hz, 1H C(2)H), 7.40 (d, *J* = 8.4 Hz, 1H, C(3)H), 7.35 (t, *J* = 7.6 Hz, 1H, C(1)H). ^13^C NMR (101 MHz, DMSO-*d*_6_) δ_C_ 173.26, 157.07, 139.07, 135.67, 127.47, 124.78, 116.32, 116.19; LRMS: *m/z*(%)= 195.1 (100) [M + H]^+^.

### Dose-response inhibition and data analysis

Approximate IC_50_ values were calculated (where possible) by analysing the inhibition of digestion of a radiolabelled RNA substrate. Gel images collected on a Typhoon scanner were analysed using Image J (NIH)^7^, determining the proportion of substrate remaining undigested (enzyme control) in comparison with the amount of that entered the gel; results are given as a percentage of digested substrate and were plotted against the log_10_ of the inhibitor concentration, dose response curves were fitted using nonlinear regression, and where possible IC_50_ values calculated. The data were plotted and IC_50_ values obtained in Graphpad Prism v8.3.0. Error is standard error.

Three or more gels were analysed for each inhibitor. Some data was excluded due to user determined factors such as: the quality of the gel being insufficient to obtain satisfactory data (e.g. gel cracked during drying), the signal of the radiation was below a minimum threshold, or the controls run on the gel were of unsatisfactory quality (e.g. bands running in a non-horizontal, or “smiling” format).

### Differential scanning spectrometry (DSF)

DSF experiments were carried out using the method of Niesen *et al*^8^. Each of the compounds were serially diluted in DSF buffer (25 mM HEPES pH 7.5, 50 mM NaCl, 5 mM MgCl_2_, 5% (v/v) glycerol, and 1 mM DTT). Note that DTT was excluded for experiments with thiram, disulfiram, and ebselen. The nsp14-10 complex (1 µM) was incubated in the presence of the compounds for 10 minutes at room temperature prior to the addition of SYPRO^TM^ Orange protein stain (1:10000 dilution). The fluorescence emission was measured using a fluorescence resonance energy transfer filter (560– 580 nm) with an excitation wavelength of 450–490 nm. During the DSF experiment, the temperature was increased from 25 to 95°C at an increment of 1°C per second.

## REFERENCES

1. Fung, T. S. & Liu, D. X. Human Coronavirus: Host-Pathogen Interaction. Annu Rev Microbiol 73, 529–557, doi:10.1146/annurev-micro-020518-115759 (2019).

2. Sevajol, M., Subissi, L., Decroly, E., Canard, B. & Imbert, I. Insights into RNA synthesis, capping, and proofreading mechanisms of SARS-coronavirus. Virus Res 194, 90–99, doi:10.1016/j.virusres.2014.10.008 (2014).

3. Bouvet, M. et al. RNA 3’-end mismatch excision by the severe acute respiratory syndrome coronavirus nonstructural protein nsp10/nsp14 exoribonuclease complex. Proc Natl Acad Sci U S A 109, 9372–9377, doi:10.1073/pnas.1201130109 (2012).

4. Eckerle, L. D., Lu, X., Sperry, S. M., Choi, L. & Denison, M. R. High fidelity of murine hepatitis virus replication is decreased in nsp14 exoribonuclease mutants. J Virol 81, 12135–12144, doi:10.1128/JVI.01296-07 (2007).

5. Smith, E. C., Blanc, H., Surdel, M. C., Vignuzzi, M. & Denison, M. R. Coronaviruses lacking exoribonuclease activity are susceptible to lethal mutagenesis: evidence for proofreading and potential therapeutics. PLoS Pathog 9, e1003565, doi:10.1371/journal.ppat.1003565 (2013).

6. Denison, M. R., Graham, R. L., Donaldson, E. F., Eckerle, L. D. & Baric, R. S. Coronaviruses: an RNA proofreading machine regulates replication fidelity and diversity. RNA Biol 8, 270–279, doi:10.4161/rna.8.2.15013 (2011).

7. Subissi, L. et al. SARS-CoV ORF1b-encoded nonstructural proteins 12-16: replicative enzymes as antiviral targets. Antiviral Res 101, 122–130, doi:10.1016/j.antiviral.2013.11.006 (2014).

8. Ma, Y. et al. Structural basis and functional analysis of the SARS coronavirus nsp14-nsp10 complex. Proc Natl Acad Sci U S A 112, 9436–9441, doi:10.1073/pnas.1508686112 (2015).

9. Minskaia, E. et al. Discovery of an RNA virus 3’->5’ exoribonuclease that is critically involved in coronavirus RNA synthesis. Proc Natl Acad Sci U S A 103, 5108–5113, doi:10.1073/pnas.0508200103 (2006).

10. Ferron, F. et al. Structural and molecular basis of mismatch correction and ribavirin excision from coronavirus RNA. Proc Natl Acad Sci U S A 115, E162–E171, doi:10.1073/pnas.1718806115 (2018).

11. Ogando, N. S. et al. The Curious Case of the Nidovirus Exoribonuclease: Its Role in RNA Synthesis and Replication Fidelity. Front Microbiol 10, 1813, doi:10.3389/fmicb.2019.01813 (2019).

12. Eckerle, L. D. et al. Infidelity of SARS-CoV Nsp14-exonuclease mutant virus replication is revealed by complete genome sequencing. PLoS Pathog 6, e1000896, doi:10.1371/journal.ppat.1000896 (2010).

13. Chen, Y. et al. Functional screen reveals SARS coronavirus nonstructural protein nsp14 as a novel cap N7 methyltransferase. Proc Natl Acad Sci U S A 106, 3484–3489, doi:10.1073/pnas.0808790106 (2009).

14. Jin, X. et al. Characterization of the guanine-N7 methyltransferase activity of coronavirus nsp14 on nucleotide GTP. Virus Res 176, 45–52, doi:10.1016/j.virusres.2013.05.001 (2013).

15. Bouvet, M. et al. In vitro reconstitution of SARS-coronavirus mRNA cap methylation. PLoS Pathog 6, e1000863, doi:10.1371/journal.ppat.1000863 (2010).

16. Posthuma, C. C., Te Velthuis, A. J. W. & Snijder, E. J. Nidovirus RNA polymerases: Complex enzymes handling exceptional RNA genomes. Virus Res 234, 58–73, doi:10.1016/j.virusres.2017.01.023 (2017).

17. Hackbart, M., Deng, X. & Baker, S. C. Coronavirus endoribonuclease targets viral polyuridine sequences to evade activating host sensors. Proc Natl Acad Sci U S A 117, 8094–8103, doi:10.1073/pnas.1921485117 (2020).

18. Williams, J. S., Lujan, S. A. & Kunkel, T. A. Processing ribonucleotides incorporated during eukaryotic DNA replication. Nat Rev Mol Cell Biol 17, 350–363, doi:10.1038/nrm.2016.37 (2016).

19. Manners, O., Baquero-Perez, B. & Whitehouse, A. m(6)A: Widespread regulatory control in virus replication. Biochim Biophys Acta Gene Regul Mech 1862, 370–381, doi:10.1016/j.bbagrm.2018.10.015 (2019).

20. Netzband, R. & Pager, C. T. Epitranscriptomic marks: Emerging modulators of RNA virus gene expression. Wiley Interdiscip Rev RNA 11, e1576, doi:10.1002/wrna.1576 (2020).

21. Trott, O. & Olson, A. J. AutoDock Vina: improving the speed and accuracy of docking with a new scoring function, efficient optimization, and multithreading. J Comput Chem 31, 455–461, doi:10.1002/jcc.21334 (2010).

22. Sengerova, B. et al. Characterization of the human SNM1A and SNM1B/Apollo DNA repair exonucleases. J Biol Chem 287, 26254–26267, doi:10.1074/jbc.M112.367243 (2012).

23. Hallick, R. B., Chelm, B. K., Gray, P. W. & Orozco, E. M., Jr. Use of aurintricarboxylic acid as an inhibitor of nucleases during nucleic acid isolation. Nucleic Acids Res 4, 3055–3064, doi:10.1093/nar/4.9.3055 (1977).

24. Chapman, T. M. et al. N-Hydroxyimides and hydroxypyrimidinones as inhibitors of the DNA repair complex ERCC1-XPF. Bioorg Med Chem Lett 25, 4104–4108, doi:10.1016/j.bmcl.2015.08.024 (2015).

25. Tumey, L. N. et al. The identification and optimization of a N-hydroxy urea series of flap endonuclease 1 inhibitors. Bioorg Med Chem Lett 15, 277–281, doi:10.1016/j.bmcl.2004.10.086 (2005).

26. Exell, J. C. et al. Cellularly active N-hydroxyurea FEN1 inhibitors block substrate entry to the active site. Nat Chem Biol 12, 815–821, doi:10.1038/nchembio.2148 (2016).

27. Savarino, A. In-Silico docking of HIV-1 integrase inhibitors reveals a novel drug type acting on an enzyme/DNA reaction intermediate. Retrovirology 4, 21, doi:10.1186/1742-4690-4-21 (2007).

28. Hare, S. et al. Molecular mechanisms of retroviral integrase inhibition and the evolution of viral resistance. Proc Natl Acad Sci U S A 107, 20057–20062, doi:10.1073/pnas.1010246107 (2010).

29. Noguchi, N. Ebselen, a useful tool for understanding cellular redox biology and a promising drug candidate for use in human diseases. Arch Biochem Biophys 595, 109–112, doi:10.1016/j.abb.2015.10.024 (2016).

30. Jin, Z. et al. Structure of M(pro) from SARS-CoV-2 and discovery of its inhibitors. Nature 582, 289–293, doi:10.1038/s41586-020-2223-y (2020).

31. Sauna, Z. E., Shukla, S. & Ambudkar, S. V. Disulfiram, an old drug with new potential therapeutic uses for human cancers and fungal infections. Mol Biosyst 1, 127–134, doi:10.1039/b504392a (2005).

32. Antony, S. & Bayse, C. A. Density functional theory study of the attack of ebselen on a zinc-finger model. Inorg Chem 52, 13803–13805, doi:10.1021/ic401429z (2013).

33. Sekirnik, R. et al. Inhibition of the histone lysine demethylase JMJD2A by ejection of structural Zn(II). Chem Commun (Camb), 6376-6378, doi:10.1039/b916357c (2009).

34. Blessing, H., Kraus, S., Heindl, P., Bal, W. & Hartwig, A. Interaction of selenium compounds with zinc finger proteins involved in DNA repair. Eur J Biochem 271, 3190–3199, doi:10.1111/j.1432-1033.2004.04251.x (2004).

35. Jacob, C., Maret, W. & Vallee, B. L. Ebselen, a selenium-containing redox drug, releases zinc from metallothionein. Biochem Biophys Res Commun 248, 569–573, doi:10.1006/bbrc.1998.9026 (1998).

## REFERENCES

1. Savitsky, P. et al. High-throughput production of human proteins for crystallization: the SGC experience. J Struct Biol 172, 3–13, doi:10.1016/j.jsb.2010.06.008 (2010).

2. Dominy, C. N. & Andrews, D. W. Site-directed mutagenesis by inverse PCR. Methods Mol Biol 235, 209–223, doi:10.1385/1-59259-409-3:209 (2003).

3. Birch, A. J., Salahud-Din, M. & Smith, D. C. C. The synthesis of (±)-xanthorrhoein. Journal of the Chemical Society C: Organic, 523–527, doi:10.1039/J39660000523 (1966).

4. Falsini, M. et al. 3-Hydroxy-1H-quinazoline-2,4-dione as a New Scaffold To Develop Potent and Selective Inhibitors of the Tumor-Associated Carbonic Anhydrases IX and XII. J Med Chem 60, 6428–6439, doi:10.1021/acs.jmedchem.7b00766 (2017).

5. Tang, J. et al. 3-Hydroxypyrimidine-2,4-diones as an inhibitor scaffold of HIV integrase. J Med Chem 54, 2282–2292, doi:10.1021/jm1014378 (2011).

6. Khokhlov, P. S., Osipov, V. N. & Roshchin, A. V. 3-Hydroxy- and 3-alkoxy-2-sulfanylquinazolin-4(3H)-ones: synthesis and reactions with alkylating and acylating agents. Russian Chemical Bulletin 60, 153–156, doi:10.1007/s11172-011-0022-1 (2011).

7. Schneider, C. A., Rasband, W. S. & Eliceiri, K. W. NIH Image to ImageJ: 25 years of image analysis. Nat Methods 9, 671–675, doi:10.1038/nmeth.2089 (2012).

8. Niesen, F. H., Berglund, H. & Vedadi, M. The use of differential scanning fluorimetry to detect ligand interactions that promote protein stability. Nat Protoc 2, 2212–2221, doi:10.1038/nprot.2007.321 (2007).

